# FDR control in GWAS with population structure

**DOI:** 10.1101/2020.08.04.236703

**Authors:** Matteo Sesia, Stephen Bates, Emmanuel Candès, Jonathan Marchini, Chiara Sabatti

**Affiliations:** Department of Data Sciences and Operations, University of Southern California; Departments of Statistics and of EECS, University of California, Berkeley; Departments of Statistics and of Mathematics, Stanford University; Regeneron Genetics Center, Regeneron Pharmaceuticals; Departments of Statistics and of Biomedical Data Sciences, Stanford University

## Abstract

We present a comprehensive statistical framework to analyze data from genome-wide association studies of polygenic traits, producing distinct and interpretable discoveries while controlling the false discovery rate. This approach leverages sophisticated multivariate models, correcting for linkage disequilibrium, and accounts for population structure and relatedness, adapting to the characteristics of the samples at hand. A key element is the recognition that the observed genotypes can be considered as a random sample from an appropriate model, encapsulating our knowledge of genetic inheritance and human populations. This allows us to generate imperfect copies (knockoffs) of these variables which serve as ideal negative controls; knockoffs are indistinguishable from the original genotypes in distribution, and independent from the phenotype. In sharp contrast with state-of-the-art methods, the validity of our inference in no way depends on assumptions about the unknown relation between genotypes and phenotype. We develop and leverage a model for the genotypes that accounts for arbitrary and unknown population structure, which may be due to diverse ancestries or familial relatedness. We build a pipeline that is robust to the most prominent possible confounders, facilitating the discovery of causal variants. Validity and effectiveness are demonstrated by extensive simulations with real data, as well as by the analysis of several phenotypes in the UK Biobank. Finally, fast software is made available for researchers to apply the proposed methodology to Biobank-scale data sets.

## 1 Statistical analysis of GWAS data

Genome-wide association studies shaped the research into the genetic basis of human traits for the past 15 years. While family studies had previously been the cornerstone of genetics, Risch and Merikangas [1] in 1996 described the power that large population samples held for the study of polygenic phenotypes, those influenced by many loci, each with relatively small effect. Ten years of biotech development resulted in the capacity to genotype hundreds of thousands of single nucleotide polymorphisms (SNPs) in thousands of individuals, and in 2007 the first large-scale association studies were published [2]. As of 2021, the NHGRI-EBI GWAS catalog [3] contains over 4800 publications and 240k associations, implicating almost 150k SNPs for a diverse set of traits. The predictions in [1] have been confirmed: there are thousands associations to traits such as high cholesterol or autism—complex traits whose inheritance appeared hard to explain—that are now candidate for follow-up studies. New challenges have thus emerged: how to sort through all these genetic variants? How to identify those that are more likely to be causal? How to select variants that can be used to construct robust prediction scores, maintaining validity across human populations? GWAS were designed based on statistical considerations and a closer look at the inferential methods used today to analyze these data shows progress, many success stories, and some open challenges.

### 1.a The requisites

An effective analysis of GWAS data should (a) account for the role played by multiple genetic variants, and (b) adjust for multiplicity. These studies were intended to uncover the genetic basis of polygenic traits; therefore, the data should be analyzed with statistical models that take into account the effects of all relevant variants. Such multivariate models have two additional benefits. On the one hand, they can explain a higher fraction of phenotypic variance, which benefits prediction—witness the rise of multivariate statistical/machine learning algorithms—and can facilitate the discovery of new loci. On the other hand, they bring us closer to the identification of variants with causal effects; see [4, 5].

Indeed, the main purpose of a GWAS is not to construct a black-box model that predicts phenotypic outcomes given genotype information, but to identify precisely which genetic variants have an impact on the phenotype, uncovering the underlying biological pathway. A meaningful error measure should be based on the number of falsely discovered genetic loci and, as we are exploring the potential effects of hundreds of thousands of variables, a multiplicity adjustment is needed to guarantee the reproducibility of any findings. The false discovery rate (FDR) is a particularly appropriate control target: as we expect to make hundreds or thousands of true discoveries (corresponding to the polygenic nature of the trait), we can certainly tolerate a few false ones [6, 7].

### 1.b The challenges

While striving to achieve the above desiderata, the analysis of GWAS data encountered several obstacles. A first challenge arises from the problem dimensions. The simplest polygenic model [8] describes a phenotype as the result of additive genetic effects, suggesting linear regression of the trait on the genotyped SNPs as an inferential method. However, given that the number of explanatory variables (hundreds of thousands) is larger than the sample size (historically in the thousands, nowadays routinely in the tens of thousands), classical linear regression is not a viable approach. Penalized regression and Bayesian models have been proposed as alternatives [9–11], but they also have shortcomings. The Lasso [12] is geared towards optimizing prediction and does not allow researchers to make statements on error control for the selected variables. While the recent literature on inference after selection [13] attempts to remedy this limitation, the most practical solutions in the context of GWAS are based on sample-splitting [14, 15], with consequent loss of power, and do not guarantee FDR control in finite samples. Bayesian procedures encounter substantial computational complexity; the efficient sampling strategies that have been proposed over the years [**logsdon2010**, 16] often rely on approximations of the posterior distribution that are too crude for precise variable selection.

A second difficulty arises from the strong local dependence between SNPs, known as linkage disequilibrium (LD) [17]. The genotyped SNPs are chosen as to “cover” the genome: even though the true causal variants may not be directly observed, data on sufficiently close SNPs can act as proxy. Genotyping platforms thus rely on many closely spaced SNPs whose alleles are strongly dependent. This design choice poses some challenges for the statistical analysis. In multivariate regression models, dependencies between different variables make it difficult to select any one of them, as others provide very similar signals. Marginal tests for independence between the trait and individual neighboring variants correspond to redundant hypotheses; therefore, such discoveries are more difficult to interpret, and straightforward applications of FDR-controlling methods [18] become particularly problematic [19].

A third challenge is due to the lack of independence across samples, or population structure. In fact, even if the study does not explicitly focus on families, it often includes individuals sharing some degree of relatedness and common ancestry, or otherwise belonging to identifiable subgroups. This problem is accentuated in the large studies carried out today, since even relatively weak spurious associations can become statistically significant if the sample size is sufficiently large.

### 1.c Standard pipeline

The standard statistical pipeline consists of multiple steps, which evolved in response to the above challenges. Typically, one identifies promising signals by testing for the association of the phenotype with one variant at a time, through a simple linear model. To correct for population structure, the SNP-by-SNP marginal regression models may include additional covariates, such as the top principal components of the genotype matrix [20]. Alternatively, random effects may be utilized to account for relatedness and, partially, for the effect of other genetic loci [21–24]. The p-values thus computed are thresholded to approximately control the family wise error rate (FWER). To eliminate the redundancy in the findings, variants significantly associated with the phenotype and highly correlated with one another are “clumped” into distinct groups, utilizing procedures such as that implemented in PLINK [25]. The results of these univariate tests are then taken as input by two different multivariate analyses: fine mapping [26] and polygenic risk scores [27]. The former aims to identify the causal variants among many similarly associated SNPs in LD. The latter seeks to construct a predictor of the trait for future samples based on a large number of variants across the genome.

This pipeline is a patchwork of different approaches, based on strong (e.g., an exact linear model with Gaussian errors) and sometimes contradictory hypotheses (e.g., the number of fixed effects to include in the model). This pipeline is well established, despite the lack of rigorous theoretical grounding, partly because it has led to the discovery of a number of loci which appear to be reproducibly associated with the traits of interest. Indeed, geneticists have identified more statistically significant loci than it is currently practical to investigate in follow-up studies. However, the limitations of the standard approach become more evident when one tries to identify causal variants and leverage them to predict disease risk. On the one hand, despite some interesting fine-mapping methods, there have been few mechanistic validations of loci identified by GWAS [28]: the outputs of the pipeline are hard to interpret directly and expensive to investigate in follow-up studies. On the other hand, the performance of polygenic risk scores is not robust across populations, which highlights the difficulty of identifying causal variants, and raises questions of equity and fairness [29, 30]. A statistical method that pursues more directly the original GWAS requisites may improve performance in both tasks: the discovery of variants or genes that directly influence the phenotype, and the use of such information to estimate individualized disease risk [31, 32].

### 1.d A new framework

We present a statistical approach to GWAS that directly accounts for the role of multiple variants, and adequately adjusts for multiplicity as well as for population structure. This approach, which we call *KnockoffGWAS*, is the culmination of years of developments.

Our work begins with *knockoffs*, introduced in the statistical literature by [33] and later extended by [34]. Knockoffs are randomly generated negative control variables, carefully constructed in such a way as to be indistinguishable from the original *null* variables (those not directly associated with the trait), even with regard to their dependence with the causal ones. Such exchangeability allows researchers to tease apart variants that truly ‘influence’ the trait, since those are the only ones whose association to the phenotype is significantly stronger than that of their knockoffs. This idea is implemented by the *knockoff filter* [33]: the exact algorithm computing a knockoff-based significance threshold guaranteed to control the FDR. Importantly, this filter can be applied to any kind of association statistics (as long as they treat the original variables and the knockoffs fairly), which allows us to exploit the power of modern machine learning algorithms while retaining provably valid inferences [34]. A necessary ingredient for the construction of valid knockoffs is an accurate model for the distribution of the original variables, which we can fortunately obtain in the case of GWAS data. At the same time, knockoffs require no assumptions on the relation between genotypes and phenotype, in stark contrast to the traditional GWAS analysis.

The novelty of this paper lies in a series of technical advances, which cumulatively offer a complete data analysis pipeline accounting for the remaining source of confounding which is of most concern in GWAS, population structure, thus bringing us closer to making proper causal inferences [35]. In particular, we improve on earlier works [36, 37], which focused on knockoffs for individuals from a homogeneous population, by introducing novel methods to handle the possible presence of relatedness, diverse ancestries, and admixture. This allows us to analyze complex data sets in their entirety for the first time, thereby providing inferential results that are both theoretically well grounded and practically relevant.

## 2 Methodology

### 2.a Notation, problem statement and basic assumptions

Consider a data set with genotype and phenotype information for *n* individuals, where *X*^(*i*)^ *∈ {*0, 1, 2*}*^*p*^ counts the minor alleles of the *i*th subject at each of *p* markers, and *Y* ^(*i*)^ *∈𝒴* is the phenotype measurement (which may take either discrete or continuous values in *𝒴*). The *n* individuals are divided into disjoint subsets (either self-reported or inferred), {*F*} _*F ∈ℱ*_, where ℱ is a partition of {1, …, *n*}. We shall refer to these subsets of individuals as *families*, although the grouping is more flexible than in classical family studies.

Our goal is to detect variants (or groups thereof) containing distinct associations with *Y*, as precisely as possible; that is, we ask whether the conditional distribution of *Y* | *X* depends on a variant *X*_*j*_ or a fixed group of variants *X*_*G*_ = *{X*_*j*_*}*_*j∈G*_. For simplicity, we will assume all SNPs in each group *X*_*G*_ to be physically contiguous; see [37] for a thorough justification of this simplification. Let then *𝒢* = *{G}*_*G∈𝒢*_ be any partition of all markers *{*1, …, *p}* into contiguous groups. If *X*_*G*_ denotes the genotypes for all SNPs in group *G*, and *X*_−*G*_ the genotypes for those outside it, we would like to test *conditional* null hypotheses [34] of the form:

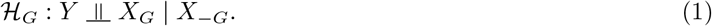

In words, ℋ_*G*_ is true if and only if knowledge of *X*_*G*_ provides no information about *Y* beyond what can be gathered from the knowledge of the rest of the genome.^1^ Testing the null in (1) accounts for population structure because the latter is determined by the genotypes and can be reconstructed almost exactly by looking at *X*_−*G*_. (Since *G* is a relatively small set, *X*_−*G*_ collects almost all measured variants across the genome.) That is, by conditioning on *X*_−*G*_, we are automatically conditioning on ethnicity, subpopulation information, and so on. This is akin to using principal components to capture population structure [20], but *X*_−*G*_ contains even more information: the top principal components can be essentially reconstructed from *X*_−*G*_. This argument is formalized with a causal inference model in Section S1.a and Figure S1, SI. Thus, testing (1) correctly addresses the requisite that the analysis of GWAS data should promote the discovery of interesting biological effects rather than just any association.

Our approach draws strength from modeling the randomness that we already understand, not from making assumptions about the relation between the genetic variants and the trait, which is unknown. In this sense, our method is Fisherian in nature. For instance, we do not posit a (generalized) linear model, although it would be convenient, because we have a priori no way to tell whether it is realistic. Instead, we model the joint distribution of genetic variants because the mechanism by which they are inherited is already well understood. We assume that distinct families are independent of one another but we allow the genotypes of different individuals within the same family to be dependent, encoding the idea that these share close common ancestors. Within each family, the genotypes will be jointly modeled by suitable hidden Markov models (HMMs) that take into account both the observed patterns of close relatedness and the overall ancestry, or ethnic descent, of each individual; this HMM will then be leveraged to generate negative controls, or knockoffs.

### 2.b Exchangeable negative controls

A random matrix 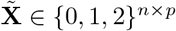 is a knockoff of **X**, with respect to a partition *𝒢* of *{*1, …, *p}*, if it satisfies two properties. First, 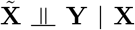, which says that 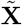 provides no additional information about **Y** (this is always true if 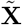 is generated looking at **X** but not at **Y**). Second, the joint distribution of 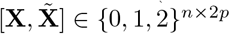 must be invariant upon swapping *X*_*G*_ with the corresponding knockoffs, *simultaneously* for all individuals in any family *F* :

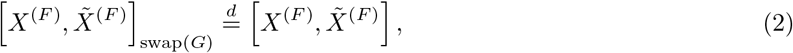

*∀G* ∈ *𝒢, F* ∈ *ℱ*. Above, swap(*G*) swaps all columns of *X*^(*F*)^ indexed by *G* with the corresponding columns of 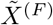, see Figure S2, SI. This means that, upon seeing a list of unordered pairs 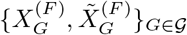, we have no way to tell which genetic variants are original and which are knockoffs. (Of course, looking at *Y* may allow us to tell some variables apart because, conditional on *Y*, the symmetry between *X*_*G*_ and 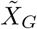 is lost for non-nulls, and this is the whole point of knockoff testing.) The exchangeability property in (2) implies the knockoffs have the same distribution as the genotypes, although it is stronger. For example, permuting the rows of **X** would lead to dummy variables **X′** with the same distribution as **X**, but not in LD with them, and hence not satisfying (2). In fact, any swapping of *X*_*G*_ with 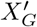 would be noticeable if we compare them to the real variants from neighboring groups. By contrast, our knockoffs 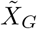 will be in LD with 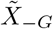 as well as with *X*_−*G*_, thanks to Monte Carlo sampling techniques for HMMs developed in [36, 37]; see Methods. Importantly, in order to satisfy (2), our knockoffs do not simply preserve short-range LD but they are also consistent with the ancestries and family structures reconstructed from all remaining variants. This is non-trivial, especially if the population structure is unknown a priori, and requires a novel approach; see Section 2.c for the key ideas, while the details are in Methods and Sections S1.b–S1.c, SI.

Knockoffs for a partition 𝒢 are designed to test ℋ _*G*_ in (S3). Thus, the partition controls power and resolution in our hypothesis testing problem. Smaller groups of SNPs allow us to test more informative but fundamentally more challenging hypotheses. As a result, higher-resolution knockoffs must satisfy stronger exchangeability in (2) and tend to be individually more similar to the real genotypes (Figure S3, SI), which reduces power [37]. Larger groups of SNPs relax the constraint in (2), which increases power but makes any discoveries less informative. In practice, we will consider different genome partitions, ranging from that in which each group contains a single typed SNP to partitions in which each group spans a few hundreds of kilo-bases.

We demonstrate empirically we can construct knockoffs that are nearly indistinguishable, in the above sense, from the real genotypes in the UK Biobank data. Figure 1 visualizes different measures of exchangeability comparing our knockoffs to real genotypes in terms of population structure, familial relatedness, and LD. For simplicity, we focus on the simplest partition containing exactly one SNP per group, which must follow the strictest exchangeability requirements (analogous diagnostics for low-resolution knockoffs are in Figure S4, SI). Figure 1 (a) shows that a principal component analysis on 10,000 individuals with extremely diverse ancestries (Table S1, SI) gives very similar results when performed on either **X** or 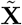; this is consistent with the requirement that knockoffs preserve population structure; see Section S1.a, SI, for more details about knockoff exchangeability and population structure. By contrast, knockoffs based on the fastPHASE HMM [39] were not guaranteed to preserve population structure [36, 37], and indeed they do not in this case; see Figure S5. Figure 1 (b) demonstrates the estimated kinship of any two related individuals is the same regardless of whether it is based on **X** or 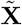. Figure 1 (c) shows knockoffs also preserve LD. The scatter plots reveal the absolute correlation coefficients between genotypes on pairs of nearby variants (within 100 kb of each other) are unchanged when either one or both variables are replaced by their knockoffs. Furthermore, our knockoffs also preserve long-range LD within the same chromosome (Figures S6–S8).

**Figure 1:**
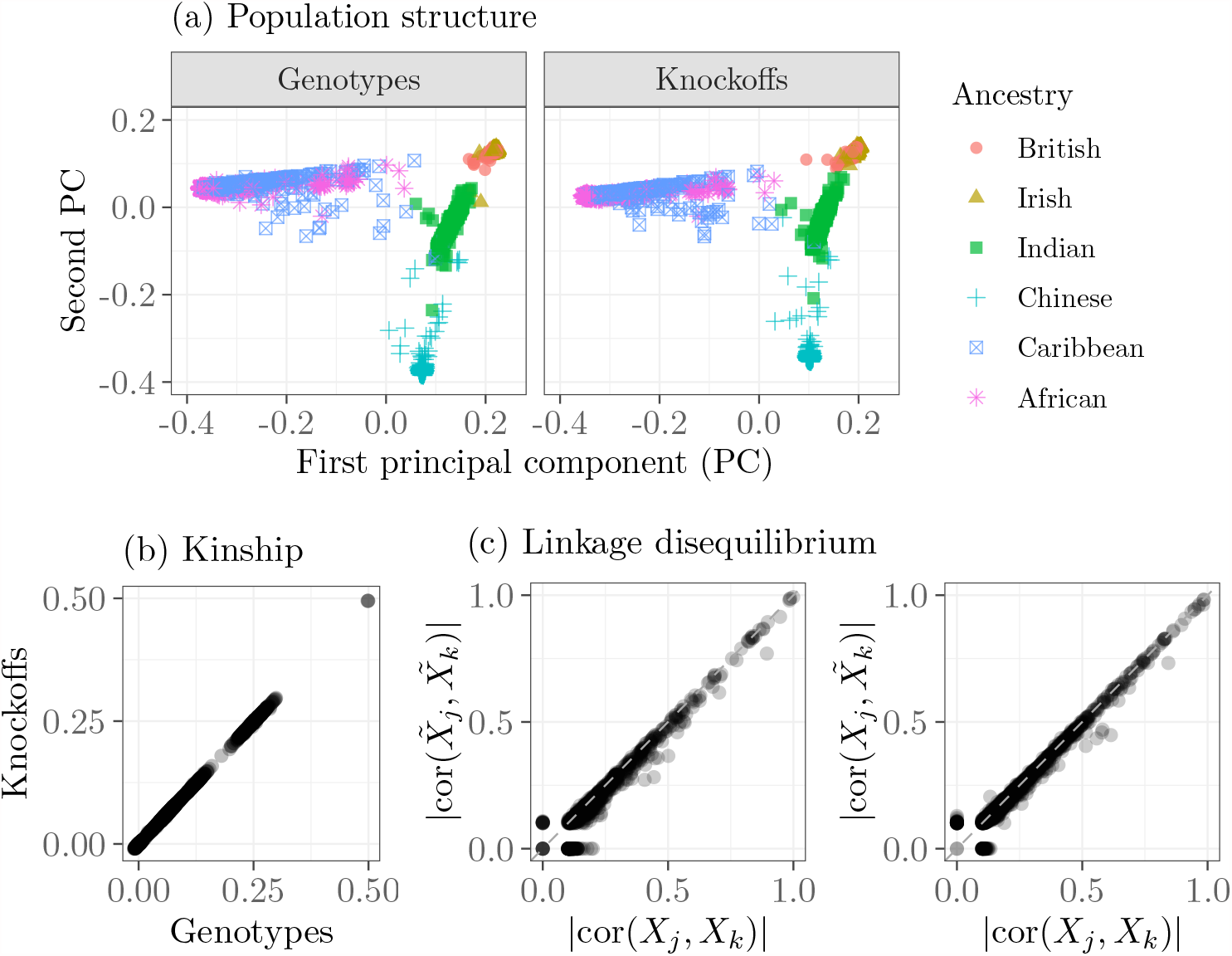
Exchangeability of knockoffs and real UK Biobank genotypes. (a) Principal component analysis for 10k individuals with diverse ancestries, separately for genotypes and knockoffs. (b) Kinship coefficients between 2000 pairs of related individuals, computed separately on genotypes and knockoffs. Kinship is measured by means of KING coefficients [40] so that a value of 0.5 indicates monozygotic twins and 0 indicates no relatedness. (c) Pairwise correlations between nearby variants on chromosome 22 (minor allele frequency *≥*0.01) for the same individuals as in (a), with (left) or without (right) swapping genotypes (*X*) and knockoffs 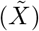.

### 2.c Modeling genotypes and constructing knockoffs

We explain here the high-level ideas of our knockoff construction, deferring the details to the Methods section. First, we assume all haplotypes have been phased and we denote by *H*^(*i,a*)^, *H*^(*i,b*)^ ∈ {0, 1}^*p*^ those inherited by individual *i* from each parent, so that *X*^(*i*)^ = *H*^(*i,a*)^ + *H*^(*i,b*)^. As in [37], we model the haplotypes and leverage them to construct phased knockoffs, namely 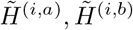, which can then be simply combined to obtain valid knockoff genotypes: 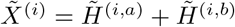.

The haplotype distribution is approximated by an HMM similar to that of SHAPEIT [41–43]. This overcomes the main limitation of the fastPHASE HMM [39], which cannot simultaneously describe LD and population structure [44], but which we previously used to construct knockoff genotypes. Thanks to this new modeling choice, we can use knockoffs for the analysis of genetic data beyond homogeneous populations [37]. The SHAPEIT HMM describes each haplotype sequence as a mosaic of *K* reference motifs corresponding to the haplotypes of other individuals in the data set, where *K* is fixed (e.g., *K* = 100). Critically, different haplotypes may have different sets of motifs, chosen based on their similarity; see Methods for details. The idea is that these references reflect the ancestry of an individual (our variable *A*); in fact, the haplotypes of someone from England should be well approximated by a mosaic of haplotypes primarily belonging to other English individuals. Conditional on the references, the identity of the motif copied at each position is described by a Markov chain with transition probabilities proportional to the genetic distances between neighboring sites; different chromosomes are treated as independent. Conditional on the Markov chain, the motifs are copied imperfectly, as relatively rare mutations can independently occur at any site. After inferring the unobserved Markov chain in this HMM, we carefully perturb it to construct knockoffs, while preserving the population structure. Figure 2 (a) presents a schematic visualization of this method.

**Figure 2:**
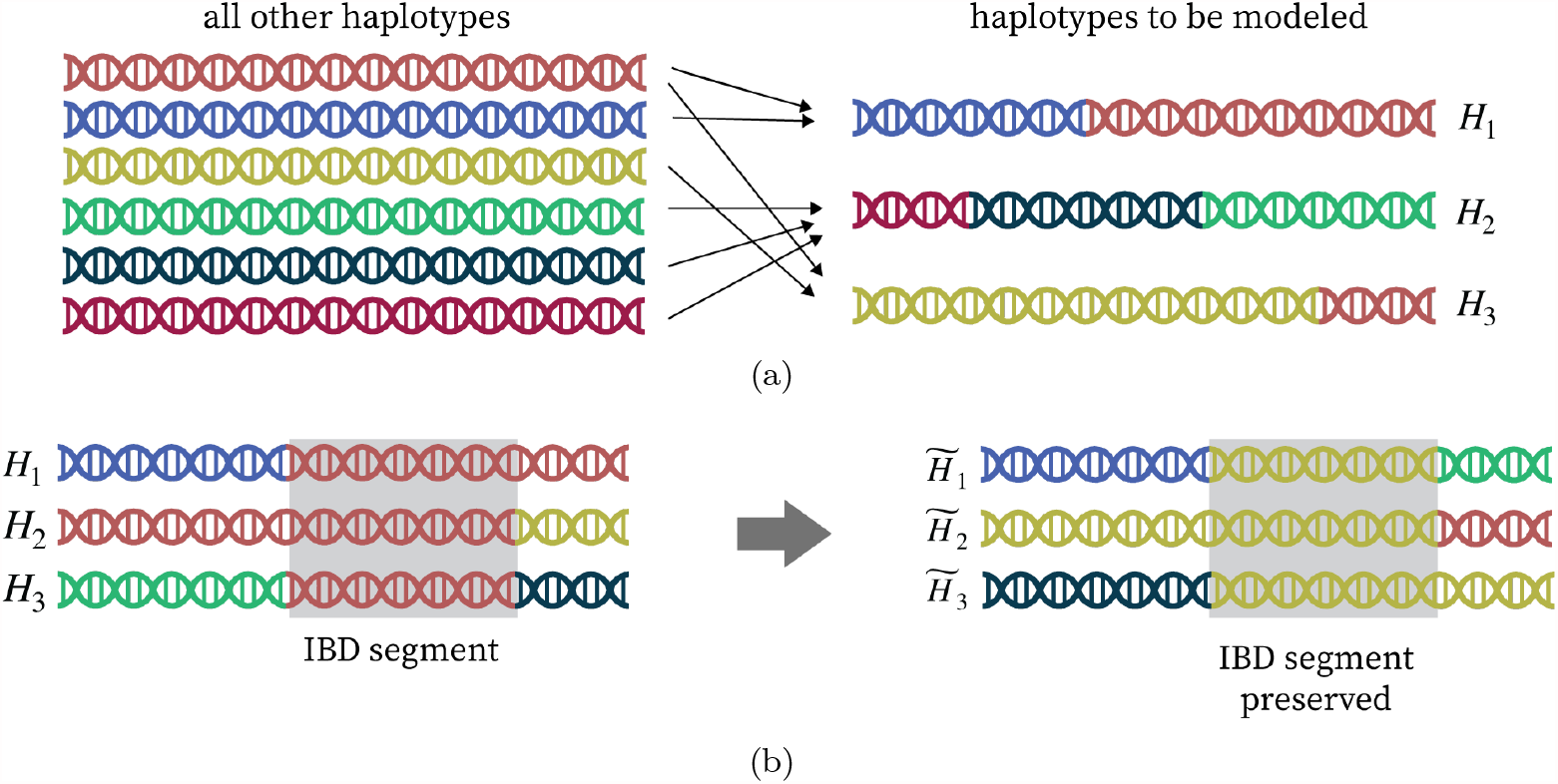
Visualization of the haplotype HMM and knockoff construction. (a) Each haplotype sequence, *H*^(*i*)^, is described as a mosaic of motifs from a subset of other haplotypes (in different colors). (b) Haplotypes and knockoffs of close relatives share IBD segments where their alleles match exactly (shaded segments).

The above model does not account for close relatedness because it describes all haplotypes as conditionally independent given the reference motifs, which cannot explain the presence of long IBD segments within families [45, 46]. To handle this, we will model jointly haplotypes in the same family. First, we detect long IBD segments in the data [47–50]. If the pedigrees are known a priori, the IBD search can be focused within the given families; otherwise, there exists software to approximately reconstruct families from data at the UK Biobank scale [51]. The results define a relatedness graph in which two haplotypes are connected if they share an IBD segment; we refer to the connected components of this graph as the *IBD-sharing families*. Second, we define a larger HMM jointly characterizing the distribution of all haplotypes in each IBD-sharing family, conditional on the location of the segments. Marginally, each haplotype is modeled by the SHAPEIT HMM; however, the Markov chains for related individuals are forced to match along the IBD segments. This coupling will be preserved in the knockoffs, in the sense that the latter will contain exchangeable IBD segments in the same locations, although the knockoffs may not necessarily have the same alleles as the original haplotypes; see Figure 2 (b) and Figure S9.

### 2.d The knockoff filter

Although knockoffs are constructed to be statistically indistinguishable from the genotypes, it may be possible to tell them apart by looking also at the phenotype, since 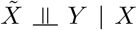 but 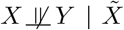. Loosely speaking, this implies differences in the comparisons of *Y* with either *X*_*G*_ or 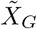 can provide evidence against the null hypothesis *X*_*G*_ *╨Y* | *X*_−*G*_. Such property is leveraged by estimating importance measures, *T*_*G*_ and 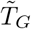, for each group of SNPs and knockoffs, respectively. The importance measures are combined into a test statistic for each group *G* ∈ *𝒢*, i.e., 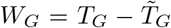. This is designed such that a large value of *W*_*G*_ *>* 0 is evidence against the null hypothesis, while the sign of statistics for null groups are independent coin flips [33]. The knockoff filter computes an adaptive significance threshold for these test statistics, which provably controls the FDR as long as the knockoffs are correctly exchangeable. See Figure S10, SI, for a schematic of the entire procedure. In addition to controlling the FDR, we can assess the significance of individual findings, either through a local estimate of the false discovery proportion (FDP) [37, 52] (which requires a sufficiently large number of discoveries), or through a q-value [53], which is defined as the smallest nominal FDR level at which that discovery could have been reported.

The FDR guarantee holds regardless of the unknown relation between *X* and *Y*, and with any importance statistics. The statistics can be computed by virtually any method and easily incorporate prior knowledge [34]. As in earlier works [34, 36, 37], we utilize a sparse generalized linear model (Lasso) [12], although our inference never assume its validity. Such an approach is computationally feasible with large data [54], interpretable [4], and powerful compared to state-of-the-art LMMs. Concretely, we fit a sparse regression of *Y* on 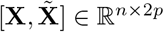, after standardizing the columns to have unit variance. The importance measures are 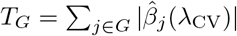 and 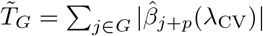, where 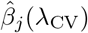 (resp. 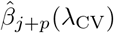) is the Lasso coefficient for *X*_*j*_ (resp. 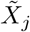) at a value of the regularization parameter tuned by cross-validation to achieve a low prediction error.

To localize causal variants as precisely as possible, we apply the knockoff filter at multiple resolutions; partitions with larger groups yield more power, but at the cost of less informative findings. If the knockoffs are valid, the FDR is controlled at each level of resolution. Further, it is possible to coordinate the results at different resolutions to ensure they are consistent with one another [37]. This requires a variation of the knockoff filter first proposed in [38], which we do not apply here in the interest of simplicity.

### 2.e Imputed variants

Imputed variants—those not directly measured but instead predicted with a model based on nearby observed variants—are commonly included in fine-mapping analyses [55]. However, correctly utilizing imputed variants requires care, since, of course, they are not as informative as the measured variants. In particular, imputed variants are conditionally independent of the trait given the observed genotypes (they are constructed based only on the latter). As such, it is impossible to attribute distinct signals to imputed variants without additional modeling assumptions; by definition, any association observed between them and the trait could be equally well explained as a (possibly nonlinear) association with the observed variants. In principle, two forms of modeling assumptions could be used to help attribute signal to an imputed variant. First, functional annotations could lead one to believe that the imputed variant is more plausibly responsible for an observed association. Second, one may posit a sparse (generalized) linear model and search for variants that individually explain as much of the response as possible. The latter is the most typical approach [56].

We do not include imputed variants in our analysis due to the fundamental issue above: one can never conclude that an unseen variant is causal based on the data alone. Our method identifies relations supported by the available evidence without the need of modeling assumptions, such as sparsity and linearity, and does not make further claims. In particular, we never single out an imputed variant as causal, but instead we identify promising regions of the genome containing at least one measured variant. This is an impartial reporting of the evidence at hand, isolating statistical associations only to the resolution achievable with the data at hand. While it may sometimes be desirable to conduct fine-mapping at the resolution of imputed variants, we keep this task distinct from our analysis because the model-based attribution of signal to imputed variants is a fundamentally different statistical problem requiring other tools and assumptions. In this sense, unmeasured variants would remain a possible source of confounding for our method (as illustrated in Figure S1) regardless of whether imputed SNPs are included in the analysis, although such confounding can be expected to be almost negligible when interpreting our findings at low resolution (e.g., hundreds of kb).

### 2.f Leveraging covariates

The flexibility of our method allows one to easily leverage additional information to compute even more powerful statistics. In our application, we include relevant measured covariates *U* ^m^ (such as sex, age and squared age, as well the the top 5 genetic principal components) in the above regression model, replacing 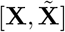 with 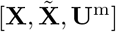. These covariates explain some of the phenotypic variation that cannot be captured by a sparse linear combination of genotypes, thus reducing the noise in the model. The coefficients for *U* ^m^ are not regularized, and they do not directly enter the calculated statistics. It must be emphasized that the validity of the knockoff filter requires the knockoffs to be exchangeable with the genotypes *conditional* on any covariates utilized in the computation of the test statistics, see (2). This is the case in our analysis, since sex and age can be safely assumed to be independent of the genotypes (we do not analyze the sex chromosomes), and the principal components are related to the genotypes through the population structure, *A*, for which our knockoffs already account. In general, one can thus explicitly analyze any subset *U* ^m^ of the covariates *U* that are already implicitly taken into account in (1).

## 3 Application to the UK Biobank data

### 3.a Pre-processing of the UK Biobank data

We test our method through simulations based on the phased haplotypes of 489k UK Biobank individuals (592k SNPs, chromosomes 1–22). After some pre-processing (Methods), we partition each autosome into contiguous groups at 7 levels of resolution, ranging from that of single SNPs to that of 425 kb-wide groups; see Table S2. The partitions are obtained from complete-linkage hierarchical clustering, cutting the dendrogram at different heights. The similarity measures are based on genetic distances computed in the European population, which serve as a simple proxy for LD and thus lead to relatively homogeneous groups of SNPs. The individuals have diverse ancestries, although most are British (430k), Irish (13k), or other Europeans (16k). There are 136,818 samples with reported close relatives, divided in 57,164 families. We apply RaPID [51] to detect IBD segments longer than 3 cM, chromosome-by-chromosome, ignoring (for simplicity) those shared by individuals who are not in the same self-reported family; this gives us 7,087,643 distinct segments over the entire genome. Then, we generate knockoffs preserving both population structure and IBD segments, as previewed in Figure 1.

### 3.b Analysis of simulated phenotypes

We simulate continuous phenotypes conditional on the real genotypes, according to a homoscedastic linear model with 4000 causal variants paired in 2000 (100 kb-wide) loci placed uniformly across the genome, so that each locus contains 2 causal variants. The total heritability is varied as a control parameter. This gives us a controlled but realistic testing environment. We take BOLT-LMM [24] as a benchmark, applying it on the same data with standard parameters. However, the comparison requires some care since LMMs are designed to test marginal associations, accounting for population structure [24] but not LD [37], and controlling the FWER instead of the FDR.

It is standard to combine (or clump) marginally significant LMM discoveries from nearby SNPs, e.g., using the standard PLINK [25] algorithm as in [24]. Ideally, this should allow each clump to be interpreted as indicating a distinct discovery; however, the solution is imperfect because even relatively far-apart SNPs may not be completely independent of each other if they are on the same chromosome. This issue is particularly important at the Biobank scale, as larger sample sizes make even weak correlations statistically significant, complicating the interpretation of marginal hypotheses. We address this difficulty by further consolidating the clumps computed by PLINK if they are within 100 kb of each other, as in [37]. In our simulations, this strategy ensures the final discoveries are distinct because the true causal loci are well spaced, but that may not always be the case in practice. Therefore, it is unclear how to best clump marginal associations in general.

Figure 3 visualizes our discoveries within a locus containing two causal variants, although the method operates genome-wide; see Figure S11. Figure 3 also shows the nearby LMM findings, clumped by PLINK but unconsolidated. Here, our method localizes the causal variants precisely, while the LMM findings span a long region including many spurious associations. Figure S12 describes how the results obtained with each methods change as the signal strength is varied.

**Figure 3:**
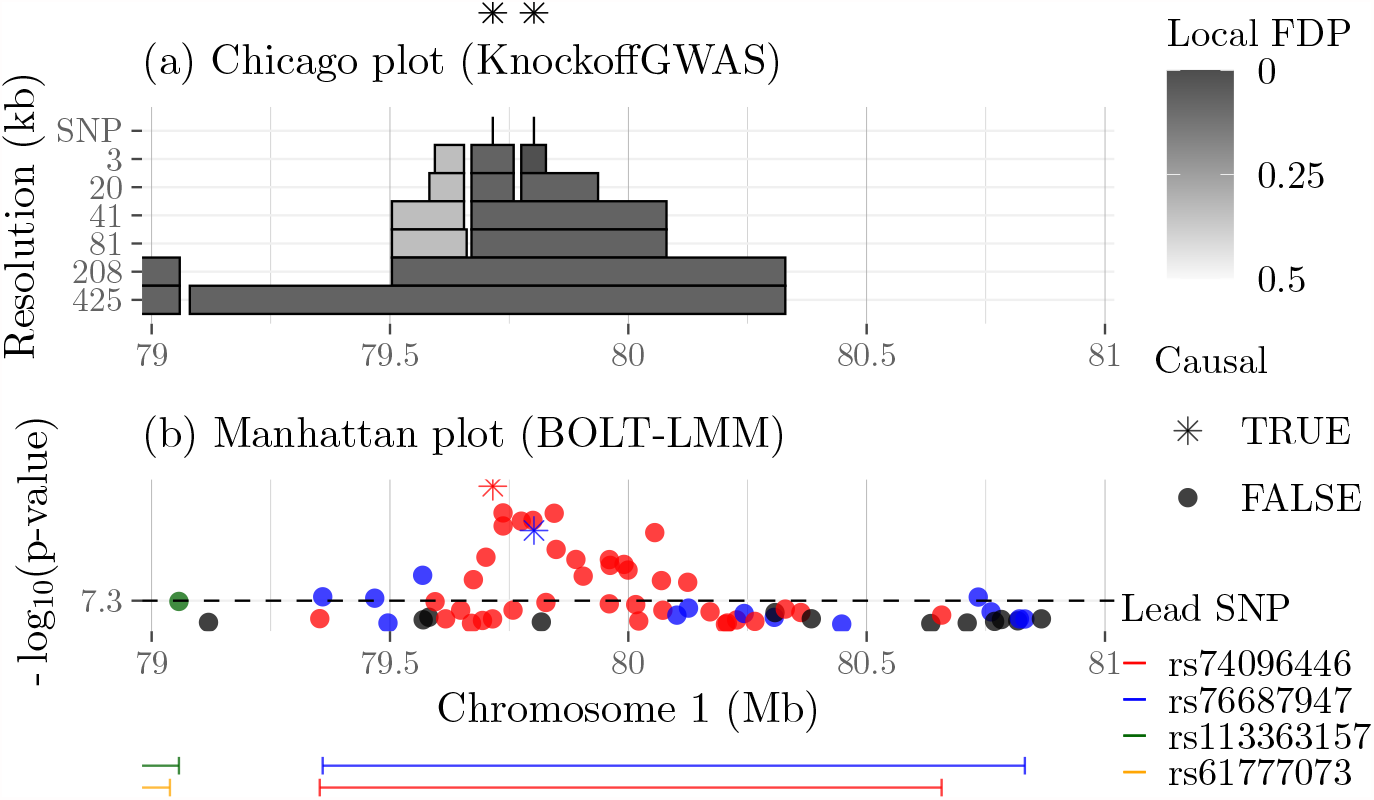
KnockoffGWAS and BOLT-LMM discoveries for a simulated trait based on the genotypes of 489k UK Biobank individuals with population structure. (a) The shaded rectangles represent our discoveries at different resolutions. The FDR level is 10%. Darker rectangles have lower estimated local FDP. The light rectangles are false discoveries because they do not contain causal variants (whose positions are marked by asterisks on top). (b) BOLT-LMM p-values from the same data, and PLINK clumps (segments below) at the genome-wide significance level (5 × 10^−8^), utilizing different colors for different clumps. The colors match those of the corresponding p-values.

Regarding the discrepancy in the target error rates, it is unfortunately difficult to use LMMs to control the FDR. The issue is this: LMMs test marginal hypotheses (and the test statistics are not independent of each other), while we ultimately need to report distinct (conditional) discoveries. This issue was discussed in [37], and it is consistent with the general difficulty of controlling the FDR after applying any sort of post-processing to the output of a testing procedure [57]. We consider two possible solutions to make the error rates more comparable, which are informative within our simulations but would not work in practice. The naive approach is to apply the Benjamini-Hochberg (BH) correction [18] to the marginal p-values before clumping. We shall see this inflates the type-I errors. The second approach involves an imaginary oracle which knows the identities of the causal variants and exploits them to automatically adjusts the significance level for the LMM p-values, such that the proportion of false discoveries (after clumping) equals the target FDR exactly. Obviously, this oracle does not exist for real phenotypes.

Figure 4 compares the performances of KnockoffGWAS and BOLT-LMM. The latter targets the FWER (5 × 10^−8^ genome-wide significance level), or the FDR with either of the two aforementioned strategies. Performance is assessed in terms of power, false discoveries, and resolution. Figure 4 (a) focuses on KnockoffGWAS at low resolution (genome partition with median group size equal to 208 kb) and BOLT-LMM with strong clumping (consolidating clumps within 100 kb of each other). Here, power is measured as the proportion of the 2000 causal loci encompassed by at least one discovery, while the FDP is defined as the fraction of findings not containing causal variants. The resolution is measured as the median width of the reported segments. The results show our method is almost as powerful as the imaginary LMM oracle, but is slightly more conservative and reports narrower (more informative) discoveries. By contrast, BOLT-LMM makes fewer discoveries when targeting the FWER, and reports too many false positives when heuristically targeting the FDR.

**Figure 4:**
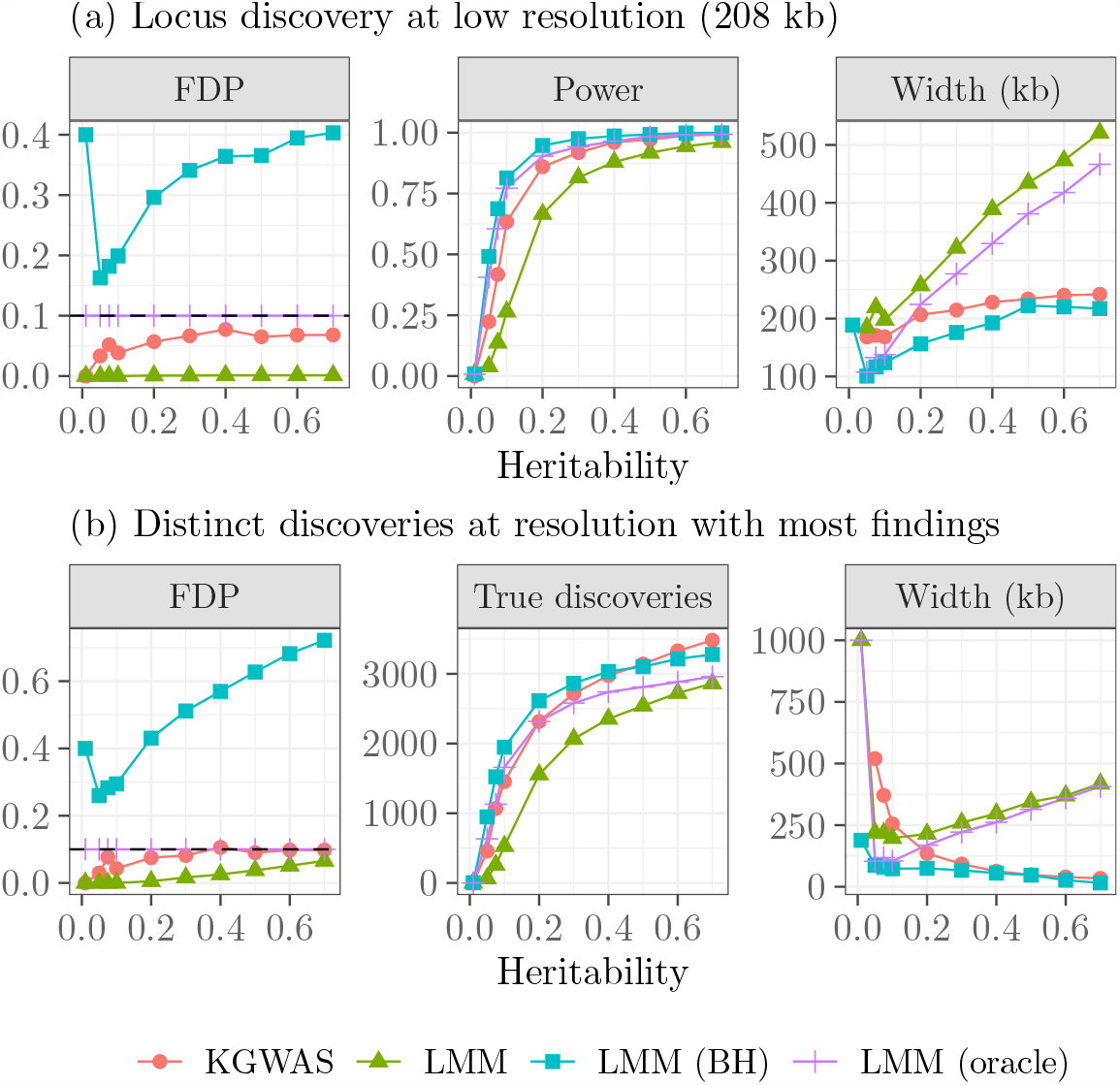
Performance on real genotypes and synthetic phenotypes from a model with 4000 causal variants. The results obtained with either KnockoffGWAS (nominal FDR 0.1) or BOLT-LMM (5 × 10^−8^, heuristic FDR, and oracle FDP calibration) are shown as a function of the total heritability. (a) Low-resolution KnockoffGWAS discoveries and strongly clumped LMM findings. (b) Multi-resolution KnockoffGWAS discoveries (reported only at the resolution with the most findings for each value of heritability) and weakly clumped BOLT-LMM findings. Other details are as in Figure 3.

Figure 4 (b) summarizes the KnockoffGWAS discoveries at different resolutions, reporting only the resolution with the most findings, for simplicity. Since the reported resolution may be different for different heritability values, we measure power in terms of the total number of true discoveries rather than as a fraction. Here, the goal is to detect as many distinct associations as possible, ideally also distinguishing between multiple causal variants in the same locus, so the LMM findings are clumped but unconsolidated. KnockoffGWAS always controls the proportion of false discoveries below the target FDR, and it is either comparable to or more powerful than the oracle. Furthermore, we localize causal variants more precisely as the heritability increases, while the LMM clumps become wider and increasingly polluted by spurious associations, as visualized in Figure S12. Additional simulations on subsets of the UK Biobank samples (Section S2.a, SI) demonstrate our method is robust and powerful even when the individuals have extremely diverse ancestries (Table S1, Figures S13–S14), or very strong familial relatedness (Table S3, Figure S15), in which cases the FDR would have been much larger than desired had we not taken the population structure into account.

### 3.c Analysis of UK Biobank phenotypes

We study 4 continuous traits (height, body mass index, platelet count, systolic blood pressure) and 4 diseases (cardiovascular disease, respiratory disease, hyperthyroidism, diabetes), as defined in Table S4. To increase power, we include a few relevant covariates, as explained in Section 2.d. Table 1 reports the numbers of low-resolution (208 kb) discoveries (target FDR 10%) and compares them to those obtained by BOLT-LMM, which are clumped with PLINK (5 × 10^−8^ significance threshold) but unconsolidated, consistent with [24]. BOLT-LMM is applied on 459k European samples [24] for all phenotypes except diabetes and respiratory disease, for which it is applied on 350k unrelated British samples [37] for the sake of consistency in phenotype definitions. These results suggest KnockoffGWAS is much more powerful; in fact, we discover almost all findings reported by the LMM and many new ones. The findings at other resolutions are summarized in Table S5. Note that the model assumed by BOLT-LMM assumes continuous-valued phenotypes, and other LMM-based methods have been specifically developed for case-control studies [58]; however, BOLT-LMM is still a standard benchmark here because the ratio of cases and controls is not very small [24]; see Table S4. Figure S16 visualizes our discoveries for cardiovascular disease in the form of Manhattan plots, using q-values [53] to measure the individual significance of each finding. The full list of our discoveries is available online at https://msesia.github.io/knockoffgwas/, along with an interactive visualization tool.

**Table 1:** KnockoffGWAS discoveries (208 kb resolution, 10% FDR) using all 487k UK Biobank samples, and corresponding BOLT-LMM findings (5 × 10^−8^). For example, we report 940 distinct discoveries for cardiovascular disease, 274 of which contain significant LMM associations. The LMM reports 257 discoveries for this phenotype, 96.9% of which overlap with at least one of our discoveries.

| Phenotype | KnockoffGWAS discoveries |  | BOLT-LMM discoveries |  |
| --- | --- | --- | --- | --- |
|  | Total | Overlap with LMM | Total | Overlap with KZ |
| bmi | 2395 | 898 (37.5%) | 697 | 689 (98.9%) |
| cvd | 940 | 274 (29.1%) | 257 | 249 (96.9%) |
| diabetes | 113 | 52 (46.0%) | 62 | 55 (88.7%) |
| height | 3339 | 2228 (66.7%) | 2464 | 2430 (98.6%) |
| hypothyroidism | 295 | 129 (43.7%) | 143 | 142 (99.3%) |
| platelet | 1743 | 1057 (60.6%) | 1204 | 1183 (98.3%) |
| respiratory | 262 | 82 (31.3%) | 94 | 92 (97.9%) |
| sbp | 1183 | 561 (47.4%) | 568 | 530 (93.3%) |

Table S6 confirms our findings are consistent with [37], although our method is now even more powerful because it leverages a larger sample—see Table S7 for a more detailed summary. The only exception is at the single-SNP resolution, which may be partially explained by noting these discoveries are fewer and thus more susceptible to the random variability of knockoffs. Table 2 summarizes the increase in discoveries at each resolution directly resulting from the inclusion of samples with close relatives or non-British ancestry. The inclusion of related individuals yields many more discoveries, while it is unsurprising that diverse ancestries bring smaller gains since there are relatively few non-British samples.

**Table 2:** Cumulative numbers of discoveries for all UK Biobank phenotypes at different resolutions, utilizing different subsets of the samples. For example, including related individuals increases by 16.4% the number of discoveries obtained from the British samples at the 208 kb resolution (from 8635 to 10049). As another example, adding non-British individuals (including related ones) increases by 2.2% the number of discoveries obtained from the British samples (including related ones) at the 208 kb resolution (from 10049 to 10270).

| Resolution | Including related samples |  |  |  | Including non-British samples |  |  |  |
| --- | --- | --- | --- | --- | --- | --- | --- | --- |
|  | Everyone |  | British |  | Related |  | Unrelated |  |
|  | Total | Change (%) | Total | Change (%) | Total | Change (%) | Total | Change (%) |
| single-SNP | 138–167 | 21.0 | 155–125 | -19.4 | 125–167 | 33.6 | 155–138 | -11.0 |
| 3 kb | 921–992 | 7.7 | 655–971 | 48.2 | 971–992 | 2.2 | 655–921 | 40.6 |
| 20 kb | 2814–3527 | 25.3 | 2808–3355 | 19.5 | 3355–3527 | 5.1 | 2808–2814 | 0.2 |
| 41 kb | 4419–5867 | 32.8 | 4353–5354 | 23.0 | 5354–5867 | 9.6 | 4353–4419 | 1.5 |
| 81 kb | 6784–8031 | 18.4 | 6676–7781 | 16.6 | 7781–8031 | 3.2 | 6676–6784 | 1.6 |
| 208 kb | 8776–10270 | 17.0 | 8635–10049 | 16.4 | 10049–10270 | 2.2 | 8635–8776 | 1.6 |
| 425 kb | 9401–10730 | 14.1 | 9028–10297 | 14.1 | 10297–10730 | 4.2 | 9028–9401 | 4.1 |
| Sample size | 408k–487k | 19.4 | 356k–430k | 20.8 | 430k–487k | 13.0 | 356k–408k | 14.6 |

### 3.d Validation of novel discoveries

We begin to validate our findings by comparing them to those in the GWAS Catalog [59], in the Japan Biobank Project [60], and in the FinnGen resource [61] (we use the standard 5 × 10^−8^ threshold for the p-values reported by the latter two). Table S8 indicates most high-resolution discoveries correspond to SNPs with a known association to the phenotype of interest. This is especially true for findings also detected by BOLT-LMM, although many of our new ones are confirmed; see Table S9. For example, we report 1089 discoveries for cardiovascular disease at the 425 kb resolution, only 255 of which are detected by BOLT-LMM; however, 85.6% of our additional 834 discoveries are confirmed in at least one of the aforementioned resources. Furthermore, Table S10 suggests most relevant associations in the Catalog (above 70%) are confirmed by our findings, which is again indicative of high power. The relative power (i.e., the proportion of known associations that we discover) seems to be above 90% for quantitative traits, but lower than 50% for all diseases except hypothyroidism, probably due to the relatively small number of cases in the UK Biobank data set compared to more targeted case-control studies.

The 5 × 10^−8^ threshold for the Japan Biobank Project and the FinnGen resource is overly conservative if the goal is to confirm selected discoveries. Therefore, we next carry out an enrichment analysis. The idea is to compare the distribution of the external statistics within our selected loci to those from the rest of the genome; see Note S2.c, SI. This approach estimates the number of replicated discoveries but cannot tell exactly which are confirmed; therefore, we will consider alternative validations later. (A more precise analysis is possible in theory, but has low power; see Note S2.c), SI. Table S11 shows many additional discoveries can thus be validated, especially at high resolution. (See Table S12 for more details about enrichment.) Table 3 summarizes the confirmatory results. Respiratory disease is excluded here because the FinnGen resource divides it among several fields, so it is unclear how to best obtain a single p-value. Regardless, the GWAS Catalog and the FinnGen resource already directly validate 90% of those findings.

**Table 3:**
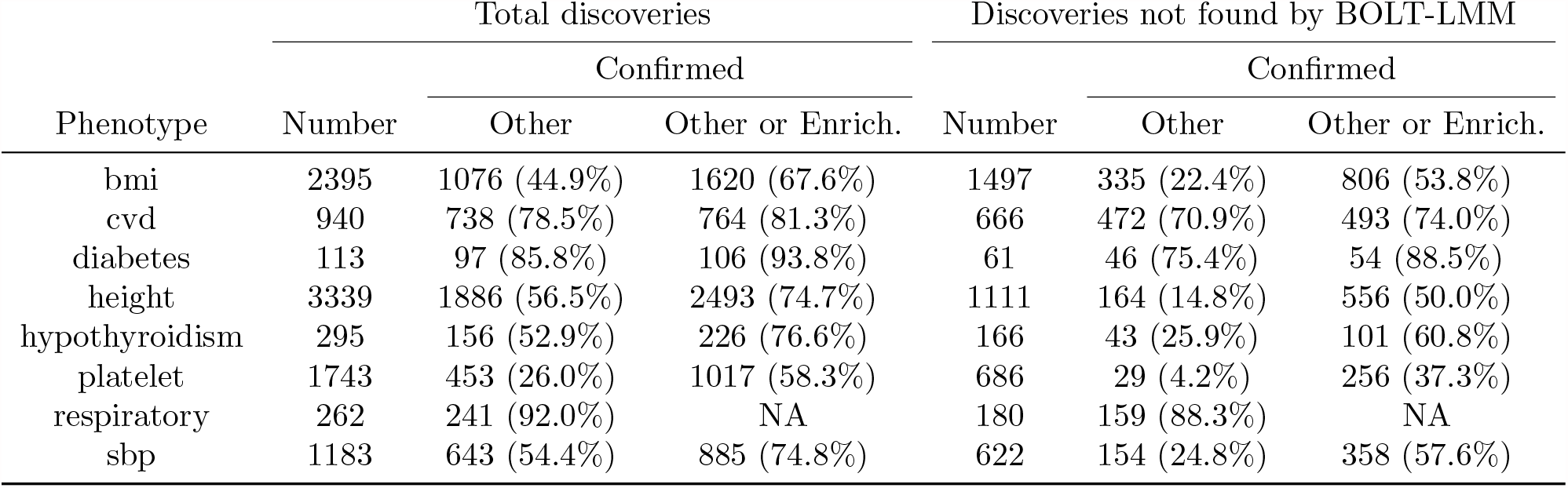
Numbers of low-resolution (208 kb) discoveries obtained with our method and confirmed by other studies, or by an enrichment analysis carried out on external summary statistics. For example, 81.3% of our 940 discoveries for cardiovascular disease are confirmed either by the results of other studies, or by the enrichment analysis. The results are stratified based on whether our findings can be detected by BOLT-LMM using the UK Biobank data (excluding non-European individuals).

| Phenotype | Total discoveries |  |  | Discoveries not found by BOLT-LMM |  |  |
| --- | --- | --- | --- | --- | --- | --- |
|  | Number | Confirmed |  | Number | Confirmed |  |
|  |  | Other | Other or Enrich. |  | Other | Other or Enrich. |
| bmi | 2395 | 1076 (44.9%) | 1620 (67.6%) | 1497 | 335 (22.4%) | 806 (53.8%) |
| cvd | 940 | 738 (78.5%) | 764 (81.3%) | 666 | 472 (70.9%) | 493 (74.0%) |
| diabetes | 113 | 97 (85.8%) | 106 (93.8%) | 61 | 46 (75.4%) | 54 (88.5%) |
| height | 3339 | 1886 (56.5%) | 2493 (74.7%) | 1111 | 164 (14.8%) | 556 (50.0%) |
| hypothyroidism | 295 | 156 (52.9%) | 226 (76.6%) | 166 | 43 (25.9%) | 101 (60.8%) |
| platelet | 1743 | 453 (26.0%) | 1017 (58.3%) | 686 | 29 (4.2%) | 256 (37.3%) |
| respiratory | 262 | 241 (92.0%) | NA | 180 | 159 (88.3%) | NA |
| sbp | 1183 | 643 (54.4%) | 885 (74.8%) | 622 | 154 (24.8%) | 358 (57.6%) |

We continue by cross-referencing with the genetics literature the novel discoveries missed by BOLT-LMM and unconfirmed by the above studies, focusing for simplicity on the 20 kb resolution. Table S13 shows most of these discoveries point to SNPs with known associations to phenotypes closely related to that of interest; see Table S14 for details. Finally, Table S15 shows that many lead SNPs (those with the largest importance measure in each group) within our unconfirmed discoveries have known consequences in the protein-coding sequence.

Figure 5 showcases a novel discovery for cardiovascular disease, which seems unlikely to be false based on the estimated local FDP. The highest resolution finding here spans 4 genes, but we could not find previously reported associations with cardiovascular disease within this locus. However, one gene (SH3TC2) is known to be associated with blood pressure [62], while another (ABLIM3) is associated with body mass index [63]. Figure S17 visualizes the same discovery in the context of a much wider portion of the chromosome.

**Figure 5:**
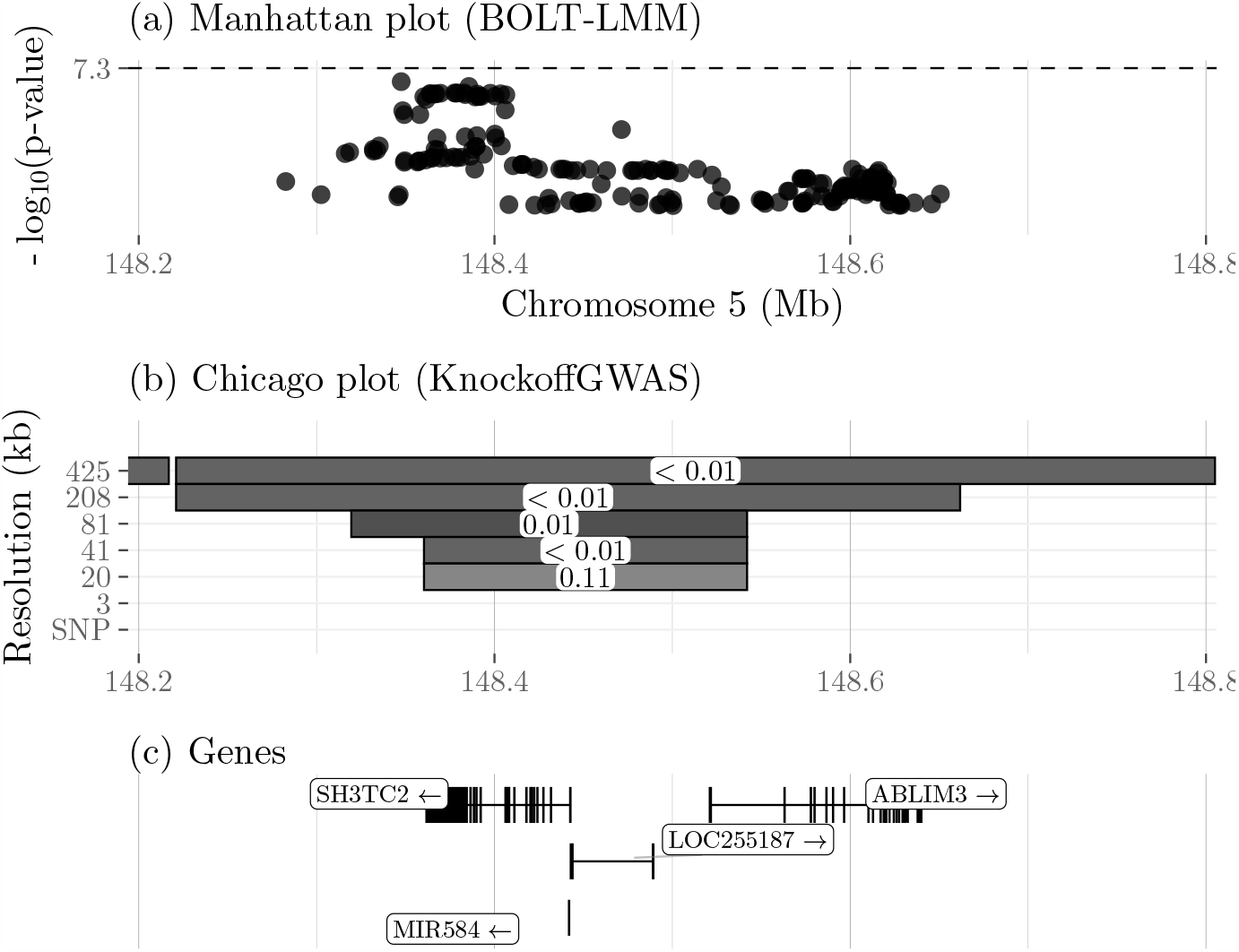
Results of the analysis of UK Biobank data on cardiovascular disease, within a small portion of chromosome five. (a) Marginal p-values computed by BOLT-LMM on the subset of samples with European ancestry [24], for genotyped and imputed variants within this locus. All p-values here are larger than 5 × 10^−8^. (b) Novel findings reported by KnockoffGWAS. The shaded rectangles indicate the genetic segments discovered at different resolutions; the darker ones are more statistically significant, i.e., with a lower estimated local FDP (white labels). (c) Location of genes in the locus spanned by our highest-resolution discovery.

## 4 Discussion

We developed a method for constructing knockoffs that preserve population structure and familial relatedness, as well as LD, thereby obtaining a fully operational strategy for the analysis of GWAS data. In particular, we can now analyze Biobank-scale data sets both efficiently (leveraging the available prior knowledge and the power of virtually any machine learning tools), and agnostically (without assumptions on how the phenotype is related to genetic variation). While we cannot identify causal variants exactly (we know there could be unaccounted confounders, including missing variants), we push further in that direction compared to the traditional pipeline.

The inclusion of related and ethnically diverse individuals is crucial for several reasons [64]. First, large studies are sampling entire populations densely [61, 65], which yields many close relatives. It would be wasteful to discard this information, and potentially dangerous not to account for relatedness. Second, the historical lack of diversity in GWAS (which mostly involve European ancestries) is a well-recognized problem [66, 67] that biases our scientific knowledge and disadvantages the health of under-represented populations. While this issue goes beyond the difficulty of analyzing diverse GWAS data, our work at least helps remove a technical barrier.

GWAS data from different populations are typically analyzed separately, and only later may the results be combined through meta analyses [68, 69], partly out of concern for population structure. However, our method makes such sample splitting unnecessary. By allowing a simultaneous analysis, we can increase power because different LD patterns uncover causal variants more effectively [70]. Furthermore, our discoveries may be useful to better explain phenotypic variation in minority populations [71, 72]. Since the UK Biobank mostly comprises of British individuals, the increase in power resulting from the analysis of other samples can only be relatively small. Nonetheless, we observe some gains when we include non-British individuals. Simulations demonstrate our inferences are valid even when the population is very heterogeneous, suggesting our approach might be particularly suitable for the analysis of more diverse data sets, such as that collected by the Million Veteran Program [73], for example.

Finally, including individuals with diverse ancestries in the GWAS analysis opens new research opportunities. For example, it would be interesting to understand which discoveries are consistently found in different populations and which are more specific, as this may help further weed out false positives, help explain observed variations in certain phenotypes, and possibly shed more light on the underlying biological pathways. Further avenues for future research may focus on the analysis of whole-genome sequencing data including rare variants. The analysis of rare variants involves additional challenges, both computational (our method scales linearly in the number of variables, but even that cost may still be high), and statistical (signals on rare variants are intrinsically more difficult to detect because they result in a smaller effective sample size). Furthermore, it may be more difficult to accurately model the distribution of rare variants, and hence construct valid knockoffs. However, our approach relies on a SHAPEIT HMM with individual-specific reference haplotypes, which is known to be relatively accurate even for rare variants [74, 75]

## Software availability

Our methods are implemented in an open-source software package available from https://msesia.github.io/knockoffgwas/. This includes a standalone program written in C++ which takes as input phased haplotypes in BGEN format [76] and outputs genotype knockoffs at the desired resolution in the PLINK [25] BED format. The package also includes R scripts to partition the genome into contiguous groups of SNPs at different resolutions, compute Lasso-based test statistics, apply the knockoff filter, and visualize the discoveries interactively. Furthermore, the repository contains Bash scripts to connect the different modules of this pipeline and carry out a complete GWAS analysis from beginning to end, an example of which can be conveniently run on a small toy data set provided with the package. Our software is specifically designed for the analysis of large data sets, as it is multi-threaded and memory efficient. Furthermore, knockoffs for different chromosomes can be generated in parallel. For reference, it took us approximately 4 days using 10 cores and 80GB of memory to generate knockoffs on chromosome 1 for the UK Biobank data (*∼*1M haplotype sequences, 600k SNPs, and 600k IBD segments).

## Methods

### The SHAPEIT HMM

We say a sequence of phased haplotypes *H* = (*H*_1_, …, *H*_*p*_) ∈ {0, 1}^*p*^ is distributed as an HMM with *K* hidden states if there exists a vector of latent random variables *Z* = (*Z*_1_, …, *Z*_*p*_) ∈ {1, …, K}^*p*^ such that:

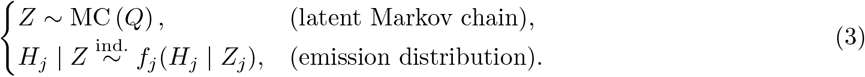

Above, MC (*Q*) is a Markov chain with initial probabilities *Q*_1_ and transition matrices (*Q*_2_, …, *Q*_*p*_).

Taking inspiration from SHAPEIT [41–43], we assume the *i*th haplotype sequence can be approximated as an imperfect mosaic of *K* other haplotypes in the data set, indexed by {*σ*_*i*1_, …, *σ*_*iK*_} ⊆ {1, …, 2*n*}\{i}. (Note the slight overload of notation: *i* denotes hereafter a *phased* haplotype sequence, two of which are available for each individual). We will discuss later how the references are determined; for now, we take them as fixed and describe the other aspects of the model. Mathematically, the mosaic is described by an HMM in the form of (3), where the *i*th Markov chain has:

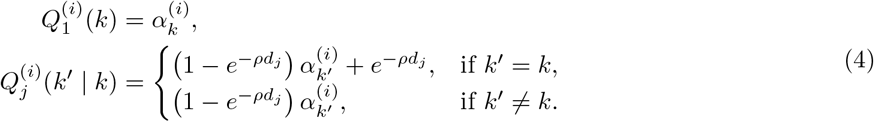

Above, *d*_*j*_ indicates the genetic distance between loci *j* and *j −* 1, which is assumed to be fixed and known (in practice, we will use distances previously estimated in a European population [77], although our method could easily accommodate different distances for different populations). The parameter *ρ >* 0 controls the rate of recombination and can be estimated with an expectation-maximization (EM) technique; see SI. However, we have observed it works well with our data to simply set *ρ* = 1; this is consistent with the approach of SHAPEIT [41–43], which also uses fixed parameters. The positive *α* weights are normalized so that their sum equals one and they can be interpreted as characterizing the ancestry of the *i*th individual. In this paper, we simply set all *α*’s to be equal to 1*/K*, although these parameters could also be estimated by EM (SI). Conditional on *Z*, each element of *H* follows an independent Bernoulli distribution:

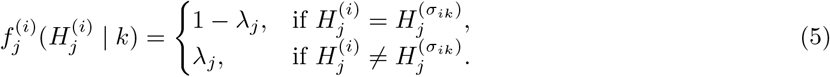

Above, *λ*_*j*_ is a site-specific mutation rate, which makes the mosaic imperfect. Earlier works that first proposed this model suggested analytic formulae for determining *ρ* and *λ* = (*λ*_1_, …, *λ*_*p*_) in terms of physical distances between SNPs and other population genetics quantities [78]. However, we choose to estimate *λ* by EM (SI) since our data set is large. We noticed it works well to also explicitly prevent *λ* from taking extreme values (e.g., 10^−6^ *≤ λ*_*j*_ *≤* 10^−3^).

To save computations and mitigate the risk of overfitting, *K* should not be too large; here, we take *K* = 100. Larger values improve the goodness-of-fit relatively little, while reducing power by increasing the similarity between variables and knockoffs. Having thus fixed *K*, the identities of the reference haplotypes for each *i*, {*σ*_*i*1_, …, *σ*_*iK*_ *}*, are chosen in a data-adaptive fashion as those whose ancestry is most likely to be similar to that of *H*^(*i*)^. Concretely, we can apply Algorithm 1 using the Hamming distance to define haplotypes similarities, chromosome-by-chromosome. Instead of computing pairwise distances between all haplotypes, which would be computationally unfeasible, we first divide them into clusters of size *N*, with *K ≪N≪* 2*n* (i.e., *N ≈*5000), through recursive 2-means clustering, and then we only compute distances within clusters, following in the footsteps of SHAPEIT v3 [43].

#### Algorithm 1 Choosing the HMM reference haplotypes

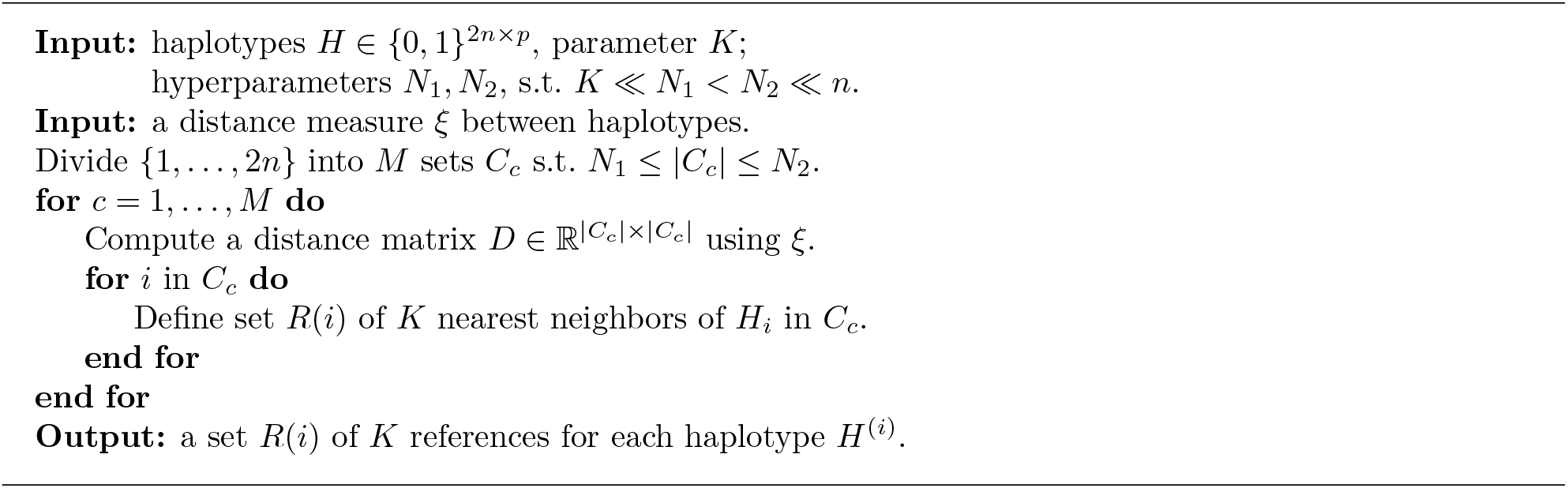

In practice, it is preferable to apply a more sophisticated variation of Algorithm 1, which utilizes a different set of local references in different parts of the chromosome. We will describe this extension later, after discussing the novel knockoff generation algorithm.

### Generating knockoffs preserving population structure

Above, we have described each *H*^(*i*)^ as an HMM conditional on the references, *σ*_*i*1_, …, *σ*_*iK*_ . Therefore, it suffices to apply the general algorithm from [37] on each customized HMM to generate knockoffs. Algorithm 2 outlines the solution; concretely, *Z*^(*i*)^ is sampled from ℙ [*Z*^(*i*)^ |*H*^(*i*)^] with step I of Algorithm 3 in [37], 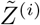 is obtained from step II of the same algorithm, and 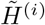 from step III.

#### Algorithm 2 Knockoffs preserving population structure

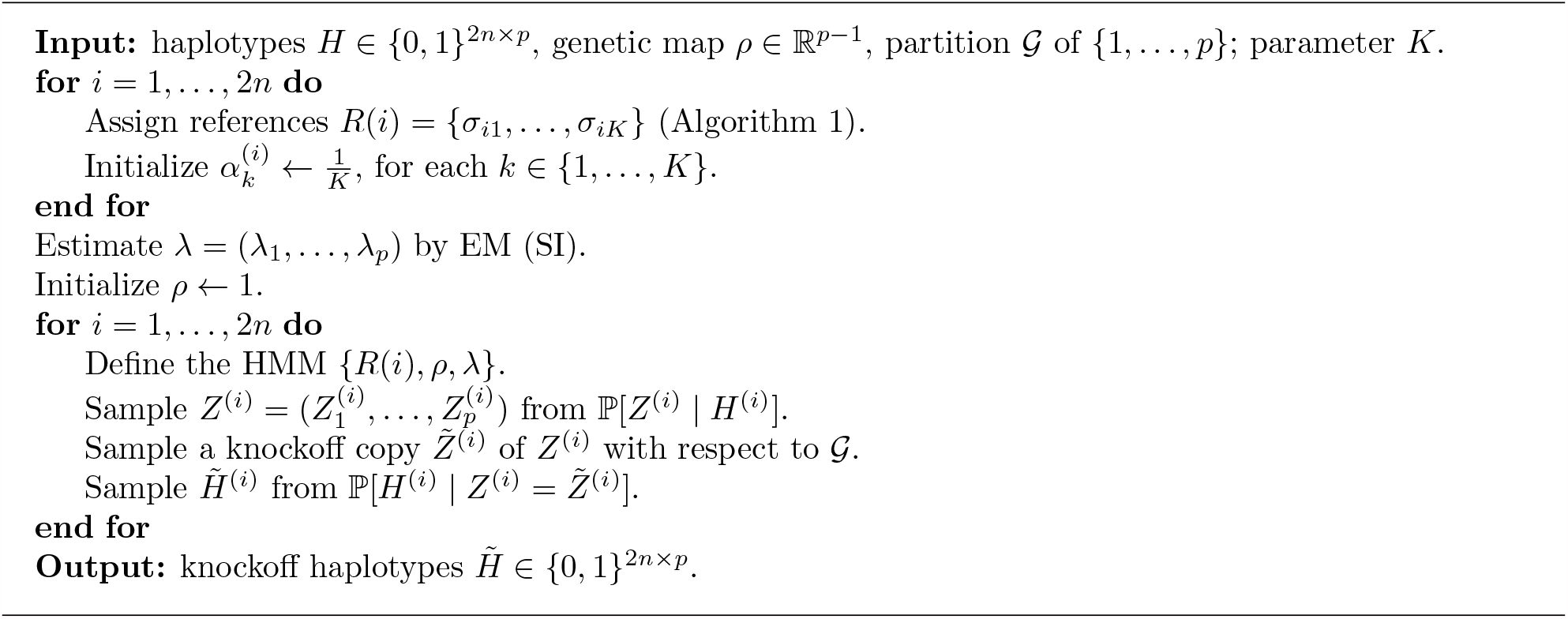

### Knockoffs with local reference motifs based on hold-out distances

Relatedness is not necessarily homogeneous across the genome. This is particularly evident in the case of admixture, which may cause an individual to share haplotypes with a certain population only in part of a chromosome. Therefore, we extend Algorithms 1–2 to accommodate different local references within the same chromosome.

First, we divide each chromosome into relatively wide (e.g., 10 Mb) genetic windows; then, we choose the references separately within each, based on their similarities outside the window of interest. Similarities are computed looking only at the two neighboring windows, to make the references as locally adaptive as possible.

This approach is inspired by SHAPEIT v3 [43], although the latter does not hold out the SNPs in the current window to determine local similarity. Our approach is better suited for knockoff generation because it reduces overfitting—knockoffs too similar to the original variables—and consequently increases power. Having assigned the local references, we apply Algorithm 2 window-by-window. To avoid discontinuities at the boundaries, we consider overlapping windows (expanded by 10 Mb on each side). More precisely, we condition on all SNPs within 10 Mb when sampling the latent Markov chain but then we only generate knockoffs within the window of interest.

### A generalized HMM with IBD segments

We jointly model all haplotypes within an IBD-sharing family *F* = {1, …, *m}*, namely *H*^(*F*)^ = (*H*^(1)^, …, *H*^(*m*)^), as an HMM with a *K*^*m*^-dimensional Markov chain. We write the latter as *Z*^(*F*)^ = (*Z*^(1)^, …, *Z*^(*m*)^), where *Z*^(*i*)^ *∈{*1, …, *K}* ^*p*^. Conditional on *Z*^(*i*)^, each element of *H*^(*i*)^ is independent and follows the same emission distribution as in (3)–(5):

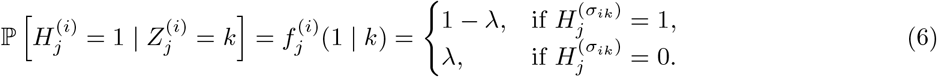

This would reduce to *m* HMMs as in (3)–(5) if the *Z*^(*i*)^’s were independent of each other. However, we couple the *Z*^(*i*)^’s along the (a priori) fixed IBD segments to account for relatedness.

Define *∂*(*i, j*) *⊂ {*1, …, *m}* as the set of haplotype indices that share an IBD segment with *H*^(*i*)^ at position *j*, and *η*_*i,j*_ = 1*/*(1 + |*∂*(*i, j*)|) *∈* (0, 1]. Then, we model *Z*^(*F*)^ as follows: for 1 *< j ≤ p*,

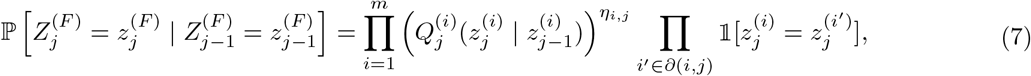

where the transition matrices 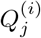 are defined as in (4), while

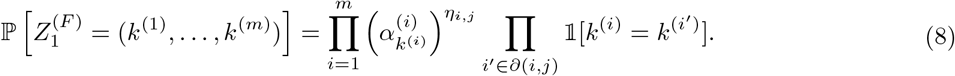

The first term on the right-hand-side of (7) describes the transitions in the Markov chain, while the second term constrains the haplotypes to match along the IBD segments. The purpose of the *η*_*i,j*_ exponent is to make the marginal distribution of each sequence as consistent as possible with the model for unrelated haplotypes. (If *η*_*i,j*_ = 1, Markov chain transitions may occur with significantly different frequency inside and outside IBD segments). In the trivial cases of size-one families, *∂*(1, *j*) =∅ and *η*_1,*j*_ = 1, for all *j ∈{*1, …, *p}*, so (7)–(8) reduce to the model in (4). In general, the latent states for different haplotypes in the same family will be identical along all IBD segments. See Figure S18 (a) for a graphical representation of this model.

### Generating knockoffs preserving IBD segments

The knockoff generation algorithm from [37] would have computational complexity 𝒪 (*npK*^*m*^) for the above model, which is unfeasible for large *n* unless *m* = 1. (Our model is an HMM with a *K*^*m*^-dimensional latent Markov chain, where each vector-valued variable corresponds to the alleles at a specific site for all individuals in the family.) Fortunately, the joint distribution of (*Z*^(*F*)^, *H*^(*F*)^) can be equivalently seen as a more general Markov random field [79] with 2 × *m* × *p* variables, each taking values in {1, …, *K}* or {0, 1}, respectively. See Figure S18 (b) for a graphical representation, where each node corresponds to one of the two haplotypes of an individual at a particular marker. The random field perspective opens the door to more efficient inference and posterior sampling based on message-passing algorithms [80]. Leaving the technical details to the SI, we outline the new procedure in Algorithm 3.

In a nutshell, we follow the spirit of Algorithm 2, with the important difference that the HMM with a *K*-dimensional latent Markov chain of length *p* is replaced by a Markov random field with 2 × *m* × *p* variables; this requires some innovation.

- The *K* haplotype references in the model for each *H*^(*i*)^ are shared by the entire family; see Algorithm S1.
- *Z*^(*F*)^ |*H*^(*F*)^ is sampled with Algorithm S2, which replaces the forward-backward algorithm in [36, 37] with generalized belief propagation [80]. This generally involves some degree of approximation, but it is exact for many family structures.
- 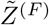 is generated with Algorithm S3, which is a variation of that from [37], circumvententing the computational difficulties of the higher-dimensional model by breaking the couplings between different haplotypes through conditioning [81] upon the extremities of the IBD segments; see also Figure S18.

To clarify, conditioning on the extremities of the IBD segments means we make 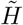 identical to *H* for a few sites in each family, which reduces power only slightly (we consider relatively long IBD segments, so there are few extremities), but greatly simplifies the computations (see SI for a full explanation). It is worth remarking that, for trivial size-one families, this is exactly equivalent to Algorithm 2. Finally, note that the extension to local references with hold-out distances discussed earlier also applies seamlessly here. Our software implements this extension, but we do not explicitly write down the algorithms with local references for lack of space.

#### Algorithm 3 Knockoffs preserving population structure and relatedness

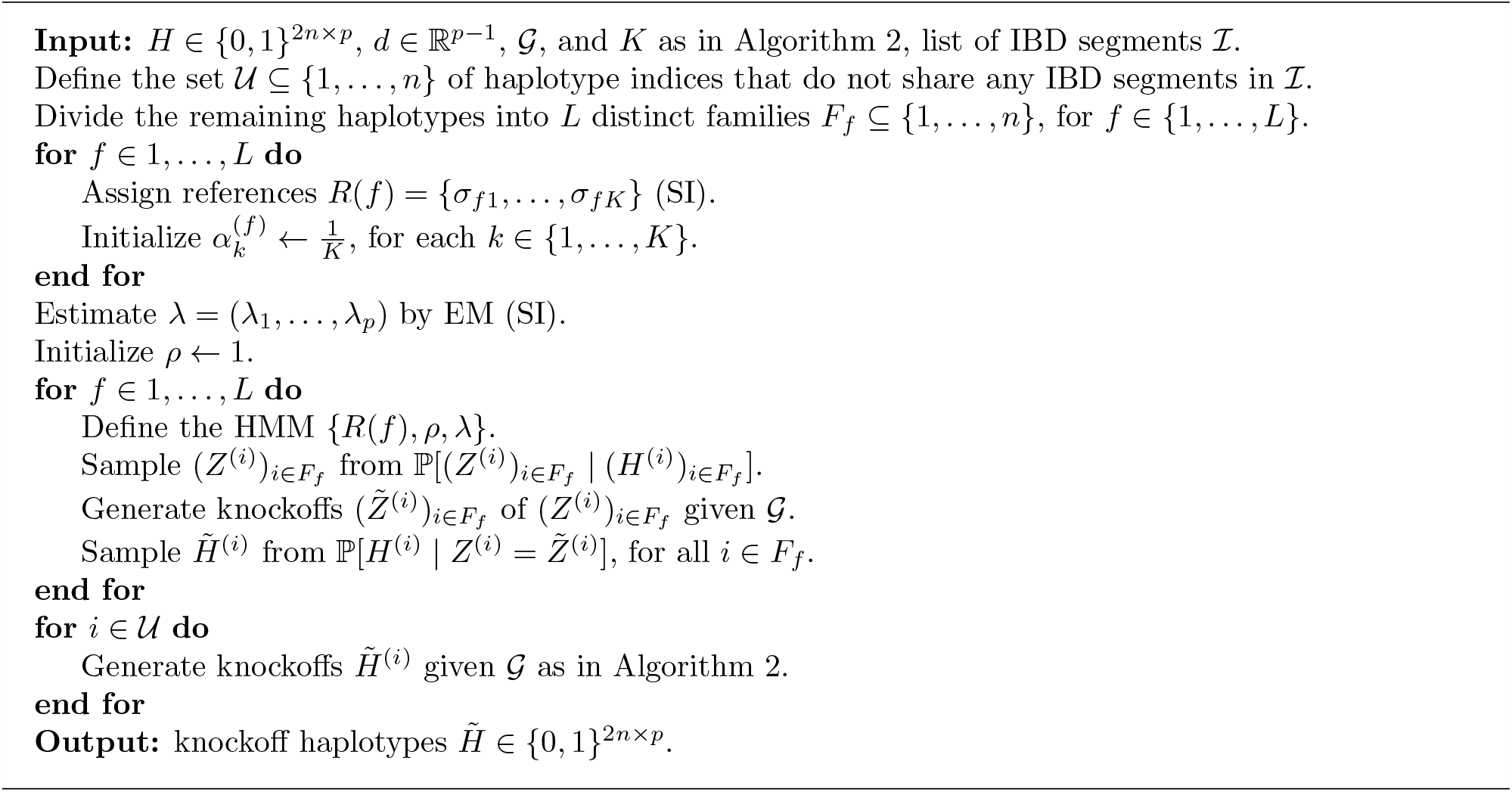

### Quality control and data pre-processing for the UK Biobank

We begin with 487,297 genotyped and phased subjects in the UK Biobank (application 27837). Among these, 147,669 have reported at least one close relative. We define families by clustering individuals with kinship greater or equal to 0.05; then we discard 322 individuals (chosen as those with the most missing phenotypes) to ensure that no families have size larger than 10. This leaves us with 136,818 individuals labeled as related, divided into 57,164 families. The median family size is 2, and the mean is 2.4). The total number of individuals that passed this quality control (including those who have no relatives) is 486,975. We only analyze biallelic SNPs with minor allele frequency above 0.1% and in Hardy-Weinberg equilibrium (10^−6^), among the subset of 350,119 unrelated British individuals previously analyzed in [37]. The final SNPs count is 591,513.

## Acknowledgements

M. S. and S. B. were advised by E. C. at Stanford University. M. S., S. B., E. C. and C. S. were supported by NSF grant DMS 1712800. S. B. was also supported by a Ric Weiland fellowship. E. C. and C. S. were also supported by NSF grant OAC 1934578 and by a Math+X grant (Simons Foundation). We thank Kevin Sharp (University of Oxford) for sharing computer code. We acknowledge the participants and investigators of the UK Biobank, the FinnGen, and the Japan Biobank projects.

## Footnotes

1 If we were to posit a linear model, 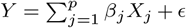, which we do not, then the null hypothesis in (1) would be equivalent to saying that *β*_*j*_ = 0 for all *j* ∈ *G*; see [34, 38].

