## Supplementary Material for "FDR control in GWAS with population structure"

### Supplementary Information for:
### FDR control in GWAS with population structure

Matteo Sesia, Stephen Bates, Emmanuel Candès, Jonathan Marchini, and Chiara Sabatti

March 10, 2021

#### S1 Supplementary Methods

##### S1.a Problem statement and assumptions in presence of confounders

Phenotypes are likely to depend on other variables (either measured or unmeasured) in addition to the genotypes. This increases both the amount of noise in the GWAS data (possibly resulting in lower power) and the risk of *confounding* (inducing spurious associations of the phenotype with non-causal variants). For example, the phenotype $Y^{(i)}$ may be affected by *individual-specific* covariates $U^{(i)}$ (e.g., diet, exercise, or environment) associated with the genotypes—people with different ancestries may differ in both lifestyle and allele frequencies. Furthermore, $Y^{(i)}$ may be influenced by *family* factors $V^{(i)}$, which we assume to be the same for all individuals within a family, and may be dependent on the genotypes, although with some restrictions that we shall discuss below. A mild assumption at this point is that the phenotypes of different individuals are independent of each other conditional on the genotypes, the covariates, and the family factors:

$$P(\mathbf{Y} \mid \mathbf{X}, \mathbf{U}, \mathbf{V}) = \prod_{i=1}^{n} P(Y^{(i)} \mid X^{(i)}, U^{(i)}, V^{(i)}). \tag{S1}$$

Above, $\mathbf{X} \in \{0,1,2\}^{n \times p}$ and $\mathbf{Y} \in \mathcal{Y}^n$ denote the full genotype-phenotype data set, while $\mathbf{U}$ and $\mathbf{V}$ collect the covariates and the family factors, respectively.

For any given genome partition $\mathcal{G}$, our ideal goal would be to know whether the conditional distribution $Y \mid X, U, V$ depends on a group of variants $X_G$, for $G \in \mathcal{G}$. That is, we would like to test:

$$\mathcal{H}_G^* : Y \perp\!\!\!\perp X_G \mid X_{-G}, U, V. \tag{S2}$$

In words, $\mathcal{H}_G^*$ is null if and only if knowledge of $X_G$ provides no information about $Y$ beyond what can be gathered from the knowledge of all other variables. Since not all possible relevant covariates and family factors may be measured in a GWAS, it is unclear how to test (S2) directly. Fortunately, the conditional hypotheses defined in (1) (main paper) are a good practical proxy for (S2) because they account for population structure, thus removing much of the confounding, as explained next. Let us define a factor $A$ that is a function of the genetic information in $X$ and summarizes the ancestry of each individual (i.e., ethnicity, admixture, or family relatedness). Note that we shall not make the definition of $A$ fully explicit (e.g., in terms of some discrete categories or continuous principal components) because the real population structure in a GWAS may be quite complicated (individuals may be stratified even within the same country, ancestries may be admixed, and families may involve more or less distant relatives). Instead, we simply use $A$ as a convenient expository tool here, to rephrase in terms of conditional independence testing the idea that our method accounts for population structure by detecting possibly complex genetic similarities between individuals in the population and then replicating them in the knockoffs (Methods).

We assume the ancestry factor $A$ may affect $U, V$, and $X$, but $U$ and $V$ are independent of $X$ given $A$. Furthermore, the phenotype may be affected by $U, V, X$, but not the other way around (which is biologically sensible); in particular, $Y \perp\!\!\!\perp A \mid U, V, X$. See Figure S1 for a graphical representation of this model. Then, any valid test of

$$\bar{\mathcal{H}}_G : Y \perp\!\!\!\perp X_G \mid X_{-G}, A \tag{S3}$$

is also a valid test of the null hypothesis in (S2). This is the statement in the following proposition.

**Proposition 1.** *In the model assumed above and represented graphically in Figure S1, if the null hypothesis $\mathcal{H}^*_G$ in* (S2) *is true, then $\bar{\mathcal{H}}_G$ in* (S3) *must also be true.*

*Proof.* Suppose $\mathcal{H}^*_G$ is true, so that $Y \perp\!\!\!\perp X_G \mid X_{-G}, U, V$. Since we assumed $Y \perp\!\!\!\perp A \mid U, V, X$, it follows from the contraction property of conditional independence that $Y \perp\!\!\!\perp (X_G, A) \mid X_{-G}, U, V$. Therefore, by the weak union property, we also have that $Y \perp\!\!\!\perp X_G \mid A, X_{-G}, U, V$. Now, note that the conditional distribution of $(Y, U, V, X_G)$ given $(A, X_{-G})$ can be factored as:

$$\begin{aligned}
&p(Y, U, V, X_G \mid A, X_{-G}) \\
&\quad = p(U, V, X_G \mid A, X_{-G}) \cdot p(Y \mid A, X_{-G}, U, V, X_G) \\
&\quad = p(X_G \mid A, X_{-G}) \cdot p(U, V \mid A, X_{-G}) \cdot p(Y \mid A, X_{-G}, U, V, X_G) \\
&\quad = p(X_G \mid A, X_{-G}) \cdot p(U, V \mid A, X_{-G}) \cdot p(Y \mid A, X_{-G}, U, V) \\
&\quad = p(X_G \mid A, X_{-G}) \cdot p(Y, U, V \mid A, X_{-G}).
\end{aligned}$$

Above, the second equality follows from the assumption that $(U, V) \perp\!\!\!\perp X \mid A$. We have thus proved that $(Y, U, V) \perp\!\!\!\perp X_G \mid A, X_{-G}$, which implies $\bar{\mathcal{H}}_G$ in (S3). □

Recall that we presented our knockoffs in Section 2.b (main paper) as negative controls designed to test the hypotheses in (1), which are similar to those in (S3) but do not condition on $A$ explicitly. However, $A$ is a function of the observed genotypes, almost all of which are in included $X_{-G}$ (we only consider relatively groups of SNPs $X_G$ spanning a few hundred kilo-bases at most); thus, testing (1) is almost equivalent to testing (S3). Furthermore, it can be argued even more directly that our knockoffs preserving population structure are valid negative controls for testing (S3) by highlighting they (approximately) satisfy the following stronger—compared to that in (2)—exchangeability property:

$$\left[X^{(F)}, \tilde{X}^{(F)}, A^{(F)}\right]_{\text{swap}(G)} \overset{d}{=} \left[X^{(F)}, \tilde{X}^{(F)}, A^{(F)}\right], \tag{S4}$$

$\forall G \in \mathcal{G}, F \in \mathcal{F}$, as suggested empirically by the statistics in Figure 1. Above, $A^{(F)}$ denotes the vector of ancestry factors for the individuals in family $F$. (Note that $A^{(F)}$ may not necessarily be constant within the family because we model separately the phased haplotypes inherited from each parent; see Methods. This accounts for the possibility that different individuals in the same family may have different patterns of admixture—think for example of a family including two parents of different races and their child.) Despite the presence of $A$ in (S4), this setup is still a special case of that in [1], only with a slightly modified notation. To follow the language of [1] exactly, one would also need a knockoff copy of $A$, but that is unnecessary here because we are only interested in testing the SNPs. Therefore, our knockoffs satisfying (S4) are valid for testing (S3).

Finally, note that the assumption that the genotypes are independent of the covariates and family factors conditional on our practical approximation of the population structure (the factor $A$) is relatively strong and implies our method may not be robust to all possible confounders. This issue prevents us from obtaining rigorous causal inferences, such as those that can be drawn focusing only on parent-child trio data [2]. Furthermore, there is of course an even deeper limitation hiding in the assumption that the genotypes do not directly cause the covariates. For example, if there exists a specific gene that directly influences an individual's predisposition to exercise, regardless of that individual's ancestry, then our method may select that gene as likely to have an effect on cardiovascular disease even if that gene has no a direct biological effect on the disease, only on a behaviour which in turn explains the disease. However, it could be argued this discovery would still be of some interest, and in any case such limitation seems unavoidable if not all possible covariates are measured.

#### S1.b Estimating model parameters by EM

We can estimate the HMM parameters $\theta = (\alpha, \lambda, \rho)$ in (4)–(5), in the main paper, with an expectation-maximization (EM) method. To write down the algorithm explicitly, we begin by noting the log-likelihood of $\theta$ given both the observable, $H$, and latent, $Z$, variables is:

$$\begin{aligned}
\ell(\theta; H, Z) = \log p(H, Z \mid \theta) &= \sum_{i=1}^{n} \log p(H^{(i)}, Z^{(i)} \mid \theta) \\
&= \sum_{i=1}^{n} \log \left\{ \prod_{j=1}^{p} Q_j(Z_j^{(i)} \mid Z_{j-1}^{(i)}) \prod_{j=1}^{p} f_j^{(i)}(H_j^{(i)} \mid Z_j^{(i)}) \right\} \\
&= \sum_{i=1}^{n} \sum_{j=1}^{p} \log Q_j(Z_j^{(i)} \mid Z_{j-1}^{(i)}) + \sum_{i=1}^{n} \sum_{j=1}^{p} \log f_j^{(i)}(H_j^{(i)} \mid Z_j^{(i)}).
\end{aligned}$$

This log-likelihood cannot be directly minimized because we cannot observe $Z$. Instead, given an initial estimate of the model parameters, $\theta^{(t-1)}$, we iteratively update $\theta^{(t)}$ by minimizing

$$\begin{aligned}
\mathcal{L}(\theta, \theta^{(t-1)}) &= \mathbb{E}_Z \left[ \ell(\theta; H, Z) \mid H, \theta^{(t-1)} \right] \\
&= \sum_{i=1}^{n} \sum_{j=1}^{p} \mathbb{E}_Z \left[ \log Q_j(Z_j^{(i)} \mid Z_{j-1}^{(i)}) \mid H^{(i)}, \theta^{(t-1)} \right] \\
&\quad + \sum_{i=1}^{n} \sum_{j=1}^{p} \mathbb{E}_Z \left[ \log f_j^{(i)}(H_j^{(i)} \mid Z_j^{(i)}) \mid H^{(i)}, \theta^{(t-1)} \right].
\end{aligned} \tag{S5}$$

This quantity can be computed and minimized efficiently by leveraging the Markov property, as in the Baum-Welch algorithm.

Let us begin by defining, for any fixed $j \in \{1, \ldots, p\}$, the posterior marginals

$$\gamma_j^{(i)}(k) = \mathbb{P}\left[ Z_j^{(i)} = k \mid H^{(i)}, \theta^{(t-1)} \right].$$

It is well-known that these quantities can be computed efficiently with the classical forward-backward iteration that defines the *expectation* (E) step of the EM algorithm. What remains to be developed explicitly is the *maximization* (M) step of the EM algorithm; we will do this in the following, separately for $\alpha$, $\lambda$, and $\rho$. These are fairly standard calculations but we outline the details here for completeness.

##### S1.b.1 Estimating the site-specific mutation rates

For any $j \in \{1, \ldots, p\}$, the parameter $\lambda_j$ appears in the second term of (S5):

$$\begin{aligned}
&\frac{1}{n}\sum_{i=1}^{n} \mathbb{E}_Z \left[\log f_j^{(i)}(H_j^{(i)} \mid Z_j^{(i)}) \mid H^{(i)}, \theta^{(t-1)}\right] \\
&\quad = \frac{1}{n}\sum_{i=1}^{n}\sum_{z} \log f_j^{(i)}(H_j^{(i)} \mid Z_j^{(i)}) \mathbb{P}\left[Z^{(i)} = z \mid H^{(i)}, \theta^{(t-1)}\right] \\
&\quad = \frac{1}{n}\sum_{i=1}^{n}\sum_{k} \log f_j^{(i)}(H_j^{(i)} \mid Z_j^{(i)} = k) \mathbb{P}\left[Z_j^{(i)} = k \mid H^{(i)}, \theta^{(t-1)}\right] \\
&\quad = \frac{1}{n}\sum_{i=1}^{n}\sum_{k} \log f_j^{(i)}(H_j^{(i)} \mid Z_j^{(i)} = k) \gamma_j^{(i)}(k) \\
&\quad = \frac{1}{n}\sum_{i=1}^{n}\sum_{k} \log \left[(1-\lambda_j)\delta_{H_j^{(i)}, R_j^{(i)}(k)} + \lambda_j(1-\delta_{H_j^{(i)}, R_j^{(i)}(k)})\right] \gamma_j^{(i)}(k) \\
&\quad = \log(1-\lambda_j)\frac{1}{n}\sum_{i=1}^{n}\sum_{k} \delta_{H_j^{(i)}, R_j^{(i)}(k)} \gamma_j^{(i)}(k) + \log(\lambda_j)\frac{1}{n}\sum_{i=1}^{n}\sum_{k}(1-\delta_{H_j^{(i)}, R_j^{(i)}(k)})\gamma_j^{(i)}(k) \\
&\quad = \log(1-\lambda_j)(1-\Gamma_j) + \log(\lambda_j)\Gamma_j,
\end{aligned}$$

where we have defined:

$$\Gamma_j = \frac{1}{n}\sum_{i=1}^{n}\sum_{k}(1-\delta_{H_j^{(i)}, R_j^{(i)}(k)})\gamma_j^{(i)}(k).$$

The above is maximized at $\lambda_j = \Gamma_j$. Therefore, the update rule for $\lambda_j$ in the M step is: $\lambda_j \leftarrow \Gamma_j$.

##### S1.b.2 Estimating the recombination scale

The parameter $\rho$ appears in the first term of (S5) through:

$$\begin{aligned}
\mathbb{E}_Z \left[\log Q_j(Z_j^{(i)} \mid Z_{j-1}^{(i)}) \mid H^{(i)}, \theta^{(t-1)}\right] &= \sum_{z} \log Q_j(z_j \mid z_{j-1}) \mathbb{P}\left[Z^{(i)} = z \mid H^{(i)}, \theta^{(t-1)}\right] \\
&= \sum_{k,l} \log Q_j(k \mid l) \sum_{z_{-(j,j-1)}} \mathbb{P}\left[Z^{(i)} = (k, l, z_{-(j,j-1)} \mid H^{(i)}, \theta^{(t-1)}\right] \\
&= \sum_{k,l} \log Q_j(k \mid l) \mathbb{P}\left[Z_j^{(i)} = k, Z_{j-1}^{(i)} = l \mid H^{(i)}, \theta^{(t-1)}\right].
\end{aligned}$$

By defining

$$\xi_j^{(i)}(k,l) = \mathbb{P}\left[Z_j^{(i)} = k, Z_{j-1}^{(i)} = l \mid H^{(i)}, \theta^{(t-1)}\right],$$

we can write

$$\sum_{i=1}^{n}\sum_{j=1}^{p} \mathbb{E}_Z \left[\log Q_j(Z_j^{(i)} \mid Z_{j-1}^{(i)}) \mid H^{(i)}, \theta^{(t-1)}\right] = \sum_{i=1}^{n}\sum_{j=1}^{p}\sum_{k,l} \log Q_j(k \mid l)\xi_j^{(i)}(k,l).$$

We will discuss later how to compute $\xi$. Now, assume $\xi$ is available and we want to optimize the above with respect to the parameter $\rho$, which is hidden inside the transition matrices $Q$. For simplicity, we also assume

$\alpha_k^{(i)} = 1/K$, $\forall i, k$ (we omit the computations for the general case, which are more complicated). Note that

$$\begin{aligned}
\log Q_j(k \mid l) &= \log\left(\frac{1-b_j}{K} + b_j \delta_{k,l}\right) \\
&= \log\left(\frac{1-b_j}{K}\right) + \left[\log\left(\frac{1-b_j}{K} + b_j\right) - \log\left(\frac{1-b_j}{K}\right)\right]\delta_{k,l} \\
&= \text{const.} + \log(1-b_j) + \left[\log(1+(K-1)b_j) - \log(1-b_j)\right]\delta_{k,l},
\end{aligned}$$

where $b_j = b_j(\rho) = e^{-\rho d_j}$. Therefore,

$$\begin{aligned}
&\frac{1}{n}\sum_{i=1}^{n}\sum_{j=1}^{p}\sum_{k,l} \log Q_j(k \mid l)\xi_j^{(i)}(k,l) \\
&\quad = \frac{1}{n}\sum_{i=1}^{n}\sum_{j=1}^{p}\log(1-b_j)\sum_{k,l}\xi_j^{(i)}(k,l) + \frac{1}{n}\sum_{i=1}^{n}\sum_{j=1}^{p}\left[\log(1+(K-1)b_j) - \log(1-b_j)\right]\sum_{k}\xi_j^{(i)}(k,k) \\
&\quad = \sum_{j=1}^{p}\log(1-b_j) + \sum_{j=1}^{p}\left[\log(1+(K-1)b_j) - \log(1-b_j)\right]\frac{1}{n}\sum_{i=1}^{n}\sum_{k}\xi_j^{(i)}(k,k) \\
&\quad = \sum_{j=1}^{p}\log(1-b_j) + \sum_{j=1}^{p}\left[\log(1+(K-1)b_j) - \log(1-b_j)\right]\Xi_j,
\end{aligned}$$

where we have defined:

$$\Xi_j = \frac{1}{n}\sum_{i=1}^{n}\sum_{k}\xi_j^{(i)}(k,k).$$

It is easy to verify that the above function is strictly quasiconcave in $\rho$, so it can be optimized numerically by solving for its first derivative to be equal to zero. We will include the details of our procedure later for completeness. Meanwhile, note that the computation of $\xi_j^{(i)}(k,l)$ can be easily obtained from the M step:

$$\begin{aligned}
\xi_j^{(i)}(k,l) &= \mathbb{P}\left[Z_{j-1}^{(i)} = l, Z_j^{(i)} = k \mid H^{(i)}\right] \propto \mathbb{P}\left[Z_{j-1}^{(i)} = l, Z_j^{(i)} = k, H^{(i)}\right] \\
&\propto F_{j-1}^{(i)}(l) Q_j^{(i)}(k \mid l) f_j^{(i)}(k \mid H_j^{(i)}) B_j^{(i)}(k) = \bar{\xi}_j^{(i)}(k,l),
\end{aligned}$$

where $F$ and $B$ denote the forward and backward weights. The normalization constant for $\xi_j^{(i)}(k,l)$ is:

$$\begin{aligned}
\sum_k\sum_l \bar{\xi}_j^{(i)}(k,l) &= \sum_k\sum_l F_{j-1}^{(i)}(l) Q_j^{(i)}(k \mid l) f_j^{(i)}(k \mid H_j^{(i)}) B_j^{(i)}(k) \\
&= \sum_l F_{j-1}^{(i)}(l)\sum_k [a_j + b_j\delta_{k,l}] f_j^{(i)}(k \mid H_j^{(i)}) B_j^{(i)}(k) \\
&= a_j\left(\sum_l F_{j-1}^{(i)}(l)\right)\sum_k f_j^{(i)}(k \mid H_j^{(i)}) B_j^{(i)}(k) + b_j\sum_k F_{j-1}^{(i)}(k) f_j^{(i)}(k \mid H_j^{(i)}) B_j^{(i)}(k) \\
&= a_j\sum_k f_j^{(i)}(k \mid H_j^{(i)}) B_j^{(i)}(k) + b_j\sum_k F_{j-1}^{(i)}(k) f_j^{(i)}(k \mid H_j^{(i)}) B_j^{(i)}(k) \\
&= \sum_k f_j^{(i)}(k \mid H_j^{(i)}) B_j^{(i)}(k)\left[a_j + b_j F_{j-1}^{(i)}(k)\right].
\end{aligned}$$

The diagonal elements of $\xi$ are proportional to:

$$\begin{aligned}
\bar{\xi}_j^{(i)}(k,k) &= F_{j-1}^{(i)}(k) Q_j^{(i)}(k \mid k) f_j^{(i)}(k \mid H_j^{(i)}) B_j^{(i)}(k) \\
&= F_{j-1}^{(i)}(k)\left[a_j + b_j\right] f_j^{(i)}(k \mid H_j^{(i)}) B_j^{(i)}(k).
\end{aligned}$$

Recall that we care about

$$\Xi_j = \frac{1}{n}\sum_{i=1}^{n}\sum_{k}\xi_j^{(i)}(k,k) = \frac{1}{n}\sum_{i=1}^{n}\frac{1}{\sum_k\sum_l \bar{\xi}_j^{(i)}(k,l)}\sum_k \bar{\xi}_j^{(i)}(k,k),$$

which we can compute starting from

$$\begin{aligned}\sum_k \bar{\xi}_j^{(i)}(k,k) &= \sum_k F_{j-1}^{(i)}(k)\,(a_j+b_j)\,f_j^{(i)}(k \mid H_j^{(i)})B_j^{(i)}(k) \\ &= (a_j+b_j)\sum_k F_{j-1}^{(i)}(k)f_j^{(i)}(k \mid H_j^{(i)})B_j^{(i)}(k).\end{aligned}$$

Going back to the details of optimizing

$$\frac{1}{n}\sum_{i=1}^{n}\sum_{j=1}^{p}\sum_{k,l}\log Q_j(k \mid l)\xi_j^{(i)}(k,l),$$

note that differentiating with respect to $\rho$ yields:

$$0 = -\sum_{j=1}^{p}\frac{b'_j}{1-b_j} + \sum_{j=1}^{p} b'_j\left[\frac{K-1}{1+(K-1)b_j}+\frac{1}{1-b_j}\right]\Xi_j.$$

By definition of $b_j(\rho) = e^{-\rho d_j}$, it follows that $b'_j = -d_j b_j$. Therefore,

$$\sum_{j=1}^{p}\frac{d_j b_j}{1-b_j} = \sum_{j=1}^{p} d_j b_j\left[\frac{K-1}{1+(K-1)b_j}+\frac{1}{1-b_j}\right]\Xi_j = \Psi(\rho),$$

where we have defined:

$$\Psi(\rho) = \sum_{j=1}^{p} d_j b_j(\rho)\left[\frac{K-1}{1+(K-1)b_j(\rho)}+\frac{1}{1-b_j(\rho)}\right]\Xi_j.$$

Define also $\bar{d} = \frac{1}{p}\sum_{j=1}^{p} d_j$. Then, we want to solve

$$\Psi(\rho) = \sum_{j=1}^{p}\frac{d_j b_j}{1-b_j} = e^{-\rho\bar{d}}\sum_{j=1}^{p}\frac{d_j}{1-b_j}e^{-\rho(d_j-\bar{d})} = e^{-\rho\bar{d}}\Phi(\rho),$$

where

$$\Phi(\rho) = \sum_{j=1}^{p}\frac{d_j}{1-b_j}e^{-\rho(d_j-\bar{d})}.$$

Therefore, we can solve iteratively for $\rho^*$:

$$\rho^* = -\frac{1}{\bar{d}}\log\left(\frac{\Psi(\rho^*)}{\Phi(\rho^*)}\right).$$

Upon convergence (which we observe but do not prove), the solution $\rho^*$ gives the M update for $\rho$ in the EM algorithm: $\rho \leftarrow \rho^*$.

##### S1.b.3 Estimating the motif prevalences

For any fixed $j \in \{1,\dots,p\}$, the parameter $\alpha_k^{(i)}$ appears in the first term of (S5) through:

$$\begin{aligned}\log Q_j(k \mid l) &= \log\left((1-b_j)\alpha_k^{(i)} + b_j\delta_{k,l}\right) \\ &= \log\left((1-b_j)\alpha_k^{(i)}\right) + \left[\log\left((1-b_j)\alpha_k^{(i)}+b_j\right) - \log\left((1-b_j)\alpha_k^{(i)}\right)\right]\delta_{k,l} \\ &= (1-\delta_{k,l})\log\alpha_k^{(i)} + \log\left[(1-b_j)\alpha_k^{(i)}+b_j\right]\delta_{k,l}.\end{aligned}$$

Therefore,

$$\sum_{j=1}^{p}\sum_{k,l}\log Q_j(k \mid l)\xi_j^{(i)}(k,l)$$

$$= \sum_k \log(\alpha_k^{(i)}) \sum_{j=1}^{p}\sum_{l} \xi_j^{(i)}(k,l) - \sum_k \log(\alpha_k^{(i)}) \sum_{j=1}^{p} \xi_j^{(i)}(k,k)$$

$$+ \sum_{j=1}^{p}\sum_{k} \log\left[(1-b_j)\alpha_k^{(i)} + b_j\right] \xi_j^{(i)}(k,k).$$

Differentiating this with respect to $\alpha_k^{(i)}$ gives:

$$0 = \frac{1}{\alpha_k^{(i)}} \sum_{j=1}^{p}\sum_{l} \xi_j^{(i)}(k,l) - \frac{1}{\alpha_k^{(i)}} \sum_{j=1}^{p} \xi_j^{(i)}(k,k) + \sum_{j=1}^{p} \frac{1-b_j}{(1-b_j)\alpha_k^{(i)} + b_j} \xi_j^{(i)}(k,k)$$

$$= \frac{\eta(k) - \bar{\eta}}{\alpha_k^{(i)}} + \sum_{j=1}^{p} \frac{1-b_j}{(1-b_j)\alpha_k^{(i)} + b_j} \xi_j^{(i)}(k,k),$$

where

$$\eta(k) = \sum_{j=1}^{p}\sum_{l} \xi_j^{(i)}(k,l), \qquad \bar{\eta} = \sum_{j=1}^{p} \xi_j^{(i)}(k,k).$$

In order to impose the constraint $\sum_k \alpha_k^{(i)} = 1$, we add a Lagrange multiplier $W$:

$$0 = -W + \frac{\eta(k) - \bar{\eta}}{\alpha_k^{(i)}} + \sum_{j=1}^{p} \frac{1-b_j}{(1-b_j)\alpha_k^{(i)} + b_j} \xi_j^{(i)}(k,k)$$

$$= -W\alpha_k^{(i)} + (\eta(k) - \bar{\eta}) + \alpha_k^{(i)} \sum_{j=1}^{p} \frac{1-b_j}{(1-b_j)\alpha_k^{(i)} + b_j} \xi_j^{(i)}(k,k).$$

Therefore,

$$\alpha_k^{(i)} = \frac{1}{W}\left[\eta(k) - \bar{\eta} + \alpha_k^{(i)} \sum_{j=1}^{p} \frac{1-b_j}{(1-b_j)\alpha_k^{(i)} + b_j} \xi_j^{(i)}(k,k)\right].$$

We can solve this iteratively, setting $W = \sum_k \alpha_k^{(i)}$ after each update of $\alpha^{(i)}$. Upon convergence (which we observe empirically but do not prove), the solution $\alpha^{(i)*}$ will then give the M update in the EM algorithm: $\alpha^{(i)} \leftarrow \alpha^{(i)*}$.

#### S1.c Knockoffs preserving familial relatedness

##### S1.c.1 Choosing the haplotype references

Algorithm S1 modifies Algorithm 1 to ensure: (i) IBD-sharing haplotypes are not used as references for one another; (ii) all haplotypes in the same IBD-sharing family have the same references.

**Algorithm S1** Choosing reference haplotypes preserving familial constraints

**Input:** $H \in \{0,1\}^{2n \times p}$, $K$, and $N_1, N_2$ as in Algorithm 1;
a collection of IBD-sharing families $F_1, \ldots, F_L$, a distance measure $\xi$ between haplotypes.
Divide the haplotypes into $M$ sets $C_c$ using $\xi$ as in Algorithm 1, preserving the family structure.
**for** $c = 1, \ldots, M$ **do**
  Compute a distance matrix $D \in \mathbb{R}^{|C_c| \times |C_c|}$ for all haplotypes in $C_c$.
  **for** $i$ in $C_c$ **do**
    **if** $\exists\, l$ such that $i \in F_l$ **then**
      Define $R(i)$ as the set of $K$ nearest neighbors of $H_i$ in $C_c \setminus F_l$.
    **else**
      Define $R(i)$ as the set of $K$ nearest neighbors of $H_i$ in $C_c$.
    **end if**
  **end for**
**end for**
**for** $l$ in $1, \ldots, L$ **do**
  Initialize $\bar{R}(l) = \cap_{i \in F_l} R(i)$.
  **for** $i \in F_l$ **do**
    Update $R(i) = R(i) \setminus \bar{R}(l)$.
    **if** $|\bar{R}(l)| < K$ **then**
      Update $\bar{R}(l) = \bar{R}(l) \cup R(i)$.
    **else**
      **break**.
    **end if**
  **end for**
  **for** $i \in F_l$ **do**
    Set $R(i) = \bar{R}(l)$.
  **end for**
**end for**
**Output:** a set $R(i)$ of $K$ references for each haplotype $H^{(i)}$.

##### S1.c.2 Posterior sampling via belief propagation

Conditional on $H^{(1:m)}$, the distribution of $Z^{(1:m)}$ is a Markov random field with $m \times p$ variables, characterized by Equations (6)–(8) in the main paper. In order to sample $Z^{(1:m)} \mid H^{(1:m)}$, we implement belief propagation [3] (BP) as follows. For any $i \in \{1, \ldots, m\}$ and $j \in \{1, \ldots, p-1\}$, denote by $\hat{\mu}_{(i,j)\to(i,j+1)} \in \mathbb{R}^K$ the forward message from $Z_j^{(i)}$ to $Z_{j+1}^{(i)}$. It is easy to verify that this must satisfy the following recursive definition:

$$\hat{\mu}_{(i,j)\to(i,j+1)}(k) = \sum_{l=1}^{K} \left[ Q_{j+1}^{(i)}(k \mid l) \right]^{\eta_{i,j+1}} \cdot f_j^{(i)}(H_j^{(i)} \mid l) \cdot \hat{\mu}_{(i,j-1)\to(i,j)}(l) \prod_{i' \in \partial(i,j)} \hat{\mu}_{(i',j)\to(i,j)}(l),$$

where it is understood that $\hat{\mu}_{(i,0)\to(i,1)}(k) = 1$, for all $i$ and $k$. Above, $\hat{\mu}_{(i',j)\to(i,j)}$ indicates the vertical message from $Z_j^{(i')}$ to $Z_j^{(i)}$, for any $i \in \partial(i,j)$. By the BP rules, this satisfies:

$$\hat{\mu}_{(i',j)\to(i,j)}(k) = \sum_{l=1}^{K} \delta_{k,l} \cdot \hat{\mu}_{(i',j-1)\to(i',j)}(l) \cdot \hat{\mu}_{(i',j+1)\to(i',j)}(l) \prod_{i'' \in \partial(i',j)\setminus\{i\}} \hat{\mu}_{(i'',j)\to(i',j)}(l),$$

where $\delta_{k,l} = 1$ if $k = l$ and 0 otherwise. Above, $\hat{\mu}_{(i',j+1)\to(i',j)}(l)$ indicates the backward message from $Z_{j+1}^{(i')}$ to $Z_j^{(i')}$, which is defined recursively as:

$$\hat{\mu}_{(i,j)\to(i,j-1)}(k) = \sum_{l=1}^{K} \left[ Q_j^{(i)}(l \mid k) \right]^{\eta_{i,j}} \cdot f_j^{(i)}(H_j^{(i)} \mid l) \cdot \hat{\mu}_{(i,j+1)\to(i,j)}(l) \prod_{i' \in \partial(i,j)} \hat{\mu}_{(i',j)\to(i,j)}(l).$$

Again, it is understood that $\hat{\mu}_{(i,p+1)\to(i,p)}(k) = 1$, for all $i$ and $k$. Combined, the above updates define a BP algorithm that is in principle already applicable to approximately sample $Z^{(1:m)} \mid H^{(1:m)}$. However, these recursion relations can be simplified by observing that $Z_j^{(i)} = Z^{(i')}$ whenever $i' \in \partial(i,j)$. Therefore, the corresponding nodes in the Markov random field can be collapsed and treated as a single unit in the generalized belief propagation framework [3] (GBP). Thus, after defining

$$\begin{aligned} \phi_j^{(i)}(l) &= f_j^{(i)}(H_j^{(i)} \mid l) \prod_{i' \in \partial(i,j)} f_j^{(i')}(H_j^{(i')} \mid l), \\ \psi_j^{(i)}(k \mid l) &= \left[Q_j^{(i)}(k \mid l)\right]^{\eta_{i,j}} \prod_{i' \in \partial(i,j)} \left[Q_j^{(i')}(k \mid l)\right]^{\eta_{i,j}}, \end{aligned} \tag{S6}$$

it is not difficult to verify that the GBP messages are given by:

$$\begin{aligned} \mu_{(i,j)\to(i,j+1)}(k) &= \sum_{l=1}^{K} \psi_{j+1}^{(i)}(k \mid l) \cdot \phi_j^{(i)}(l) \cdot \mu_{(i,j-1)\to(i,j)}(l) \prod_{i' \in \partial(i,j)\setminus\partial(i,j-1)} \mu_{(i',j-1)\to(i,j)}(l) \\ &\quad \cdot \prod_{i' \in \partial(i,j)\setminus\partial(i,j+1)} \mu_{(i',j+1)\to(i,j)}(l), \\ \mu_{(i,j)\to(i,j-1)}(k) &= \sum_{l=1}^{K} \psi_j^{(i)}(lDOTidk) \cdot \phi_j^{(i)}(l) \cdot \mu_{(i,j+1)\to(i,j)}(l) \prod_{i' \in \partial(i,j)\setminus\partial(i,j+1)} \mu_{(i',j+1)\to(i,j)}(l) \\ &\quad \cdot \prod_{i' \in \partial(i,j)\setminus\partial(i,j-1)} \mu_{(i',j-1)\to(i,j)}(l), \\ \mu_{(i,j)\to(i',j+1)}(k) &= \mu_{(i,j)\to(i,j+1)}(k), \qquad \forall i' \in \partial(i,j+1), \\ \mu_{(i,j)\to(i',j-1)}(k) &= \mu_{(i,j)\to(i,j+1)}(k), \qquad \forall i' \in \partial(i,j-1). \end{aligned} \tag{S7}$$

The GBP rules written above can be simplified even further analytically. Assuming for simplicity that $\alpha_k^{(i)} = 1/K$ (as it is the case in our applications), we can write the transition matrices $Q$ as:

$$Q_j^{(i)}(k \mid l) = Q_j(k \mid l) = a_j + b_j \mathbb{1}[k = l], \qquad a_j = \frac{1}{K}\left(1 - e^{-\rho d_j}\right), \qquad b_j = e^{-\rho d_j}.$$

Therefore,

$$\psi_j^{(i)}(k \mid l) = \left[Q_j^{(i)}(k \mid l)\right]^{\eta_{i,j}} \prod_{i' \in \partial(i,j)} \left[Q_j^{(i')}(k \mid l)\right]^{\eta_{i,j}} = \left[Q_j(k \mid l)\right]^{\eta_{i,j}(1+|\partial(i,j)|)} = Q_j(k \mid l).$$

This simplification allows us to equivalently rewrite the forward update rule in (S7) as:

$$\begin{aligned} &\mu_{(i,j)\to(i,j+1)}(k) \\ &= \sum_{l=1}^{K} \left[a_{j+1} + b_{j+1}\mathbb{1}[k = l]\right] \cdot \phi_j^{(i)}(l) \cdot \mu_{(i,j-1)\to(i,j)}(l) \prod_{i' \in \partial(i,j)\setminus\partial(i,j-1)} \mu_{(i',j-1)\to(i,j)}(l) \\ &\quad \cdot \prod_{i' \in \partial(i,j)\setminus\partial(i,j+1)} \mu_{(i',j+1)\to(i,j)}(l) \\ &= a_{j+1} \sum_{l=1}^{K} \phi_j^{(i)}(l) \cdot \mu_{(i,j-1)\to(i,j)}(l) \prod_{i' \in \partial(i,j)\setminus\partial(i,j-1)} \mu_{(i',j-1)\to(i,j)}(l) \prod_{i' \in \partial(i,j)\setminus\partial(i,j+1)} \mu_{(i',j+1)\to(i,j)}(l) \\ &\quad + b_{j+1}\phi_j^{(i)}(k) \cdot \mu_{(i,j-1)\to(i,j)}(k) \prod_{i' \in \partial(i,j)\setminus\partial(i,j-1)} \mu_{(i',j-1)\to(i,j)}(k) \prod_{i' \in \partial(i,j)\setminus\partial(i,j+1)} \mu_{(i',j+1)\to(i,j)}(k), \end{aligned} \tag{S8}$$

which can be evaluated with complexity $\mathcal{O}(K)$ instead of $\mathcal{O}(K^2)$. Similarly, we can rewrite the backward update rule in such a way that it can also be evaluated at cost $\mathcal{O}(K)$:

$$
\begin{aligned}
&\mu_{(i,j)\to(i,j-1)}(k) \\
&= \sum_{l=1}^{K} [a_j + b_j \mathbb{1}[k=l]] \cdot \phi_j^{(i)}(l) \cdot \mu_{(i,j+1)\to(i,j)}(l) \prod_{i' \in \partial(i,j)\setminus\partial(i,j+1)} \mu_{(i',j+1)\to(i,j)}(l) \\
&\quad \cdot \prod_{i' \in \partial(i,j)\setminus\partial(i,j-1)} \mu_{(i',j-1)\to(i,j)}(l) \\
&= a_j \sum_{l=1}^{K} \phi_j^{(i)}(l) \cdot \mu_{(i,j+1)\to(i,j)}(l) \prod_{i' \in \partial(i,j)\setminus\partial(i,j+1)} \mu_{(i',j+1)\to(i,j)}(l) \prod_{i' \in \partial(i,j)\setminus\partial(i,j-1)} \mu_{(i',j-1)\to(i,j)}(l) \\
&\quad + b_j \cdot \phi_j^{(i)}(k) \cdot \mu_{(i,j+1)\to(i,j)}(k) \prod_{i' \in \partial(i,j)\setminus\partial(i,j+1)} \mu_{(i',j+1)\to(i,j)}(k) \prod_{i' \in \partial(i,j)\setminus\partial(i,j-1)} \mu_{(i',j-1)\to(i,j)}(k).
\end{aligned}
\tag{S9}
$$

The GBP formulation incorporates the IBD-sharing constraints implicitly, removing the vertical messages and the corresponding small loops in the Markov random field. Even though some loops may remain in the graphical model (e.g., if the same two haplotypes share two different IBD segments), these will generally be large compared to the range of background LD, since we only consider relatively long IBD segments. Therefore, we can expect the GBP approximation to work well in general. Furthermore, in many practical cases, the resulting Markov random field is a tree, so the GBP solution will be very fast to compute and provide exact posterior probabilities [3].

GBP randomly initializes the messages $\mu_{(i,j)\to(i',j+1)}$ and $\mu_{(i,j)\to(i',j-1)}$, for all $i, j$ and $i' \in \partial(i,j)$, and then recursively updates them until convergence according to the rules in (S7). Figure S18 shows a schematic of the updates. Even though convergence to an exact solution is only theoretically guaranteed if the underlying graph structure is a tree, the method often performs well in practice, especially if the graph is *locally tree-like* (i.e., it may have long loops but no short ones) [4].

Upon convergence, the posterior distribution of $Z_j^{(i)} \mid H^{(1:m)}$ can be approximated with the product of its incoming messages:

$$
\begin{aligned}
\mathbb{P}\left[Z_j^{(i)} = k \mid H^{(1:m)}\right] &\approx \mu_{(i,j-1)\to(i,j)}(k) \cdot \mu_{(i,j+1)\to(i,j)}(k) \prod_{i' \in \partial(i,j)\setminus\partial(i,j-1)} \mu_{(i',j-1)\to(i,j)}(k) \\
&\quad \cdot \prod_{i' \in \partial(i,j)\setminus\partial(i,j+1)} \mu_{(i',j+1)\to(i,j)}(k).
\end{aligned}
\tag{S10}
$$

Crucially, the above relation is exact in the case of trees, which includes the previously well-known example of a single haplotype sequence [5, 6], as well as many non-trivial family structures (e.g., two haplotypes sharing one IBD segment).

Since we are ultimately interested in sampling all coordinates of $Z^{(1:m)} \mid H^{(1:m)}$ jointly, our procedure does not end with (S10). In general, after sampling $Z_j^{(i)} \mid H^{(1:m)}$ for some $i$ and $j$, one should update the Markov random field by conditioning on the observed value of $Z_j^{(i)}$ and update all messages until convergence before sampling the next variable, which is computationally unfeasible. Fortunately, this procedure can be greatly simplified in our case because we only have relatively long IBD segments, and thus there are few loops in the graphical model. We leverage this fact by first sampling $Z_j^{(i)}$ for all $(i,j)$ in the set $\mathcal{J} \subseteq \{1,\dots,m\} \times \{1,\dots,p\}$ of junction nodes:

$$
\mathcal{J} = \{(i,j) \text{ s.t. } \partial(i,j) \neq \partial(i,j-1) \text{ or } \partial(i,j) \neq \partial(i,j+1)\}. \tag{S11}
$$

Although this requires running $|\mathcal{J}|$ instances of GBP, this quantity will typically be small. Furthermore, warm starts decrease the number of required iterations. Once $Z_j^{(i)}$ has been sampled for all $(i,j) \in \mathcal{J}$, the remaining random field is a collection of disjoint Markov chains, as visualized in Figure S18. Therefore, posterior sampling can be carried out very efficiently with a simple forward-backward procedure that does not involve running BP at each step, as outlined in Algorithm S2.

**Algorithm S2** Posterior sampling preserving familial constraints

---

**Input:** $H \in \{0,1\}^{m \times p}$, $K$, list of IBD segments $\{\partial(i,j)\}_{i\in\{1,\ldots,m\},j\in\{1,\ldots,p\}}$;
  a set $R(i)$ of $K$ references for each haplotype $H^{(i)}$.
Define the list of junction nodes $\mathcal{J} = \{(i,j) \text{ s.t. } \partial(i,j) \neq \partial(i,j-1) \text{ or } \partial(i,j) \neq \partial(i,j+1)\}$.
Initialize the list of active nodes $\mathcal{A} = \{1,\ldots,m\} \times \{1,\ldots,p\}$ and denote its complement as $\mathcal{A}^c$.
Initialize the forward messages $\mu_{(i,j)\to(i,j+1)}(k) = \frac{1}{K}$, for all $i \in \{1,\ldots,m\}$ and $j \in \{1,\ldots,p-1\}$.
Initialize the backward messages $\mu_{(i,j)\to(i,j-1)}(k) = \frac{1}{K}$, for all $i \in \{1,\ldots,m\}$ and $j \in \{2,\ldots,p\}$.
**for** $(i^*,j^*) \in \mathcal{J} \cap \mathcal{A}$ **do**
  **while** messages not converged **do**
    **for** $j = 1,\ldots,p-1$ **do**
      **for** $i = 1,\ldots,m$ **do**
        **if** $(i,j) \in \mathcal{A}$ **then**
          Update $\mu_{(i,j)\to(i,j+1)}(k)$, for all $k \in \{1,\ldots,K\}$, according to (S8).
        **end if**
      **end for**
    **end for**
    **for** $j = p,\ldots,2$ **do**
      **for** $i = 1,\ldots,m$ **do**
        **if** $(i,j) \in \mathcal{A}$ **then**
          Update $\mu_{(i,j)\to(i,j-1)}(k)$, for all $k \in \{1,\ldots,K\}$, according to (S9).
        **end if**
      **end for**
    **end for**
  **end while**
  Approximate the posteriors $w^{(i^*)}_{j^*}(k)$ of $Z^{(i^*)}_{j^*} = k \mid H^{(1:m)}, \{Z^{(i)}_j\}_{(i,j)\in\mathcal{A}^c}$ based on (S10).
  Sample $Z^{(i^*)}_{j^*}$ from $\mathbb{P}[Z^{(i^*)}_{j^*} = k] = w^{(i^*)}_{j^*}(k)$.
  Update the list of active nodes: $\mathcal{A} \leftarrow \mathcal{A} \setminus \{(i^*,j^*)\}$.
  Update the Markov random field: $\phi^{(i^*)}_{j^*}(k) \leftarrow \mathbb{1}[k = Z^{(i^*)}_{j^*}]$, for each $k \in \{1,\ldots,k\}$.
  **for** $i' \in \partial(i^*,j^*)$ **do**
    Set $Z^{(i')}_{j^*} \leftarrow Z^{(i^*)}_{j^*}$.
    Update the list of active nodes: $\mathcal{A} \leftarrow \mathcal{A} \setminus \{(i',j^*)\}$.
    Update the Markov random field: $\phi^{(i')}_{j^*}(k) \leftarrow \mathbb{1}[k = Z^{(i')}_{j^*}]$, for each $k \in \{1,\ldots,k\}$.
  **end for**
**end for**
Sample each disjoint segment of $\{Z^{(i)}_j\}_{(i,j)\in\mathcal{J}^c} \mid H^{(1:m)}, \{Z^{(i)}_j\}_{(i,j)\in\mathcal{J}}$, with standard forward-backward [6].
**Output:** a latent Markov random field $Z \in \{1,\ldots,K\}^{m \times p}$ that preserves the IBD structure.

---

##### S1.c.3 Knockoff generation via conditioning

Having sampled $Z^{(1:m)} \mid H^{(1:m)}$ with the procedure described above, we proceed to develop a method for generating knockoff copies $\tilde{Z}^{(1:m)}$. Even though constructing exact knockoffs for a general Markov random field may be computationally unfeasible, we can simplify the problem by conditioning on some variables [7]. In particular, we condition on all variables at the junction of any IBD segment, i.e., those in the set $\mathcal{J}$ defined in (S11). This transforms the model for the remaining variables into a collection of disjoint one-dimensional chains, for which knockoffs can be generated independently with existing methods [5, 6]; see Figure S18. This solution is summarised in Algorithm S3.

**Algorithm S3** Related knockoff haplotypes via conditioning

**Input:** $H \in \{0,1\}^{m \times p}$, $d \in \mathbb{R}^{p-1}$, $\mathcal{G}$, and $K$ as in Algorithm 2;
IBD segments $\{\partial(i,j)\}_{i\in\{1,\ldots,m\},j\in\{1,\ldots,p\}}$;
a set $R(i)$ of $K$ references for each haplotype $H^{(i)}$;
Markov random field states $Z \in \{1,\ldots,K\}^{m\times p}$.
Define the list of junction nodes $\mathcal{J} = \{(i,j) \text{ s.t. } \partial(i,j) \neq \partial(i,j-1) \text{ or } \partial(i,j) \neq \partial(i,j+1)\}$.
**for** $(i,j) \in \mathcal{J}$ **do**
  Define $G$ as the group in partition $\mathcal{G}$ to which variant $j$ belongs.
  **for** $j' \in G$ **do**
    Expand the list of junction nodes: $\mathcal{J} \leftarrow \mathcal{J} \cup \{(i,j')\}$.
  **end for**
**end for**
**for** $(i,j) \in \mathcal{J}$ **do**
  Make a trivial knockoff: $\tilde{Z}_j^{(i)} \leftarrow Z_j^{(i)}$.
**end for**
**for** each connected component $C$ in $\{1,\ldots,m\} \times \{1,\ldots,p\} \setminus \mathcal{J}$ **do**
  Generate group knockoffs $\{\tilde{Z}_j^{(i)}\}_{(i,j)\in C}$ of $\{Z_j^{(i)}\}_{(i,j)\in C} \mid \{Z_j^{(i)}\}_{(i,j)\in\mathcal{J}}$ as in previous work [6].
**end for**
**Output:** knockoff matrix $\tilde{Z} \in \{1,\ldots,K\}^{m\times p}$.

### S2 Supplementary Notes

#### S2.a Additional numerical experiments

#### S2.b Setup

We consider here additional simulations to test our method on real genotypes and synthetic phenotypes, focusing on smaller subsets individuals from the UK Biobank data set. There are two reasons why these experiments are informative. Firstly, they allow us to test the robustness of our method to very strong population structure, by eliminating most of the unrelated British individuals, which make the entire data set relatively homogeneous overall. Secondly, they are computationally cheaper, which allows us to conveniently repeat the experiments for several random realizations of the phenotypes.

In these experiments, the feature importance measures for each SNP are computed in three alternative ways: by fitting the Lasso with cross-validation and taking the absolute value of the estimated regression coefficients (as in the main paper); by running BOLT-LMM [8] and taking the negative logarithm of the marginal p-values; and by performing univariate logistic regression (in the case of binary phenotypes) and taking the negative logarithm of the marginal p-values. These models are designed to predict $Y$ given $[X, \tilde{X}]$; in the first two cases we also include the top 10 principal components of the genotype matrix (computed on the entire UK Biobank data set) as additional covariates. Then, the feature importance measures $T_j$ and $\tilde{T}_j$, for the $j$th SNP and its corresponding knockoff, are combined in the usual way to define the knockoff test statistics for each group $G \subseteq \{1,\ldots,p\}$ of variables: $W_G = \sum_{j\in G} T_j - \sum_{j\in G} \tilde{T}_j$.

##### S2.b.1 Knockoffs preserving population structure

We focus here on 10,000 unrelated individuals from the UK Biobank with one of 6 different self-reported ancestries (Table S1). We simulate continuous phenotypes, conditional on the true genotypes, from a homoscedastic linear model with 500 causal variants distributed uniformly across the genome; the total heritability is varied as a control parameter. We apply KnockoffGWAS on these data using knockoffs generated based on either the SHAPEIT or the fastPHASE model. Figure S14 shows the histogram of test statistics computed either with the usual Lasso-based approach, or with BOLT-LMM [9]; the LMM is less powerful, but it makes the increased robustness of the SHAPEIT knockoffs even more apparent. The distribution of test statistics should be symmetric around zero for null groups (i.e., those without causal variants) if the knockoffs are valid. The statistics obtained with the SHAPEIT model satisfy this property, while the fastPHASE HMM leads to a rightward bias, which may result in an excess of false positives. The power and FDR (using the Lasso-based statistics) are compared in Figure S13: the SHAPEIT model leads to slightly lower power, but always controls the FDR. Figure S13 also summarizes findings at different resolutions by counting only the most specific ones [6].

##### S2.b.2 Knockoffs preserving familial relatedness

Here, we test our method on 10,000 British individuals in 4,900 self-reported families; see Table S3 for details. According to the results of RaPID [10], these individuals share a total of 723,454 IBD segments. Their mean width is 19.6 Mb, or 26.1 cM, and each contains 4238 SNPs on average. We generate knockoffs preserving these IBD segments, and compare the results with those obtained disregarding relatedness.

Figure S9 shows that knockoffs would not preserve IBD segments if we did not explicitly enforce such constraint, especially at low resolution. Furthermore, the diagnostics in Figure S9 confirm that our method correctly preserves LD, and that accounting for relatedness does not decrease power; to the contrary, it can increase it by ensuring that closely related haplotypes are not used as references for one another, which would reduce the desired contrast between genotypes and knockoffs.

We simulate binary phenotypes from a liability threshold (probit) model with 100 uniformly distributed causal variants; the numbers of cases and controls are balanced. (We consider binary phenotypes, as opposed to continuous phenotypes as in the previous section, simply to highlight the flexibility of our method, which is equally valid regardless of the distribution of the trait). We include in this model an additive random term for each family, mimicking shared family factors, whose strength is smoothly controlled by a parameter $\gamma \in [0, 1]$. More precisely, the latent Gaussian variable for the $i$th individual in the probit model is given by:

$$L^{(i)} = \sum_{j=1}^{p} \beta_j X_j^{(i)} + \gamma V^{(f)} + \sqrt{1 - \gamma^2} \epsilon^{(i)},$$

where $E^{(f)}$ and $\epsilon^{(i)}$ are independent standard normal random variables, $f$ denotes the family to which individual $i$ belongs, and $\gamma \in [0, 1]$. The magnitude of the nonzero genetic coefficients $\beta$ is varied as a parameter, to control the total heritability of the trait.

Therefore, the phenotypes of different individuals in the same family are conditionally independent given the genotypes if $\gamma = 0$, while identical twins will always have the same phenotype if $\gamma = 1$. In theory, family factors may introduce spurious associations, unless the knockoffs account for familial relatedness.

Figure S15 reports FDR and power at low-resolution, with and without preserving relatedness. This shows that preserving IBD segments enables FDR control even with extremely strong family factors ($\gamma = 1$), with virtually no power loss. However, the SHAPEIT model is reasonably robust even if relatedness is ignored, especially at higher resolution. This partly depends on the multivariate importance statistics used here (i.e., sparse logistic regression); in fact, the knockoff filter applied with marginal importance statistics is more vulnerable to confounding, as also illustrated in Figure S15.

#### S2.c Enrichment analysis with external summary statistics

We perform an enrichment analysis using external summary statistics from the Japan Biobank project [11] for the continuous traits, and from the FinnGen resource [12] for all binary traits except respiratory disease. These summary statistics were computed on data independent of those in the UK Biobank, but some care must be exercised in the interpretation of these results because: (a) the external statistics measure marginal association, not conditional importance; (b) the external sample sizes are smaller than ours, which limits power.

For each group of SNPs $G$ in the genome partition at the 20 kb resolution, we compute a chi-square statistics with Fisher's method: $\chi^2_G = -\sum_{j \in G} \log(p_j)$, where $\{p_j\}_{j \in G}$ denotes the set of external marginal p-values within the region spanned by $G$. Since the UK Biobank and the FinnGen project are based on different genome builds, our discoveries are matched to the external p-values after appropriately lifting the physical positions. We then define: $\{\chi^2_G\}_{S^{\text{novel}}}$ as the collection of external Fisher statistics corresponding to our novel discoveries in $S^{\text{novel}}$; $\{\chi^2_G\}_{S^{\text{confirmed}}}$ as the collection of external Fisher statistics corresponding to our previously confirmed discoveries (either confirmed by BOLT-LMM, or by the other studies based on the GWAS catalog, and the Japan Biobank or the FinnGen resource at the genome-wide significance level); and $\{\chi^2_G\}_{\text{background}}$ as the set of Fisher statistics for groups that are neither in $S^{\text{confirmed}}$ nor in $S^{\text{novel}}$.

We take the empirical distribution of $\{\chi^2_G\}_{\text{background}}$ as the null distribution, and invert it to compute an approximate enrichment p-value for each group in $S^{\text{novel}}$; we refer to these as $p^{\text{enrich}}_G$. The null hypothesis, under which the $p^{\text{enrich}}_G$ would be approximately uniform, is that the Fisher statistics for the novel discoveries have the same distribution as those in $\{\chi^2_G\}_{\text{background}}$; for instance, we expect this would be the case if all selected SNPs were independent of the phenotype. In theory, we could use these p-values with any multiple testing procedure; however, this turns out to have little power, due to the small sample size (compared to the UK Biobank) of the external data. However, it is clear that the distribution of $\{p^{\text{enrich}}_G\}_{S^{\text{novel}}}$ is not uniform, which suggest many discoveries are non-null. Therefore, we take an empirical Bayes approach to estimate the proportion of non-null discoveries [13], as implemented by the "quantile" method in the `fdrtool` R package [14]. This estimates the proportion of null enrichment p-values, which we bootstrap 10,000 times to assess its uncertainty. Tables 3 and S11 are based on the mean bootstrap values, while Table S12 report 90% confidence intervals.

### S3 Figures

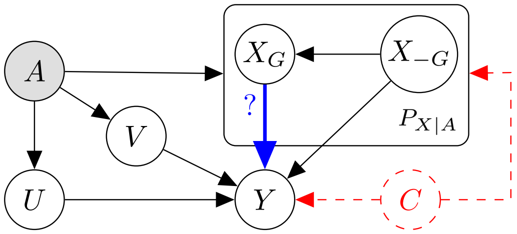


Figure S1: Graphical representation of the assumed relation between genotypes ($X$), phenotype ($Y$), ancestry ($A$), family factors ($V$), and other relevant covariates ($U$). Our method is designed to test the direct effect of any subset of genotypes $X_G$ on the phenotype. The node $C$ represents possible unaccounted confounders.

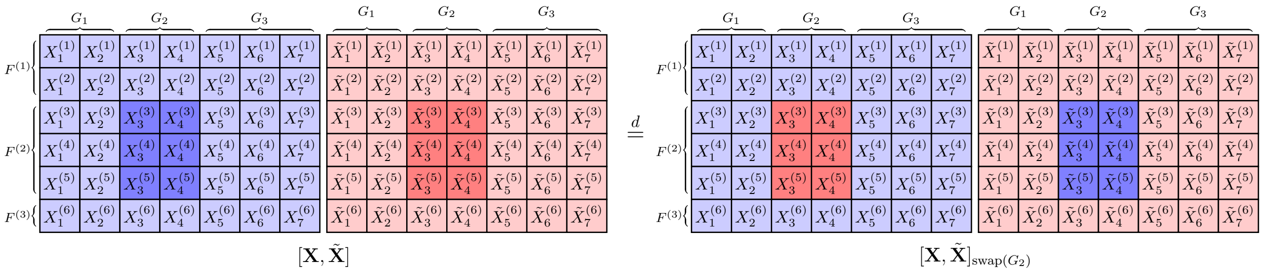


Figure S2: Visualization of the knockoff exchangeability property defined in (2), within a toy example with 6 individuals (divided into 3 families) and 7 variants (partitioned into 3 groups). In this example, the swapping operator for group $G_2$ is applied to the second family, $F^{(2)}$, and the ancestry variable $A$ is omitted.

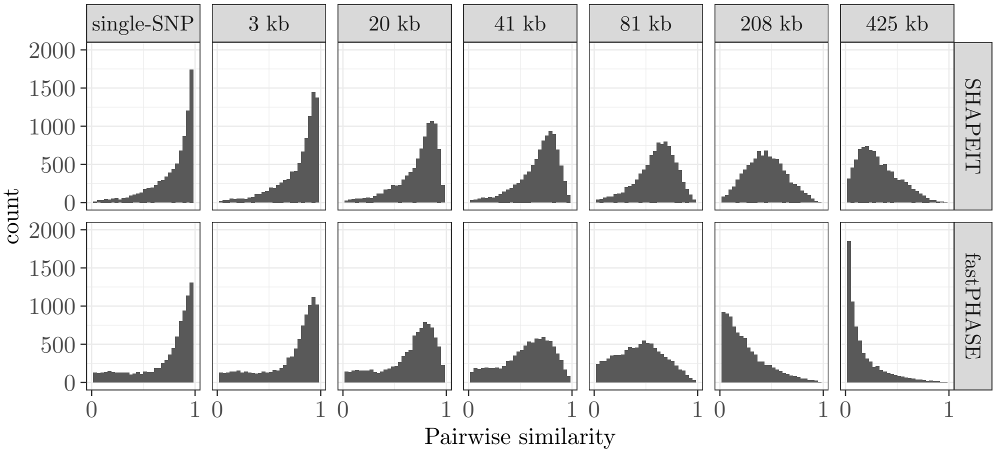


(a) Distribution of pairwise correlations for different variants on chromosome 22.

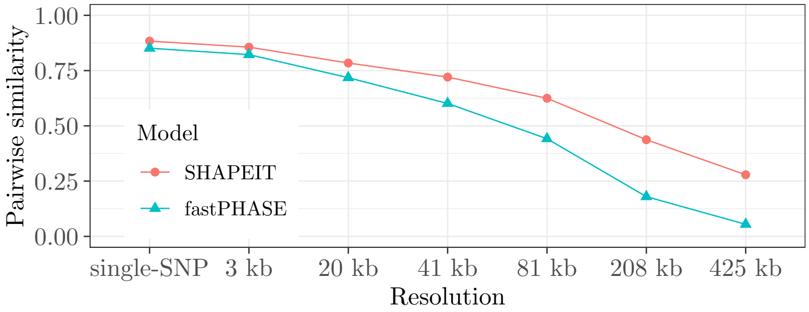


(b) Average pairwise correlations over chromosome 22.

Figure S3: Absolute pairwise correlation between genotypes and knockoffs on chromosome 22, as a function of the knockoff resolution. These statistics are computed on 10,000 UK Biobank samples with diverse ancestries, as in Figure 1. (a) Histograms. (b) Absolute correlation averaged over all variants, as a function of the resolution. Lower absolute pairwise correlations with the genotypes indicate more powerful knockoffs.

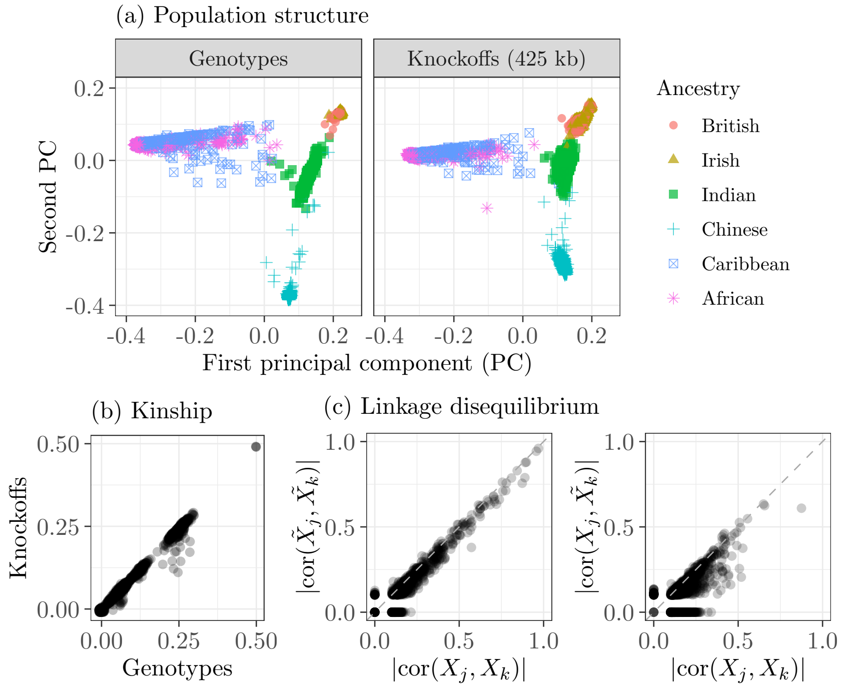


Figure S4: Exchangeability of knockoffs and genotypes in the UK Biobank. (a) Principal component analysis for 10k individuals with diverse ancestries, separately for genotypes and knockoffs. (b) Kinship coefficients between 2000 pairs of related individuals, computed separately on genotypes and knockoffs. (c) Pairwise absolute correlations between nearby variants on chromosome 22 (minor allele frequency $\geq 0.01$) for the same individuals as in (a), with (left) or without (right) swapping genotypes ($X$) and knockoffs ($\tilde{X}$). Resolution: 425 kb. In the case of (b), we only show pairs of variants in different groups. Other details are as in Figure 1.

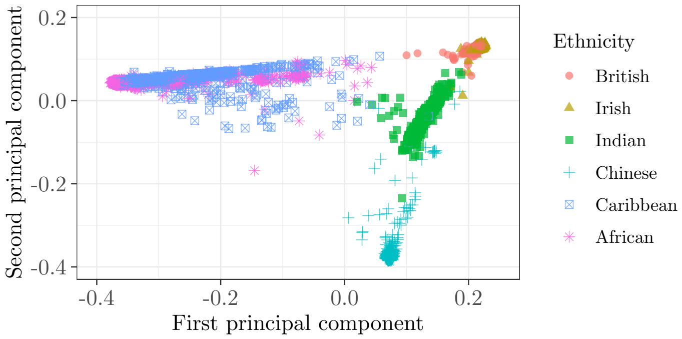


(a) Real genotypes.

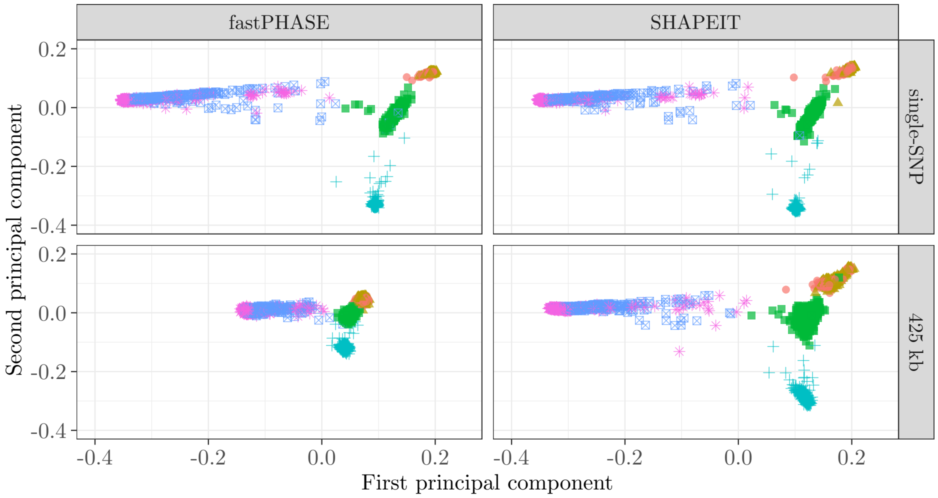


(b) Knockoffs.

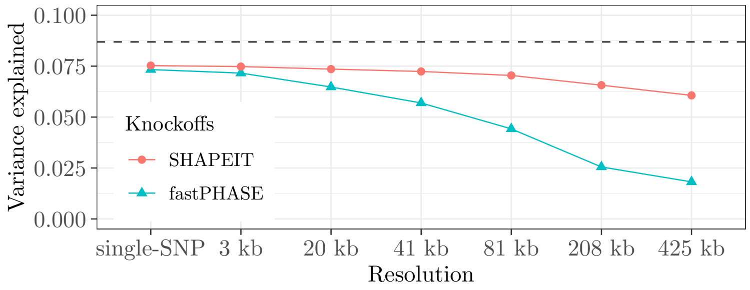


(c) Comparison of PCA on genotypes and knockoffs.

Figure S5: PCA of individuals with diverse ancestries, and of knockoffs constructed based on different HMMs. (a) The first two genetic principal components of 10,000 individuals in the UK Biobank (as in Figures 1 and S4) are compared to (b) the corresponding quantities computed on knockoffs at different resolutions. The knockoffs based on the SHAPEIT HMM preserve population structure quite accurately, even at low resolution. By contrast, the fastPHASE HMM tends to produce knockoffs that shrink together individuals with diverse ancestries, thus breaking population structure. (c) Proportion of genetic variance explained by the first ten principal components of knockoffs at different resolutions, for samples with diverse ancestries. The dashed horizontal line indicates the corresponding quantity computed on the original data.

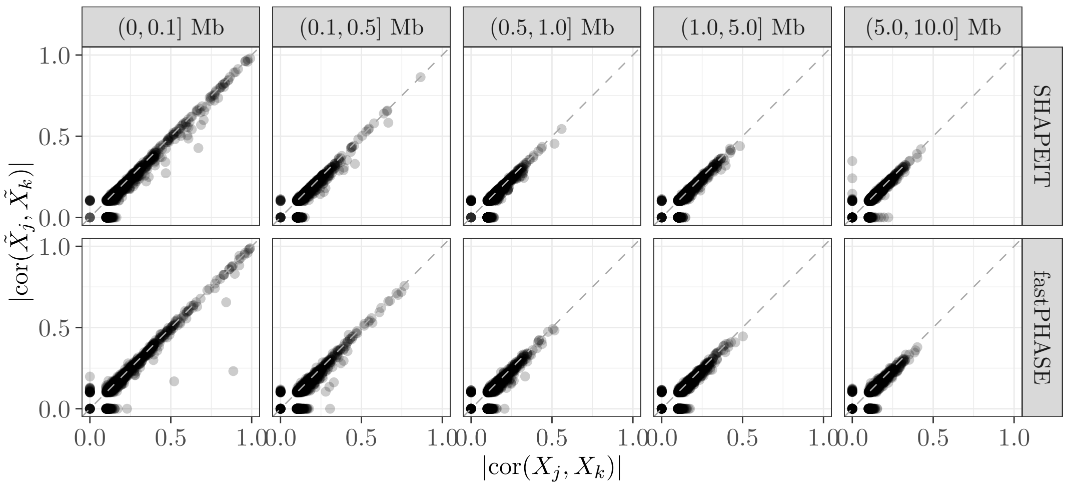


(a) High-resolution knockoffs.

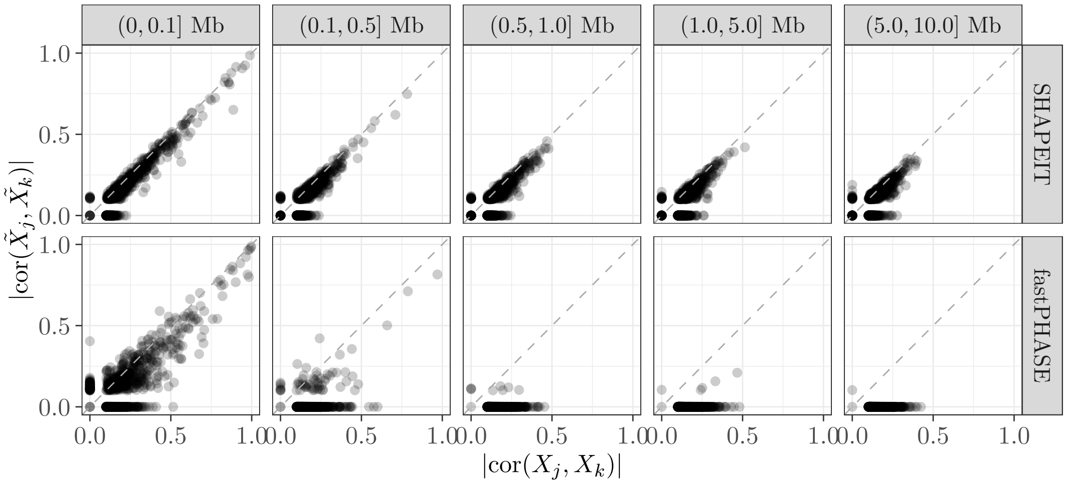


(b) Low-resolution knockoffs.

Figure S6: Knockoff exchangeability measured in terms of pairwise correlations between different SNPs, among 10,000 individuals with diverse ancestries, as in Figure 1. We compare $|\text{cor}(X_j, X_k)|$ with $|\text{cor}(\tilde{X}_j, \tilde{X}_k)|$, for $j, k \in \{1, \ldots, p\}$, as a function of the distance between variants $j$ and $k$ on chromosome 22. Only 1000 randomly chosen points are shown, for clarity. Variants with minor allele frequency smaller than 0.01 are not shown here, due to the limited sample size. These diagnostics should approximately lie on the 45-degree line if the knockoffs are valid [6]. (a) Genome partition into single-SNP groups. (b) Genome partition into 425 kb-wide groups.

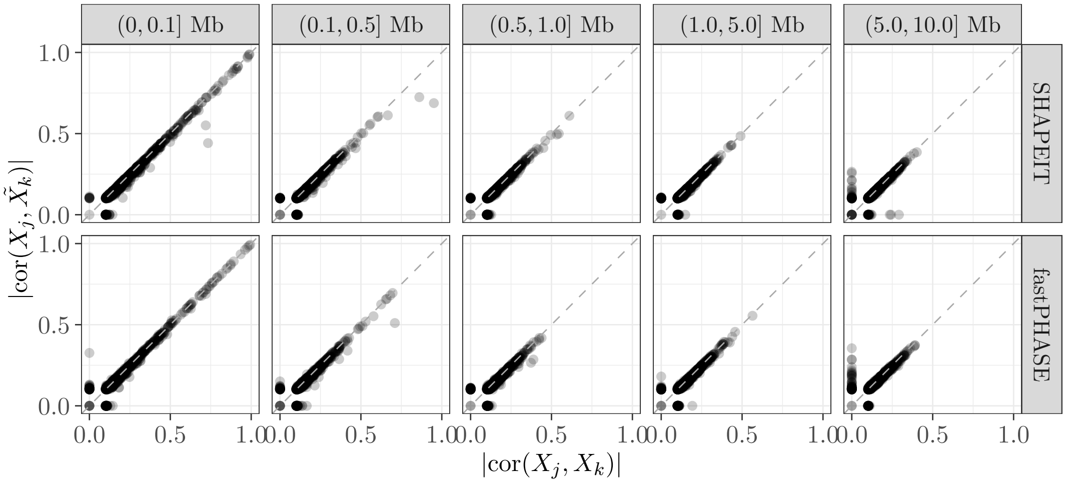


(a) High-resolution knockoffs.

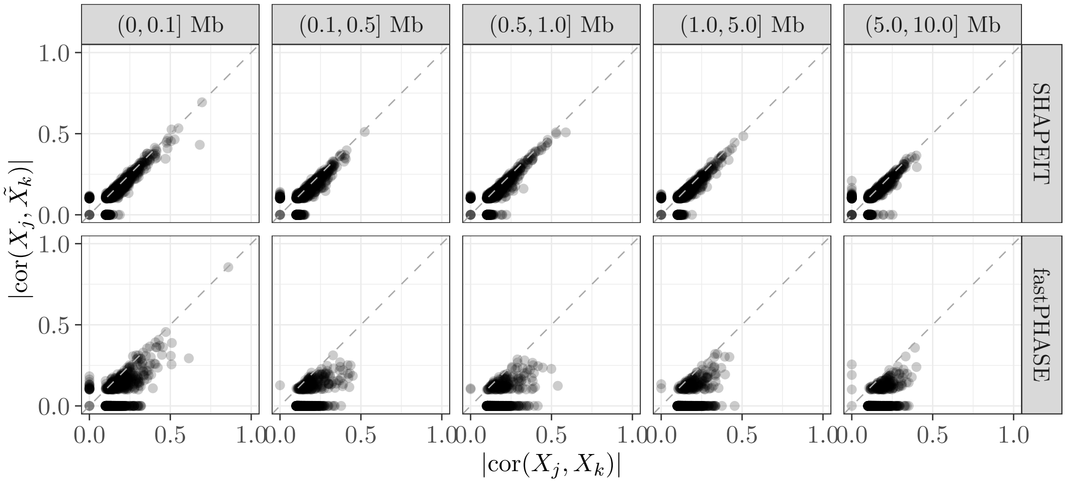


(b) Low-resolution knockoffs.

Figure S7: Additional exchangeability diagnostics comparing $|\text{cor}(X_j, X_k)|$ with $|\text{cor}(X_j, \tilde{X}_k)|$, for $j, k$ in different groups, as a function of the distance between variants $j$ and $k$. (a) Genome partition into single-SNP groups. (b) Genome partition into 425 kb-wide groups. Other details are as in Figure S6.

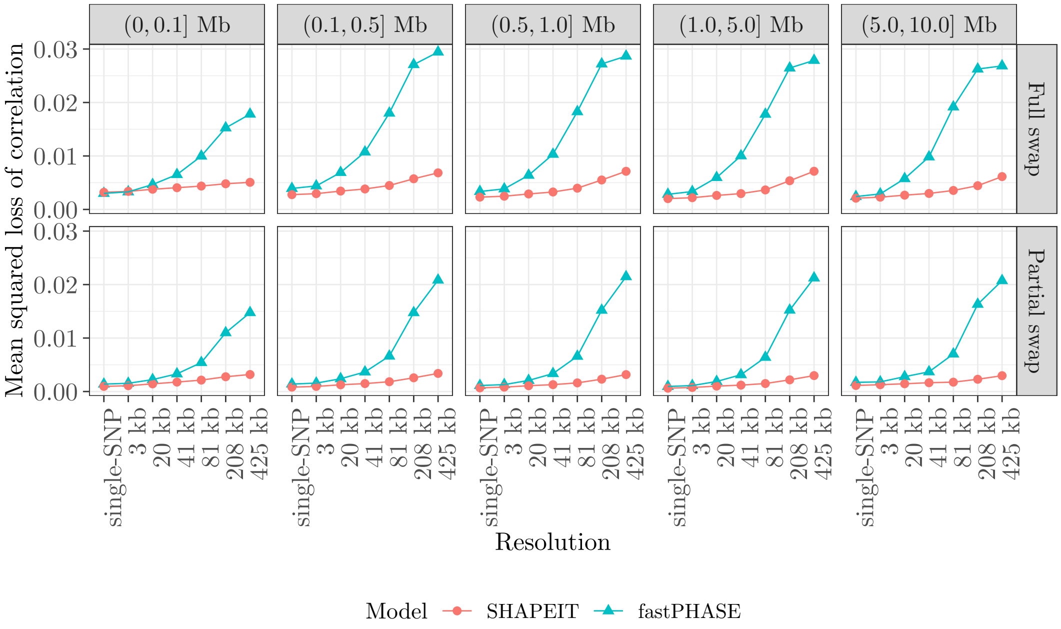


Figure S8: Knockoff exchangeability measured in terms of pairwise correlations between different SNPs, as in Figures S6–S7. The quantity on the vertical axis measures the average distances from the 45-degree line in the scatter plots of Figures S6–S7, including also intermediate levels of resolutions. This is defined as $[\mathrm{cor}(X_j, X_k) - \mathrm{cor}(\tilde{X}_j, \tilde{X}_k)]^2$ (top), or $[\mathrm{cor}(X_j, X_k) - \mathrm{cor}(X_j, \tilde{X}_k)]^2$ (bottom), each averaged over pairs of variables $j, k$ whose physical distances are within the specified range. Valid knockoffs should have values close to zero.

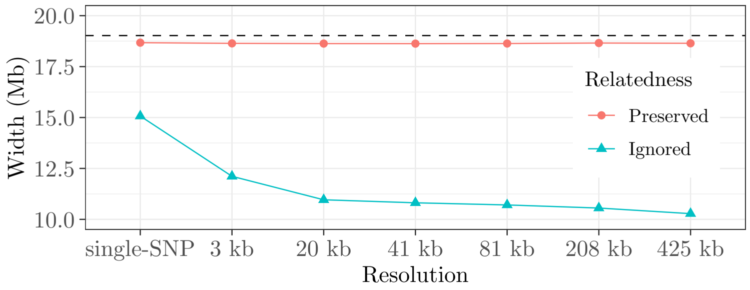


(a) Relatedness (values closer to horizontal dashed line are preferable).

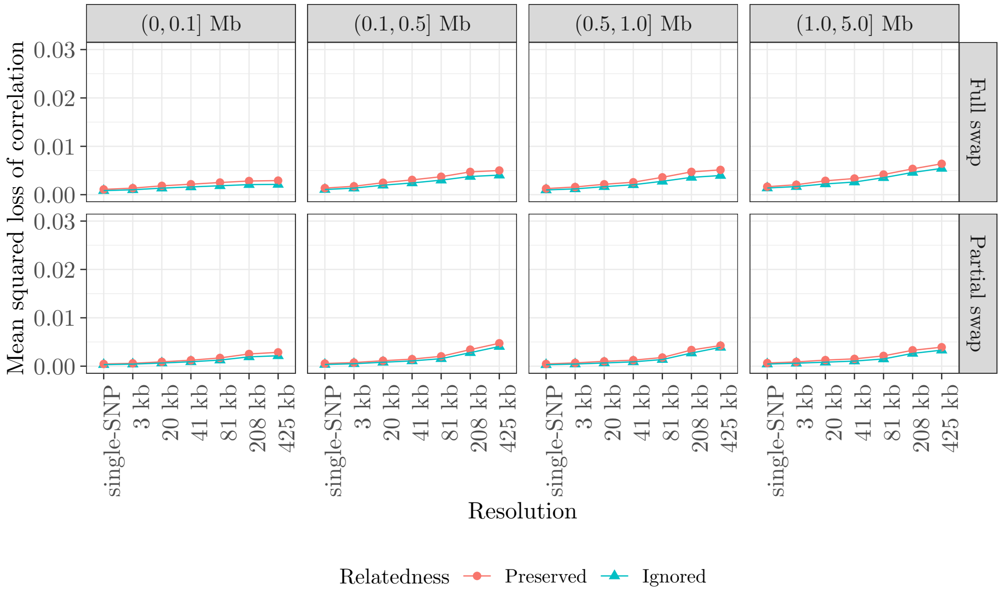


(b) Linkage disequilibrium (smaller values are preferable).

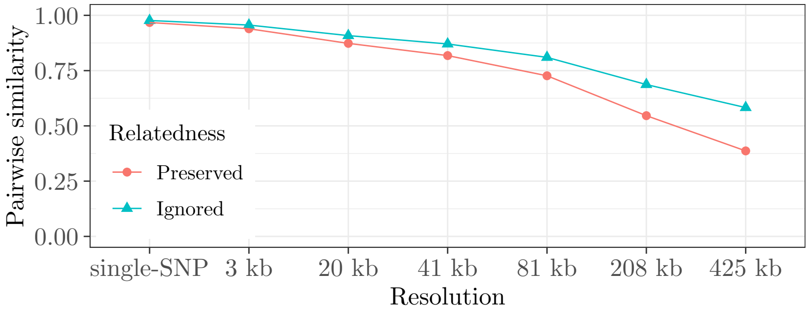


(c) Individual similarity of knockoffs and genotypes (smaller is preferable)

Figure S9: Exchangeability diagnostics for knockoffs on chromosome 22 in 10,000 related British samples from the UK Biobank. The knockoffs are generated with our new method, either preserving or ignoring familial relatedness. The diagnostics are presented as a function of the knockoff resolution. (a) Average width of IBD segments between self-reportedly related individuals, computed on either the real data or the knockoffs. (b) Exchangeability measured in terms of pairwise correlations between different SNPs, as in Figure S8. (c) Absolute pairwise correlation between genotypes and knockoffs, as in Figure S3.

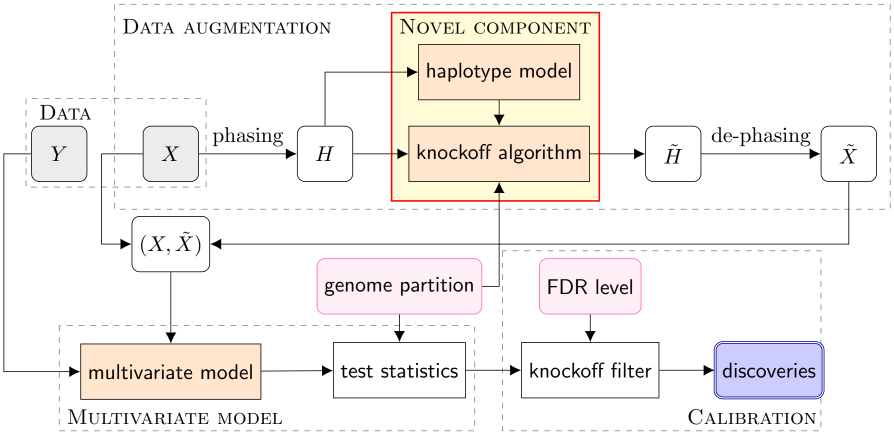


Figure S10: KnockoffGWAS workflow. The novelty introduced in this paper consists of an HMM for the distribution of haplotypes, $H$, that can account for population structure and familial relatedness as well as LD, and of the associated algorithm for generating knockoffs. For computational reasons, the genotypes are phased prior to generating knockoffs, and the knockoff haplotypes are then de-phased to obtain knockoff genotypes [6].

(a) Heritability 0.075

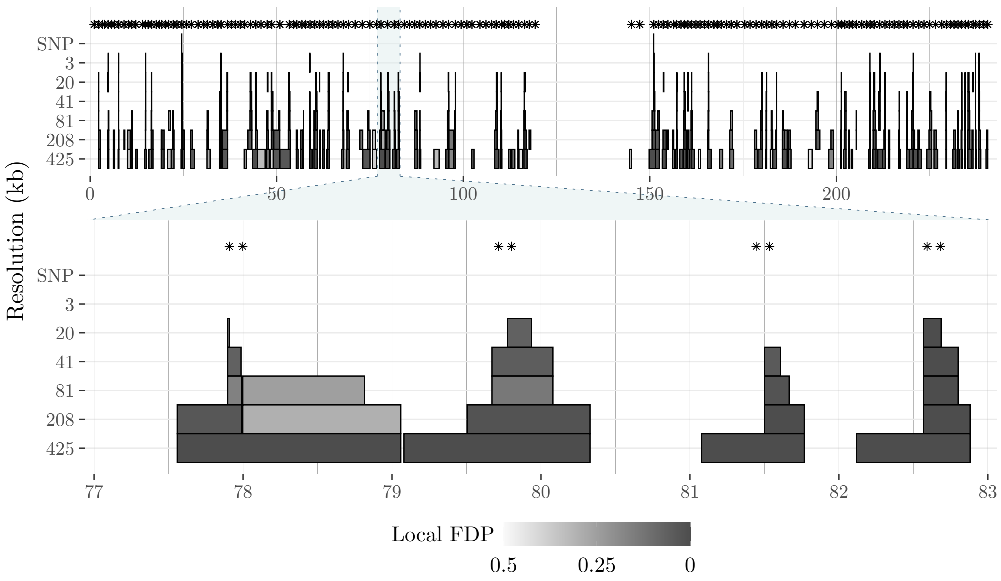


(b) Heritability 0.7

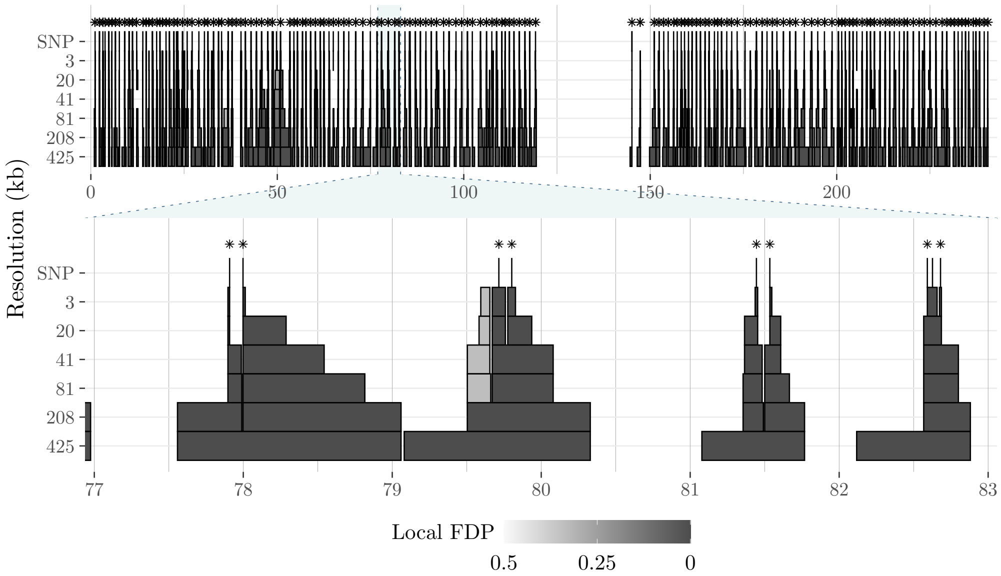


Figure S11: Chicago plots showing KnockoffGWAS discoveries for simulated traits with different levels of heritability, as in Figure 3. Each Chicago plot shows all discoveries on chromosome 1 at the top, and zooms in on a smaller genetic segment at the bottom. The asterisks indicate the position of the causal variants. Note that some "floating discoveries" are visible in the top panel; these can be explicitly avoided with a variation of the knockoff filter that simultaneously processes the results from different resolutions [6]. (a) High heritability (strong signals). (b) Low heritability (weak signals).

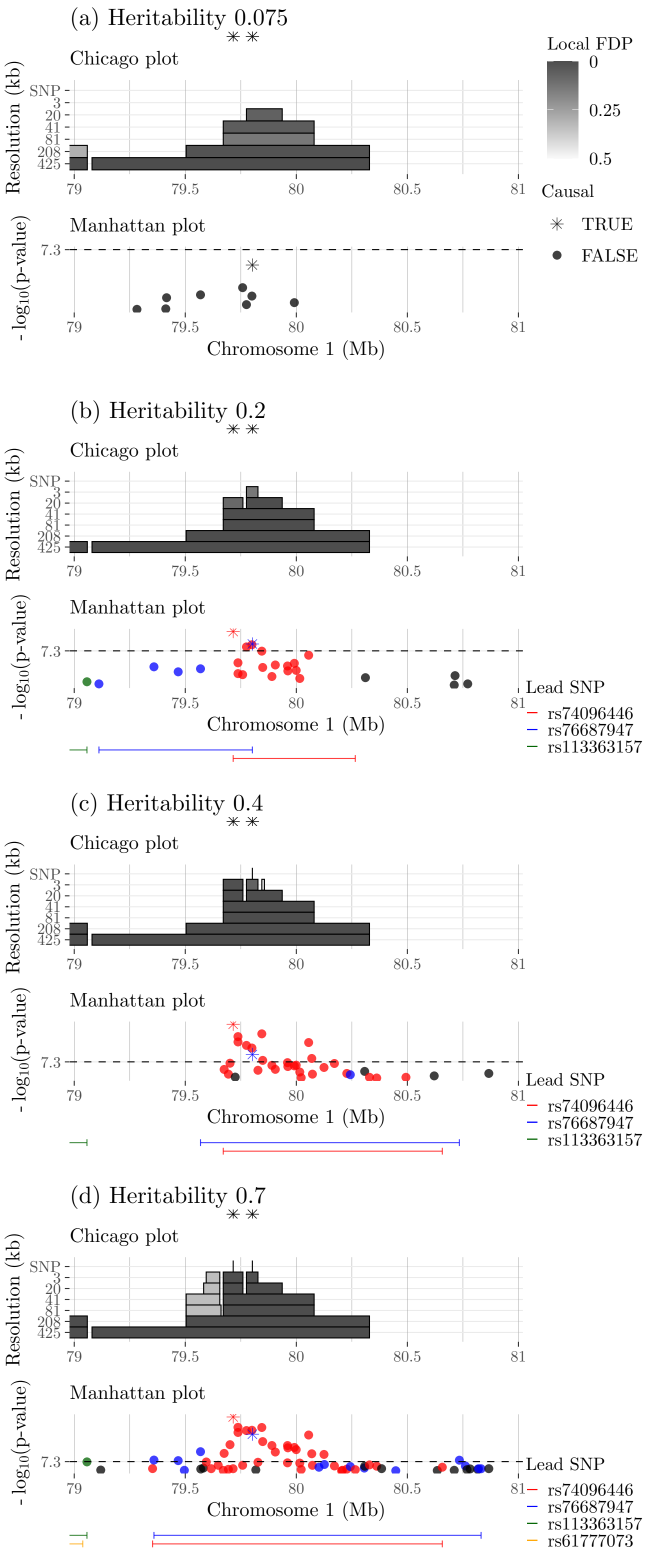


Figure S12: KnockoffGWAS and BOLT-LMM discoveries for simulated traits with different levels of heritability, as in Figure 3.

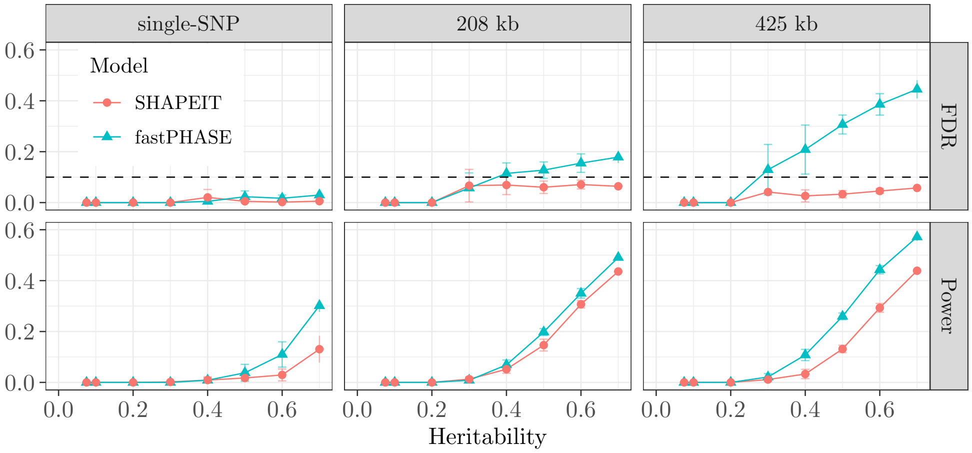


(a) Simulated phenotype with 500 causal variants. Results at three fixed levels of resolution.

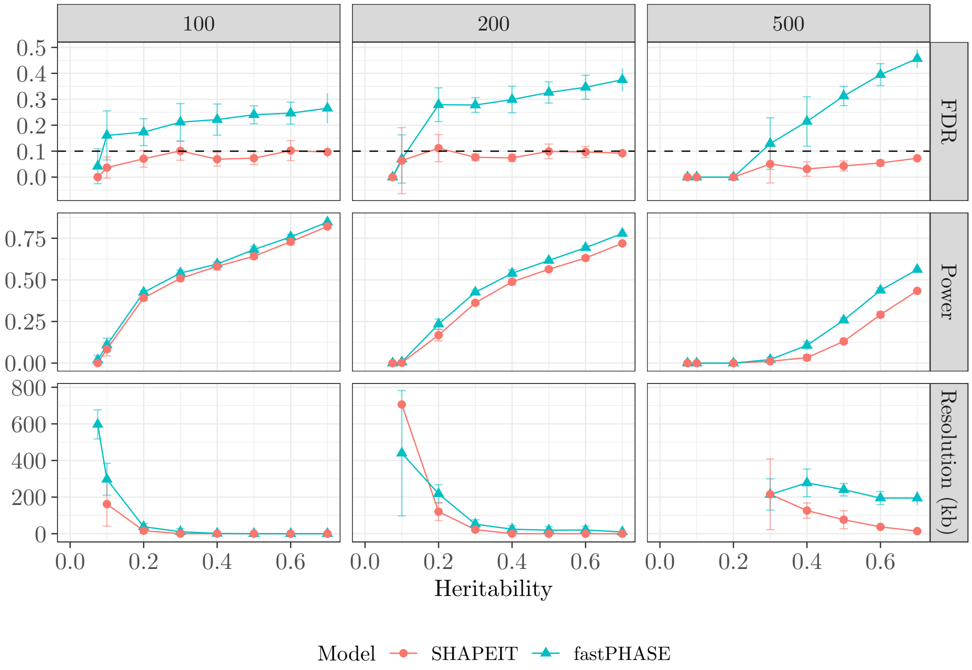


(b) Simulated phenotypes with different numbers of causal variants. Summary of results at different levels of resolution.

Figure S13: KnockoffGWAS performance in numerical experiment with real genotypes of 10,000 individuals with diverse ancestries, as in Figure 1, and simulated phenotypes. The knockoffs are generated either using the SHAPEIT or the fastPHASE [6] HMM. (a) Simulated phenotype with 500 causal variants equally spaced across the genome. (b) Simulated phenotypes with different numbers (100,200, or 500) causal variants equally spaced across the genome. Here, the discoveries at different resolutions are combined, counting only the most specific findings in each locus (this facilitates the visualization, but it is not guaranteed to control the FDR in theory [6]). Other details are as in Figure 4.

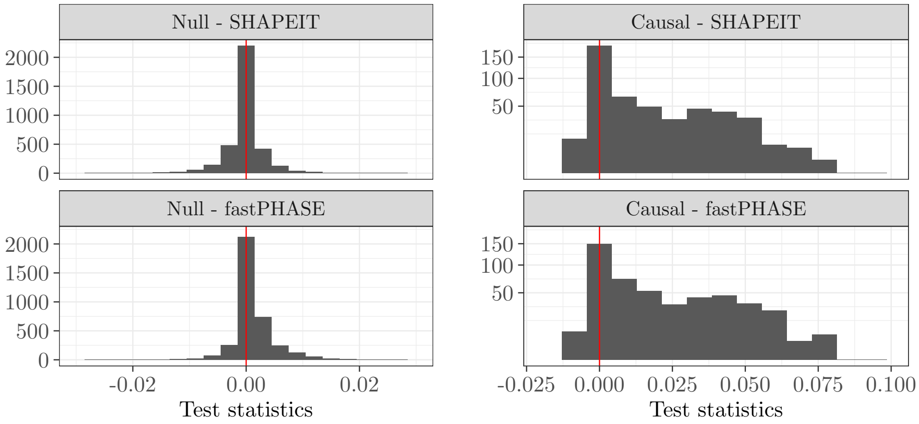


(a) Lasso-based KnockoffGWAS statistics.

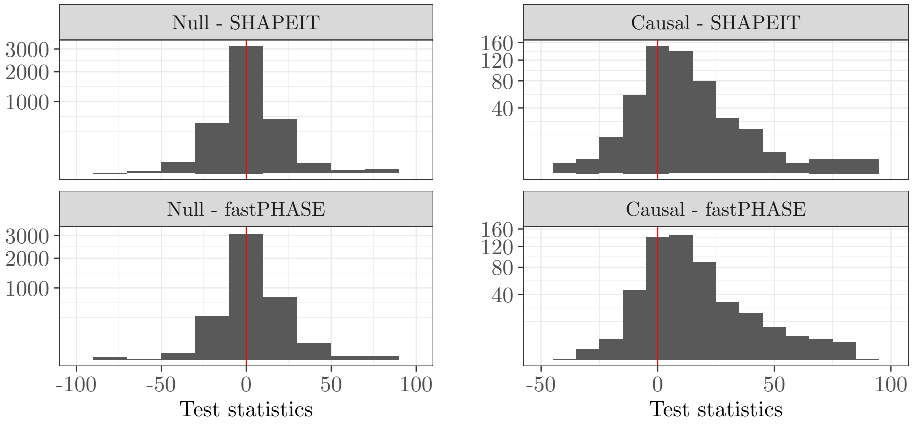


(b) LMM-based KnockoffGWAS statistics.

Figure S14: Distribution of knockoff statistics in the numerical experiment of Figure S13. The knockoffs are constructed by different algorithms at resolution equal to 425 kb. (a) Lasso-based knockoff test statistics for null (left) and causal (right) groups of variants. (b) LMM-based knockoff test statistics for null (left) and causal (right) groups of variants.

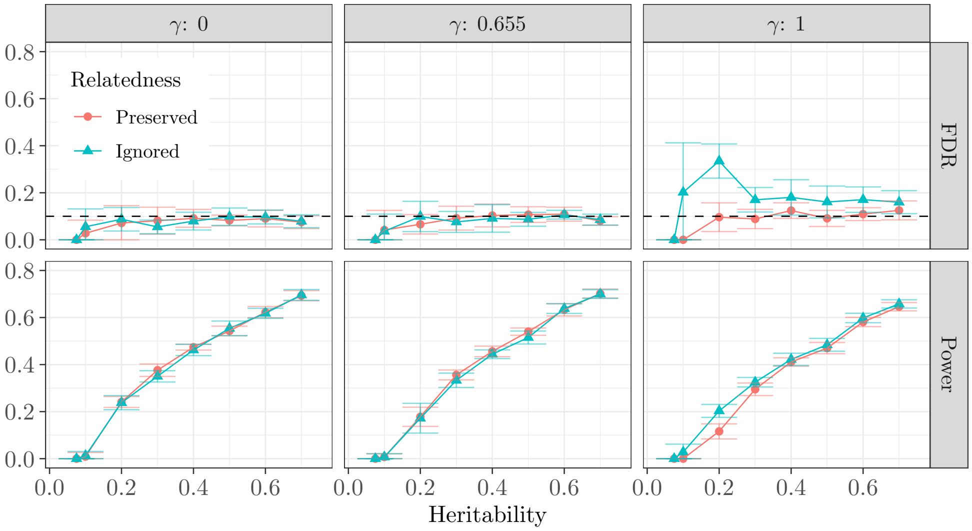


(a) KnockoffGWAS with Lasso-based statistics.

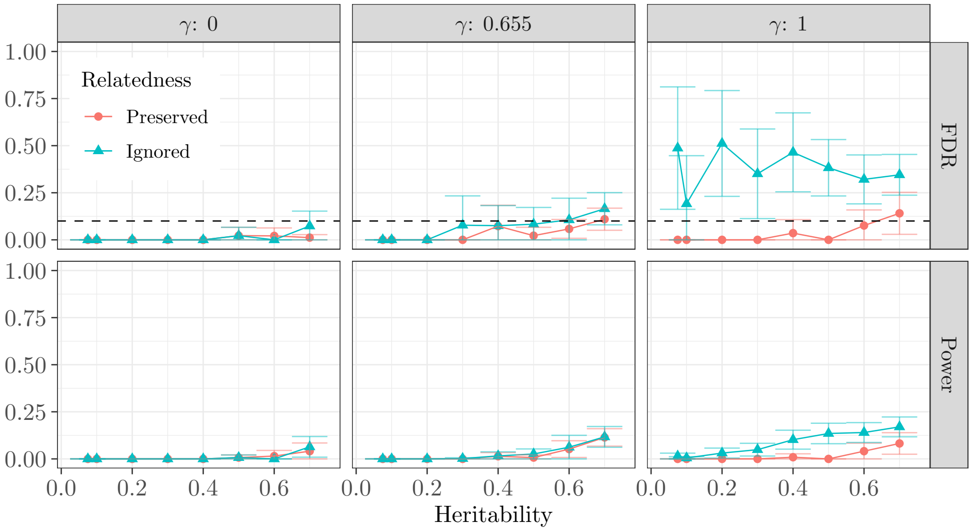


(b) KnockoffGWAS with LMM-based statistics.

Figure S15: Power and FDR in numerical experiments with real genotypes of 10,000 related British samples, and simulated phenotypes. Our method is applied with and without preserving IBD segments. Results for phenotypes with different strengths of family factors $\gamma$ are in separate columns ($\gamma = 0$: no family factors, $\gamma = 1$: strongest family factors; see Methods for more information about $\gamma$). Knockoff resolution equal to 425 kb. (a) Lasso-based test statistics. (b) Marginal test statistics. Note that marginal statistics have almost no power, although an excess of false discoveries occurs if the relatedness is not preserved.

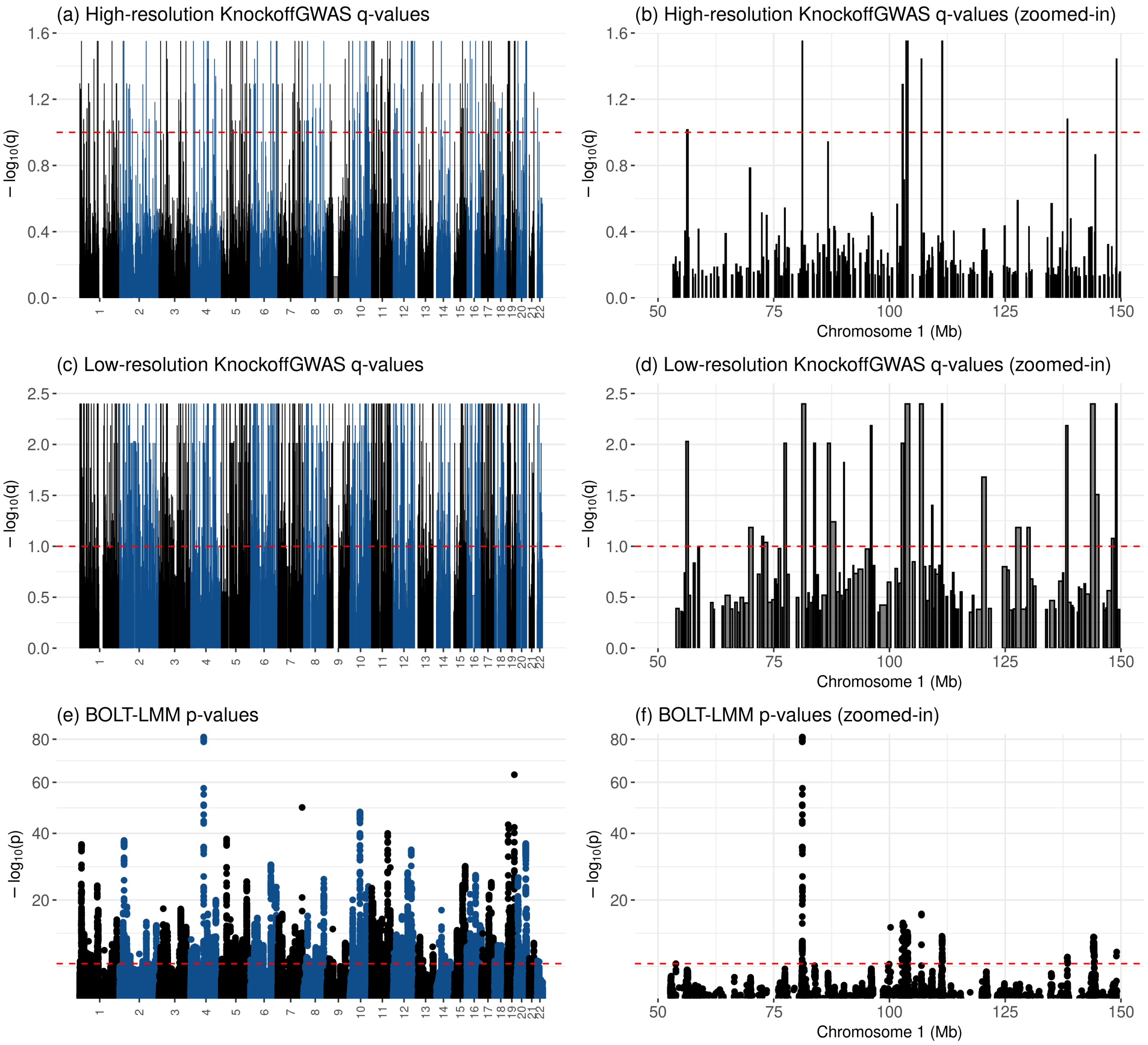


Figure S16: Manhattan plots for cardiovascular disease using the UK Biobank data. (a-b) KnockoffGWAS q-values for high-resolution conditional hypotheses (20 kb). The width of each rectangle denotes the genetic segment tested by the corresponding conditional hypothesis, while the height is the negative logarithm of the q-value. (c-d) KnockoffGWAS q-values for low-resolution conditional hypotheses (208 kb). (e-f) BOLT-LMM p-values for SNP-by-SNP marginal hypotheses. The plots in (c,d,f) are the same as those in (a,c,e), respectively, but zoom in on a portion of chromosome 1. The dashed horizontal red lines indicate the significance thresholds; 10% FDR for KnockoffGWAS, and $5 \times 10^{-8}$ for BOLT-LMM.

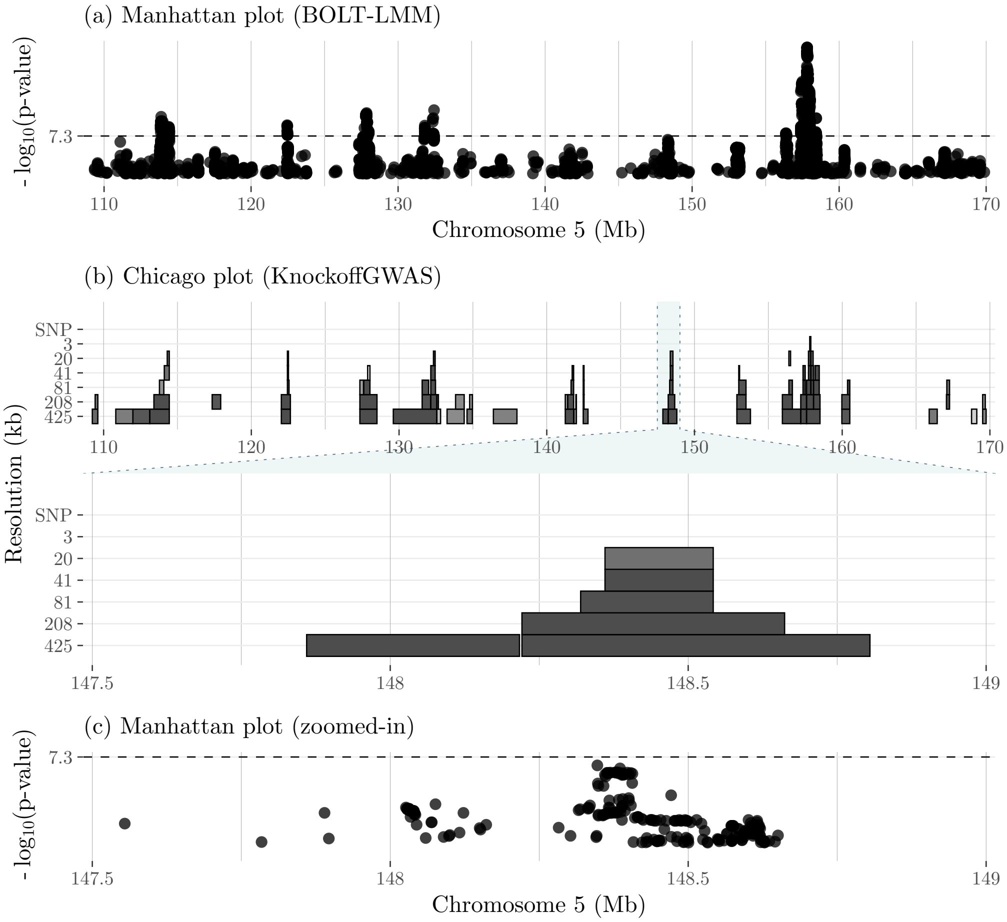


Figure S17: Visualization of some findings for cardiovascular disease on chromosome 5. The top panels visualize a wider portion of chromosome 5, as in Figure S11. Other details are as in Figure 5.

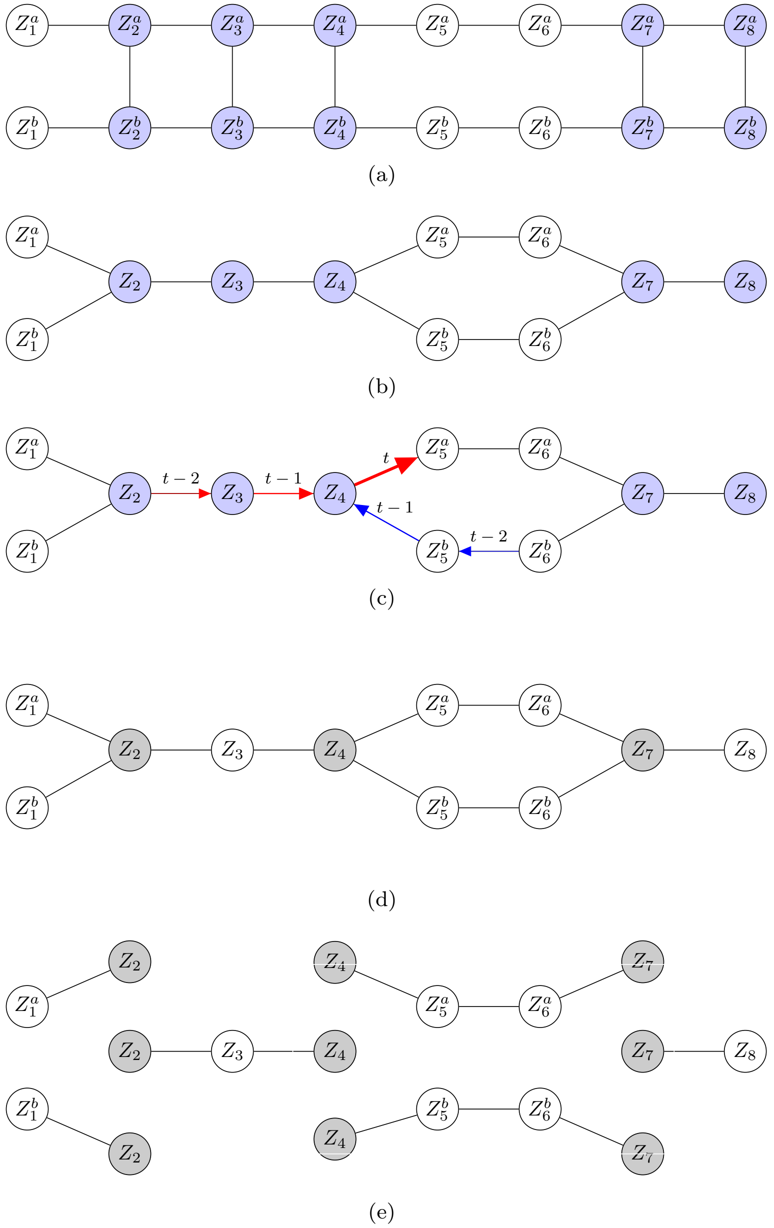


Figure S18: Graphical representation of the distribution of latent states in the generalized HMM for two haplotype sequences of length 8 sharing 2 IBD segments. (a) Representation as a Markov chain with $K^2$ possible states in each position, with the constraint that nodes connected by a vertical edge must be identical to each other. The nodes belonging to the IBD segments are shaded in blue. (b) Equivalent representation of this model as a Markov random field with 11 variables, each taking one of $K$ possible values. (c) Visualization of belief propagation for haplotype families. Belief propagation update of a message in the example of Figure S18. The new message evaluated here at time $t$ is that from the third node of the first shared IBD segment to the successive node in the first haplotype sequence (bold arrow). This is computed as a function of the messages labeled as $t-1$, which had previously been computed as a function of those labeled as $t-2$. Red: forward messages; blue: backward messages. (d) The nodes at the extremities of the IBD segments are shaded in grey. These represent the variables upon which we condition before generating knockoffs. (e) Conditional on the extremities of the IBD segments, the remaining latent nodes are distributed as independent Markov chains.

### S4 Supplementary Tables

| Ethnicity | Count |
|---|---|
| African | 1710 |
| British | 1710 |
| Caribbean | 1710 |
| Chinese | 1450 |
| Indian | 1710 |
| Irish | 1710 |

Table S1: Summary of the self-reported ethnicities for the 10,000 UK Biobank individuals in Figure 1 (a).

| Median width (kb) | Mean width (kb) | Number of groups | Median size (SNPs) | Mean size (SNPs) |
|---|---|---|---|---|
| single-SNP | single-SNP | 591513 | 1 | 1 |
| 3 | 11 | 151532 | 3 | 4 |
| 20 | 41 | 56562 | 8 | 10 |
| 41 | 74 | 33929 | 14 | 17 |
| 81 | 134 | 19500 | 26 | 30 |
| 208 | 303 | 8863 | 58 | 67 |
| 425 | 575 | 4738 | 113 | 125 |

Table S2: Summary of 7 partitions of the genome into disjoint groups of contiguous SNPs. The first column (median width in kb) is used to reference particular resolutions throughout this paper.

| Family size | Number of families | Average kinship |
|---|---|---|
| 1 | 1 | N.A. |
| 2 | 4702 | 0.273 |
| 3 | 193 | 0.270 |
| 4 | 4 | 0.265 |

Table S3: Self-reported family structure for the 10,000 British individuals used in the experiments of Figures S9 and S13. These families are chosen as those with the largest average kinship. One extra individual is included to bring the total number to a round value.

| Name | Description | Number of cases | UK Biobank Fields | UK Biobank Codes |
|---|---|---|---|---|
| bmi | body mass index | continuous | 21001-0.0 | |
| cvd | cardiovascular disease | 148715 | 20002-0.0–20002-0.32 | 1065, 1066, 1067, 1068, 1081, 1082, 1083, 1425, 1473, 1493 |
| diabetes | diabetes | 19897 | 20002-0.0–20002-0.32 | 1220 |
| height | standing height | continuous | 50-0.0 | |
| hypothyroidism | hypothyroidism | 22493 | 20002-0.0–20002-0.32 | 1226 |
| platelet | platelet count | continuous | 30080-0.0 | |
| respiratory | respiratory disease | 64945 | 20002-0.0–20002-0.32 | 1111, 1112, 1113, 1114, 1115, 1117, 1413, 1414, 1415, 1594 |
| sbp | systolic blood pressure | continuous | 4080-0.0, 4080-0.1 | |

Table S4: Definition of the UK Biobank phenotypes used in our analysis [6]. In the case of case-control phenotypes, the number of cases refers to the subset of individuals that passed our quality control.

| | KnockoffGWAS | | | BOLT-LMM | |
|---|---|---|---|---|---|
| Phenotype | Resolution | Discoveries | Overlap with LMM | Discoveries | Overlap with KZ |
| cvd | 3 kb | 22 | 22 (100.0%) | 257 | 25 (9.7%) |
| | 20 kb | 239 | 180 (75.3%) | 257 | 189 (73.5%) |
| | 41 kb | 339 | 212 (62.5%) | 257 | 213 (82.9%) |
| | 81 kb | 566 | 261 (46.1%) | 257 | 241 (93.8%) |
| | 208 kb | 940 | 274 (29.1%) | 257 | 249 (96.9%) |
| | 425 kb | 1089 | 255 (23.4%) | 257 | 254 (98.8%) |
| diabetes | 3 kb | 21 | 20 (95.2%) | 62 | 21 (33.9%) |
| | 20 kb | 61 | 45 (73.8%) | 62 | 47 (75.8%) |
| | 41 kb | 109 | 54 (49.5%) | 62 | 52 (83.9%) |
| | 81 kb | 109 | 50 (45.9%) | 62 | 54 (87.1%) |
| | 208 kb | 113 | 52 (46.0%) | 62 | 55 (88.7%) |
| | 425 kb | 194 | 57 (29.4%) | 62 | 59 (95.2%) |
| hypothyroidism | single-SNP | 19 | 19 (100.0%) | 143 | 30 (21.0%) |
| | 3 kb | 40 | 40 (100.0%) | 143 | 53 (37.1%) |
| | 20 kb | 105 | 89 (84.8%) | 143 | 101 (70.6%) |
| | 41 kb | 222 | 128 (57.7%) | 143 | 123 (86.0%) |
| | 81 kb | 277 | 133 (48.0%) | 143 | 130 (90.9%) |
| | 208 kb | 295 | 129 (43.7%) | 143 | 142 (99.3%) |
| | 425 kb | 335 | 122 (36.4%) | 143 | 142 (99.3%) |
| respiratory | 20 kb | 83 | 60 (72.3%) | 94 | 62 (66.0%) |
| | 41 kb | 123 | 74 (60.2%) | 94 | 75 (79.8%) |
| | 81 kb | 193 | 83 (43.0%) | 94 | 85 (90.4%) |
| | 208 kb | 262 | 82 (31.3%) | 94 | 92 (97.9%) |
| | 425 kb | 383 | 82 (21.4%) | 94 | 93 (98.9%) |
| bmi | 3 kb | 10 | 10 (100.0%) | 697 | 15 (2.2%) |
| | 20 kb | 343 | 309 (90.1%) | 697 | 317 (45.5%) |
| | 41 kb | 918 | 618 (67.3%) | 697 | 548 (78.6%) |
| | 81 kb | 1480 | 792 (53.5%) | 697 | 641 (92.0%) |
| | 208 kb | 2395 | 898 (37.5%) | 697 | 689 (98.9%) |
| | 425 kb | 2460 | 794 (32.3%) | 697 | 695 (99.7%) |
| height | single-SNP | 95 | 95 (100.0%) | 2464 | 225 (9.1%) |
| | 3 kb | 570 | 570 (100.0%) | 2464 | 891 (36.2%) |
| | 20 kb | 1503 | 1469 (97.7%) | 2464 | 1761 (71.5%) |
| | 41 kb | 2384 | 2167 (90.9%) | 2464 | 2167 (87.9%) |
| | 81 kb | 3006 | 2417 (80.4%) | 2464 | 2360 (95.8%) |
| | 208 kb | 3339 | 2228 (66.7%) | 2464 | 2430 (98.6%) |
| | 425 kb | 3073 | 1804 (58.7%) | 2464 | 2454 (99.6%) |
| platelet | single-SNP | 53 | 53 (100.0%) | 1204 | 131 (10.9%) |
| | 3 kb | 246 | 245 (99.6%) | 1204 | 391 (32.5%) |
| | 20 kb | 1002 | 900 (89.8%) | 1204 | 963 (80.0%) |
| | 41 kb | 1261 | 1041 (82.6%) | 1204 | 1075 (89.3%) |
| | 81 kb | 1570 | 1120 (71.3%) | 1204 | 1138 (94.5%) |
| | 208 kb | 1743 | 1057 (60.6%) | 1204 | 1183 (98.3%) |
| | 425 kb | 1653 | 911 (55.1%) | 1204 | 1195 (99.3%) |
| sbp | 3 kb | 83 | 83 (100.0%) | 568 | 101 (17.8%) |
| | 20 kb | 191 | 177 (92.7%) | 568 | 204 (35.9%) |
| | 41 kb | 511 | 366 (71.6%) | 568 | 380 (66.9%) |
| | 81 kb | 830 | 496 (59.8%) | 568 | 480 (84.5%) |
| | 208 kb | 1183 | 561 (47.4%) | 568 | 530 (93.3%) |
| | 425 kb | 1543 | 538 (34.9%) | 568 | 548 (96.5%) |

Table S5: KnockoffGWAS discoveries for different phenotypes using all UK Biobank samples vs. BOLT-LMM genome-wide significant discoveries ($5 \times 10^{-8}$). BOLT-LMM is applied on 350k unrelated British samples for diabetes and respiratory disease [6], and on 459k European samples for all other phenotypes [8].

| Phenotype | Resolution | | Discoveries with SHAPEIT model | | Discoveries with fastPHASE model | |
|---|---|---|---|---|---|---|
| | SHAPEIT | fastPHASE | Number | Overlap with fastPHASE | Number | Overlap with SHAPEIT |
| cvd | 41 kb | 42 kb | 339 | 49 (14.5%) | 51 | 46 (90.2%) |
| | 81 kb | 88 kb | 566 | 175 (30.9%) | 182 | 165 (90.7%) |
| | 208 kb | 226 kb | 940 | 449 (47.8%) | 514 | 446 (86.8%) |
| | 425 kb | 226 kb | 1089 | 453 (41.6%) | 514 | 466 (90.7%) |
| diabetes | 3 kb | 4 kb | 21 | 8 (38.1%) | 11 | 8 (72.7%) |
| | 20 kb | 18 kb | 61 | 9 (14.8%) | 10 | 9 (90.0%) |
| | 41 kb | 42 kb | 109 | 19 (17.4%) | 21 | 19 (90.5%) |
| | 81 kb | 88 kb | 109 | 28 (25.7%) | 33 | 28 (84.8%) |
| | 208 kb | 226 kb | 113 | 45 (39.8%) | 50 | 46 (92.0%) |
| | 425 kb | 226 kb | 194 | 48 (24.7%) | 50 | 48 (96.0%) |
| hypothyroidism | single-SNP | single-SNP | 19 | 8 (42.1%) | 21 | 8 (38.1%) |
| | 81 kb | 88 kb | 277 | 103 (37.2%) | 108 | 100 (92.6%) |
| | 208 kb | 226 kb | 295 | 183 (62.0%) | 212 | 186 (87.7%) |
| | 425 kb | 226 kb | 335 | 188 (56.1%) | 212 | 194 (91.5%) |
| respiratory | | | | | | |
| | 20 kb | 18 kb | 83 | 12 (14.5%) | 13 | 13 (100.0%) |
| | 41 kb | 42 kb | 123 | 35 (28.5%) | 41 | 35 (85.4%) |
| | 81 kb | 88 kb | 193 | 61 (31.6%) | 65 | 59 (90.8%) |
| | 208 kb | 226 kb | 262 | 132 (50.4%) | 176 | 140 (79.5%) |
| | 425 kb | 226 kb | 383 | 154 (40.2%) | 176 | 159 (90.3%) |
| bmi | 3 kb | 4 kb | 10 | 7 (70.0%) | 24 | 7 (29.2%) |
| | 20 kb | 18 kb | 343 | 29 (8.5%) | 33 | 30 (90.9%) |
| | 41 kb | 42 kb | 918 | 61 (6.6%) | 60 | 58 (96.7%) |
| | 81 kb | 88 kb | 1480 | 515 (34.8%) | 555 | 485 (87.4%) |
| | 208 kb | 226 kb | 2395 | 1653 (69.0%) | 1804 | 1615 (89.5%) |
| | 425 kb | 226 kb | 2460 | 1592 (64.7%) | 1804 | 1733 (96.1%) |
| height | single-SNP | single-SNP | 95 | 68 (71.6%) | 173 | 68 (39.3%) |
| | 3 kb | 4 kb | 570 | 252 (44.2%) | 336 | 251 (74.7%) |
| | 20 kb | 18 kb | 1503 | 360 (24.0%) | 388 | 350 (90.2%) |
| | 41 kb | 42 kb | 2384 | 832 (34.9%) | 823 | 780 (94.8%) |
| | 81 kb | 88 kb | 3006 | 1864 (62.0%) | 1976 | 1836 (92.9%) |
| | 208 kb | 226 kb | 3339 | 2775 (83.1%) | 3284 | 3021 (92.0%) |
| | 425 kb | 226 kb | 3073 | 2398 (78.0%) | 3284 | 3198 (97.4%) |
| platelet | single-SNP | single-SNP | 53 | 40 (75.5%) | 143 | 40 (28.0%) |
| | 3 kb | 4 kb | 246 | 136 (55.3%) | 161 | 138 (85.7%) |
| | 20 kb | 18 kb | 1002 | 264 (26.3%) | 276 | 265 (96.0%) |
| | 41 kb | 42 kb | 1261 | 398 (31.6%) | 408 | 385 (94.4%) |
| | 81 kb | 88 kb | 1570 | 856 (54.5%) | 890 | 834 (93.7%) |
| | 208 kb | 226 kb | 1743 | 1288 (73.9%) | 1460 | 1325 (90.8%) |
| | 425 kb | 226 kb | 1653 | 1162 (70.3%) | 1460 | 1393 (95.4%) |
| sbp | 41 kb | 42 kb | 511 | 86 (16.8%) | 95 | 84 (88.4%) |
| | 81 kb | 88 kb | 830 | 265 (31.9%) | 297 | 262 (88.2%) |
| | 208 kb | 226 kb | 1183 | 619 (52.3%) | 722 | 612 (84.8%) |
| | 425 kb | 226 kb | 1543 | 663 (43.0%) | 722 | 678 (93.9%) |

Table S6: KnockoffGWAS discoveries obtained from all UK Biobank British samples (SHAPEIT model) vs. discoveries obtained from 350k unrelated British samples (fastPHASE model); the latter are based on the slightly different genome partitions adopted by [6].

| | | Everyone | | British | | White (non-British) | |
|---|---|---|---|---|---|---|---|
| Phenotype | Resolution | all | unrel. | all | unrel. | all | unrel. |
| cvd | single-SNP | 0 | 0 | 0 | 0 | 0 | 0 |
| | 3 kb | 22 | 20 | 0 | 0 | 0 | 0 |
| | 20 kb | 239 | 152 | 169 | 140 | 0 | 0 |
| | 41 kb | 339 | 235 | 270 | 181 | 0 | 0 |
| | 81 kb | 566 | 428 | 611 | 462 | 0 | 0 |
| | 208 kb | 940 | 594 | 815 | 611 | 0 | 0 |
| | 425 kb | 1089 | 861 | 1004 | 711 | 0 | 0 |
| diabetes | single-SNP | 0 | 0 | 0 | 0 | 0 | 0 |
| | 3 kb | 0 | 17 | 0 | 12 | 0 | 0 |
| | 20 kb | 83 | 44 | 63 | 53 | 0 | 0 |
| | 41 kb | 123 | 86 | 82 | 57 | 0 | 0 |
| | 81 kb | 193 | 152 | 165 | 129 | 0 | 0 |
| | 208 kb | 262 | 242 | 217 | 171 | 0 | 0 |
| | 425 kb | 383 | 346 | 289 | 291 | 0 | 0 |
| hypothyroidism | single-SNP | 19 | 22 | 11 | 11 | 0 | 0 |
| | 3 kb | 40 | 42 | 60 | 32 | 0 | 0 |
| | 20 kb | 105 | 79 | 109 | 86 | 0 | 0 |
| | 41 kb | 222 | 156 | 164 | 130 | 0 | 0 |
| | 81 kb | 277 | 173 | 269 | 153 | 0 | 0 |
| | 208 kb | 295 | 257 | 288 | 256 | 0 | 0 |
| | 425 kb | 335 | 309 | 312 | 266 | 0 | 0 |
| respiratory | single-SNP | 0 | 0 | 0 | 11 | 0 | 0 |
| | 3 kb | 21 | 0 | 37 | 0 | 0 | 0 |
| | 20 kb | 61 | 33 | 54 | 33 | 0 | 0 |
| | 41 kb | 109 | 62 | 73 | 66 | 0 | 0 |
| | 81 kb | 109 | 84 | 63 | 94 | 0 | 0 |
| | 208 kb | 113 | 79 | 119 | 84 | 0 | 0 |
| | 425 kb | 194 | 102 | 186 | 139 | 0 | 0 |
| bmi | single-SNP | 95 | 64 | 80 | 69 | 0 | 10 |
| | 3 kb | 570 | 483 | 609 | 377 | 0 | 0 |
| | 20 kb | 1503 | 1294 | 1610 | 1412 | 25 | 20 |
| | 41 kb | 2384 | 1966 | 2353 | 2141 | 74 | 80 |
| | 81 kb | 3006 | 2768 | 3002 | 2681 | 91 | 80 |
| | 208 kb | 3339 | 3111 | 3370 | 3117 | 112 | 101 |
| | 425 kb | 3073 | 2922 | 2938 | 2735 | 170 | 104 |
| height | single-SNP | 0 | 0 | 0 | 12 | 0 | 0 |
| | 3 kb | 10 | 10 | 0 | 0 | 0 | 0 |
| | 20 kb | 343 | 230 | 207 | 180 | 0 | 0 |
| | 41 kb | 918 | 566 | 820 | 492 | 0 | 0 |
| | 81 kb | 1480 | 1194 | 1433 | 1280 | 0 | 0 |
| | 208 kb | 2395 | 1938 | 2381 | 1975 | 0 | 0 |
| | 425 kb | 2460 | 2109 | 2426 | 2092 | 0 | 10 |
| platelet | single-SNP | 53 | 52 | 34 | 52 | 0 | 0 |
| | 3 kb | 246 | 259 | 223 | 202 | 0 | 0 |
| | 20 kb | 1002 | 820 | 977 | 777 | 26 | 31 |
| | 41 kb | 1261 | 995 | 1171 | 944 | 52 | 44 |
| | 81 kb | 1570 | 1350 | 1502 | 1292 | 69 | 55 |
| | 208 kb | 1743 | 1583 | 1809 | 1510 | 53 | 51 |
| | 425 kb | 1653 | 1550 | 1741 | 1521 | 76 | 60 |
| sbp | single-SNP | 0 | 0 | 0 | 0 | 0 | 0 |
| | 3 kb | 83 | 90 | 42 | 32 | 0 | 0 |
| | 20 kb | 191 | 162 | 166 | 127 | 0 | 0 |
| | 41 kb | 511 | 353 | 421 | 342 | 0 | 0 |
| | 81 kb | 830 | 635 | 736 | 585 | 0 | 0 |
| | 208 kb | 1183 | 972 | 1050 | 911 | 0 | 0 |
| | 425 kb | 1543 | 1202 | 1401 | 1273 | 0 | 0 |

Table S7: Numbers of KnockoffGWAS discoveries at different resolutions, using different subsets of the UK Biobank samples.

| | | | Confirmed | | | |
|---|---|---|---|---|---|---|
| Phenotype | Resolution | Discoveries | Catalog | Japan | FinnGen | Any |
| cvd | 3 kb | 22 | 21 (95.5%) | NA | 11 (50.0%) | 22 (100.0%) |
| | 20 kb | 239 | 173 (72.4%) | NA | 81 (33.9%) | 188 (78.7%) |
| | 41 kb | 339 | 241 (71.1%) | NA | 126 (37.2%) | 266 (78.5%) |
| | 81 kb | 566 | 353 (62.4%) | NA | 251 (44.3%) | 422 (74.6%) |
| | 208 kb | 940 | 524 (55.7%) | NA | 581 (61.8%) | 738 (78.5%) |
| | 425 kb | 1089 | 671 (61.6%) | NA | 837 (76.9%) | 967 (88.8%) |
| diabetes | 3 kb | 21 | 20 (95.2%) | 13 (61.9%) | 8 (38.1%) | 20 (95.2%) |
| | 20 kb | 61 | 54 (88.5%) | 26 (42.6%) | 18 (29.5%) | 54 (88.5%) |
| | 41 kb | 109 | 88 (80.7%) | 36 (33.0%) | 30 (27.5%) | 88 (80.7%) |
| | 81 kb | 109 | 88 (80.7%) | 39 (35.8%) | 36 (33.0%) | 89 (81.7%) |
| | 208 kb | 113 | 95 (84.1%) | 49 (43.4%) | 43 (38.1%) | 97 (85.8%) |
| | 425 kb | 194 | 140 (72.2%) | 58 (29.9%) | 59 (30.4%) | 142 (73.2%) |
| hypothyroidism | single-SNP | 19 | 7 (36.8%) | NA | 3 (15.8%) | 7 (36.8%) |
| | 3 kb | 40 | 23 (57.5%) | NA | 14 (35.0%) | 24 (60.0%) |
| | 20 kb | 105 | 71 (67.6%) | NA | 20 (19.0%) | 71 (67.6%) |
| | 41 kb | 222 | 101 (45.5%) | NA | 27 (12.2%) | 105 (47.3%) |
| | 81 kb | 277 | 126 (45.5%) | NA | 38 (13.7%) | 135 (48.7%) |
| | 208 kb | 295 | 141 (47.8%) | NA | 50 (16.9%) | 156 (52.9%) |
| | 425 kb | 335 | 139 (41.5%) | NA | 74 (22.1%) | 174 (51.9%) |
| respiratory | 20 kb | 83 | 74 (89.2%) | NA | 35 (42.2%) | 76 (91.6%) |
| | 41 kb | 123 | 110 (89.4%) | NA | 58 (47.2%) | 114 (92.7%) |
| | 81 kb | 193 | 155 (80.3%) | NA | 115 (59.6%) | 174 (90.2%) |
| | 208 kb | 262 | 195 (74.4%) | NA | 202 (77.1%) | 241 (92.0%) |
| | 425 kb | 383 | 263 (68.7%) | NA | 330 (86.2%) | 357 (93.2%) |
| bmi | 3 kb | 10 | 10 (100.0%) | 4 (40.0%) | NA | 10 (100.0%) |
| | 20 kb | 343 | 307 (89.5%) | 32 (9.3%) | NA | 308 (89.8%) |
| | 41 kb | 918 | 655 (71.4%) | 53 (5.8%) | NA | 656 (71.5%) |
| | 81 kb | 1480 | 865 (58.4%) | 55 (3.7%) | NA | 865 (58.4%) |
| | 208 kb | 2395 | 1076 (44.9%) | 64 (2.7%) | NA | 1076 (44.9%) |
| | 425 kb | 2460 | 1090 (44.3%) | 68 (2.8%) | NA | 1091 (44.3%) |
| height | single-SNP | 95 | 63 (66.3%) | 57 (60.0%) | NA | 81 (85.3%) |
| | 3 kb | 570 | 357 (62.6%) | 258 (45.3%) | NA | 417 (73.2%) |
| | 20 kb | 1503 | 1032 (68.7%) | 483 (32.1%) | NA | 1102 (73.3%) |
| | 41 kb | 2384 | 1534 (64.3%) | 572 (24.0%) | NA | 1607 (67.4%) |
| | 81 kb | 3006 | 1822 (60.6%) | 590 (19.6%) | NA | 1879 (62.5%) |
| | 208 kb | 3339 | 1856 (55.6%) | 561 (16.8%) | NA | 1886 (56.5%) |
| | 425 kb | 3073 | 1653 (53.8%) | 494 (16.1%) | NA | 1669 (54.3%) |
| platelet | single-SNP | 53 | 37 (69.8%) | 22 (41.5%) | NA | 41 (77.4%) |
| | 3 kb | 246 | 153 (62.2%) | 72 (29.3%) | NA | 168 (68.3%) |
| | 20 kb | 1002 | 352 (35.1%) | 97 (9.7%) | NA | 374 (37.3%) |
| | 41 kb | 1261 | 391 (31.0%) | 97 (7.7%) | NA | 409 (32.4%) |
| | 81 kb | 1570 | 426 (27.1%) | 91 (5.8%) | NA | 436 (27.8%) |
| | 208 kb | 1743 | 445 (25.5%) | 94 (5.4%) | NA | 453 (26.0%) |
| | 425 kb | 1653 | 425 (25.7%) | 86 (5.2%) | NA | 429 (26.0%) |
| sbp | 3 kb | 83 | 69 (83.1%) | 10 (12.0%) | NA | 69 (83.1%) |
| | 20 kb | 191 | 166 (86.9%) | 17 (8.9%) | NA | 166 (86.9%) |
| | 41 kb | 511 | 358 (70.1%) | 22 (4.3%) | NA | 359 (70.3%) |
| | 81 kb | 830 | 517 (62.3%) | 22 (2.7%) | NA | 517 (62.3%) |
| | 208 kb | 1183 | 643 (54.4%) | 23 (1.9%) | NA | 643 (54.4%) |
| | 425 kb | 1543 | 709 (45.9%) | 23 (1.5%) | NA | 709 (45.9%) |

Table S8: Numbers of KnockoffGWAS discoveries at different resolutions (all UK Biobank samples) containing associations previously reported in the GWAS Catalog, Japan Biobank resource, FinnGen resource, or any of the above.

| | | Found by BOLT-LMM | | | | | Not found by BOLT-LMM | | | | |
|---|---|---|---|---|---|---|---|---|---|---|---|
| Phenotype | Resolution | Total | Catalog | Japan | FinnGen | Any | Total | Catalog | Japan | FinnGen | Any |
| cvd | 3 kb | 22 | 95.5% | NA | 50.0% | 100.0% | 0 | NA | NA | NA | NA |
| | 20 kb | 180 | 83.9% | NA | 38.3% | 88.9% | 59 | 37.3% | NA | 20.3% | 47.5% |
| | 41 kb | 212 | 86.8% | NA | 42.9% | 91.5% | 127 | 44.9% | NA | 27.6% | 56.7% |
| | 81 kb | 261 | 87.4% | NA | 53.3% | 92.7% | 305 | 41.0% | NA | 36.7% | 59.0% |
| | 208 kb | 274 | 90.5% | NA | 71.9% | 97.1% | 666 | 41.4% | NA | 57.7% | 70.9% |
| | 425 kb | 255 | 94.9% | NA | 84.7% | 99.2% | 834 | 51.4% | NA | 74.5% | 85.6% |
| diabetes | 3 kb | 20 | 95.0% | 65.0% | 40.0% | 95.0% | 1 | 100.0% | 0.0% | 0.0% | 100.0% |
| | 20 kb | 45 | 95.6% | 53.3% | 40.0% | 95.6% | 16 | 68.8% | 12.5% | 0.0% | 68.8% |
| | 41 kb | 54 | 96.3% | 50.0% | 46.3% | 96.3% | 55 | 65.5% | 16.4% | 9.1% | 65.5% |
| | 81 kb | 50 | 100.0% | 54.0% | 56.0% | 100.0% | 59 | 64.4% | 20.3% | 13.6% | 66.1% |
| | 208 kb | 52 | 98.1% | 61.5% | 61.5% | 98.1% | 61 | 72.1% | 27.9% | 18.0% | 75.4% |
| | 425 kb | 57 | 98.2% | 61.4% | 63.2% | 98.2% | 137 | 61.3% | 16.8% | 16.8% | 62.8% |
| hypothyroidism | single-SNP | 19 | 36.8% | NA | 15.8% | 36.8% | 0 | NA | NA | NA | NA |
| | 3 kb | 40 | 57.5% | NA | 35.0% | 60.0% | 0 | NA | NA | NA | NA |
| | 20 kb | 89 | 76.4% | NA | 22.5% | 76.4% | 16 | 18.8% | NA | 0.0% | 18.8% |
| | 41 kb | 128 | 64.1% | NA | 18.0% | 65.6% | 94 | 20.2% | NA | 4.3% | 22.3% |
| | 81 kb | 133 | 75.2% | NA | 22.6% | 78.9% | 144 | 18.1% | NA | 5.6% | 20.8% |
| | 208 kb | 129 | 85.3% | NA | 25.6% | 87.6% | 166 | 18.7% | NA | 10.2% | 25.9% |
| | 425 kb | 122 | 88.5% | NA | 32.8% | 93.4% | 213 | 14.6% | NA | 16.0% | 28.2% |
| respiratory | 20 kb | 60 | 98.3% | NA | 48.3% | 98.3% | 23 | 65.2% | NA | 26.1% | 73.9% |
| | 41 kb | 74 | 100.0% | NA | 51.4% | 100.0% | 49 | 73.5% | NA | 40.8% | 81.6% |
| | 81 kb | 83 | 98.8% | NA | 65.1% | 100.0% | 110 | 66.4% | NA | 55.5% | 82.7% |
| | 208 kb | 82 | 98.8% | NA | 79.3% | 100.0% | 180 | 63.3% | NA | 76.1% | 88.3% |
| | 425 kb | 82 | 96.3% | NA | 92.7% | 100.0% | 301 | 61.1% | NA | 84.4% | 91.4% |
| bmi | 3 kb | 10 | 100.0% | 40.0% | NA | 100.0% | 0 | NA | NA | NA | NA |
| | 20 kb | 309 | 94.2% | 10.4% | NA | 94.5% | 34 | 47.1% | 0.0% | NA | 47.1% |
| | 41 kb | 618 | 89.3% | 8.3% | NA | 89.5% | 300 | 34.3% | 0.7% | NA | 34.3% |
| | 81 kb | 792 | 85.2% | 6.4% | NA | 85.2% | 688 | 27.6% | 0.6% | NA | 27.6% |
| | 208 kb | 898 | 82.5% | 6.2% | NA | 82.5% | 1497 | 22.4% | 0.5% | NA | 22.4% |
| | 425 kb | 794 | 85.8% | 7.6% | NA | 85.9% | 1666 | 24.5% | 0.5% | NA | 24.5% |
| height | single-SNP | 95 | 66.3% | 60.0% | NA | 85.3% | 0 | NA | NA | NA | NA |
| | 3 kb | 570 | 62.6% | 45.3% | NA | 73.2% | 0 | NA | NA | NA | NA |
| | 20 kb | 1469 | 69.8% | 32.8% | NA | 74.5% | 34 | 20.6% | 2.9% | NA | 20.6% |
| | 41 kb | 2167 | 68.7% | 26.2% | NA | 72.0% | 217 | 20.7% | 2.3% | NA | 21.2% |
| | 81 kb | 2417 | 71.0% | 24.0% | NA | 73.1% | 589 | 18.2% | 1.5% | NA | 18.8% |
| | 208 kb | 2228 | 76.1% | 24.7% | NA | 77.3% | 1111 | 14.5% | 1.0% | NA | 14.8% |
| | 425 kb | 1804 | 81.3% | 26.6% | NA | 82.0% | 1269 | 14.7% | 1.1% | NA | 14.9% |
| platelet | single-SNP | 53 | 69.8% | 41.5% | NA | 77.4% | 0 | NA | NA | NA | NA |
| | 3 kb | 245 | 62.4% | 29.4% | NA | 68.6% | 1 | 0.0% | 0.0% | NA | 0.0% |
| | 20 kb | 900 | 38.3% | 10.8% | NA | 40.8% | 102 | 6.9% | 0.0% | NA | 6.9% |
| | 41 kb | 1041 | 36.8% | 9.3% | NA | 38.5% | 220 | 3.6% | 0.0% | NA | 3.6% |
| | 81 kb | 1120 | 36.6% | 8.0% | NA | 37.5% | 450 | 3.6% | 0.2% | NA | 3.6% |
| | 208 kb | 1057 | 39.4% | 8.5% | NA | 40.1% | 686 | 4.2% | 0.6% | NA | 4.2% |
| | 425 kb | 911 | 42.9% | 9.0% | NA | 43.4% | 742 | 4.6% | 0.5% | NA | 4.6% |
| sbp | 3 kb | 83 | 83.1% | 12.0% | NA | 83.1% | 0 | NA | NA | NA | NA |
| | 20 kb | 177 | 89.3% | 9.6% | NA | 89.3% | 14 | 57.1% | 0.0% | NA | 57.1% |
| | 41 kb | 366 | 86.1% | 6.0% | NA | 86.3% | 145 | 29.7% | 0.0% | NA | 29.7% |
| | 81 kb | 496 | 86.5% | 4.4% | NA | 86.5% | 334 | 26.3% | 0.0% | NA | 26.3% |
| | 208 kb | 561 | 87.2% | 4.1% | NA | 87.2% | 622 | 24.8% | 0.0% | NA | 24.8% |
| | 425 kb | 538 | 90.0% | 4.3% | NA | 90.0% | 1005 | 22.4% | 0.0% | NA | 22.4% |

Table S9: Numbers of KnockoffGWAS discoveries containing previously reported associations. The results are stratified based on whether they are also detected by BOLT-LMM (as in Table 1). Other details are as in Table S8.

| Phenotype | Catalog | Japan | FinnGen |
|---|---|---|---|
| bmi | 4261 / 4514 (94.4%) | 5016 / 5094 (98.5%) | NA |
| cvd | 2223 / 4229 (52.6%) | NA | 2491 / 6713 (37.1%) |
| diabetes | 709 / 1906 (37.2%) | 5904 / 8550 (69.1%) | 93 / 577 (16.1%) |
| height | 4324 / 4461 (96.9%) | 61730 / 63254 (97.6%) | NA |
| hypothyroidism | 176 / 197 (89.3%) | NA | 89 / 462 (19.3%) |
| platelet | 1121 / 1159 (96.7%) | 7797 / 8012 (97.3%) | NA |
| respiratory | 1751 / 4112 (42.6%) | NA | 1129 / 9450 (11.9%) |
| sbp | 1781 / 2048 (87.0%) | 1757 / 1817 (96.7%) | NA |

Table S10: Total numbers of reported associations in the GWAS Catalog, Japan Biobank resource, or FinnGen resource, along with the corresponding fraction confirmed in our low-resolution analysis (425 kb). Other details are as in Table S8.

| Phenotype | Resolution | Total | | | Not found by BOLT-LMM | | |
|---|---|---|---|---|---|---|---|
| | | | Confirmed | | | Confirmed | |
| | | Discover. | Other | Other or Enrich. | Discover. | Other | Other or Enrich. |
| cvd | 3 kb | 22 | 22 (100.0%) | 22 (100.0%) | 0 | NA | NA |
| | 20 kb | 239 | 188 (78.7%) | 219 (91.6%) | 59 | 28 (47.5%) | 50 (84.7%) |
| | 41 kb | 339 | 266 (78.5%) | 309 (91.2%) | 127 | 72 (56.7%) | 107 (84.3%) |
| | 81 kb | 566 | 422 (74.6%) | 495 (87.5%) | 305 | 180 (59.0%) | 240 (78.7%) |
| | 208 kb | 940 | 738 (78.5%) | 764 (81.3%) | 666 | 472 (70.9%) | 493 (74.0%) |
| | 425 kb | 1089 | 967 (88.8%) | 968 (88.9%) | 834 | 714 (85.6%) | 715 (85.7%) |
| diabetes | 3 kb | 21 | 20 (95.2%) | 20 (95.2%) | 1 | 1 (100.0%) | NA |
| | 20 kb | 61 | 54 (88.5%) | 57 (93.4%) | 16 | 11 (68.8%) | 13 (81.2%) |
| | 41 kb | 109 | 88 (80.7%) | 97 (89.0%) | 55 | 36 (65.5%) | 42 (76.4%) |
| | 81 kb | 109 | 89 (81.7%) | 99 (90.8%) | 59 | 39 (66.1%) | 48 (81.4%) |
| | 208 kb | 113 | 97 (85.8%) | 106 (93.8%) | 61 | 46 (75.4%) | 54 (88.5%) |
| | 425 kb | 194 | 142 (73.2%) | 157 (80.9%) | 137 | 86 (62.8%) | 100 (73.0%) |
| hypothyroidism | single-SNP | 19 | 7 (36.8%) | 7 (36.8%) | 0 | NA | NA |
| | 3 kb | 40 | 24 (60.0%) | 24 (60.0%) | 0 | NA | NA |
| | 20 kb | 105 | 71 (67.6%) | 91 (86.7%) | 16 | 3 (18.8%) | 8 (50.0%) |
| | 41 kb | 222 | 105 (47.3%) | 172 (77.5%) | 94 | 21 (22.3%) | 61 (64.9%) |
| | 81 kb | 277 | 135 (48.7%) | 219 (79.1%) | 144 | 30 (20.8%) | 93 (64.6%) |
| | 208 kb | 295 | 156 (52.9%) | 226 (76.6%) | 166 | 43 (25.9%) | 101 (60.8%) |
| | 425 kb | 335 | 174 (51.9%) | 231 (69.0%) | 213 | 60 (28.2%) | 116 (54.5%) |
| bmi | 3 kb | 10 | 10 (100.0%) | 10 (100.0%) | 0 | NA | NA |
| | 20 kb | 343 | 308 (89.8%) | 328 (95.6%) | 34 | 16 (47.1%) | 29 (85.3%) |
| | 41 kb | 918 | 656 (71.5%) | 821 (89.4%) | 300 | 103 (34.3%) | 234 (78.0%) |
| | 81 kb | 1480 | 865 (58.4%) | 1182 (79.9%) | 688 | 190 (27.6%) | 450 (65.4%) |
| | 208 kb | 2395 | 1076 (44.9%) | 1620 (67.6%) | 1497 | 335 (22.4%) | 806 (53.8%) |
| | 425 kb | 2460 | 1091 (44.3%) | 1567 (63.7%) | 1666 | 409 (24.5%) | 820 (49.2%) |
| height | single-SNP | 95 | 81 (85.3%) | 81 (85.3%) | 0 | NA | NA |
| | 3 kb | 570 | 417 (73.2%) | 417 (73.2%) | 0 | NA | NA |
| | 20 kb | 1503 | 1102 (73.3%) | 1351 (89.9%) | 34 | 7 (20.6%) | 20 (58.8%) |
| | 41 kb | 2384 | 1607 (67.4%) | 1997 (83.8%) | 217 | 46 (21.2%) | 111 (51.2%) |
| | 81 kb | 3006 | 1879 (62.5%) | 2386 (79.4%) | 589 | 111 (18.8%) | 314 (53.3%) |
| | 208 kb | 3339 | 1886 (56.5%) | 2493 (74.7%) | 1111 | 164 (14.8%) | 556 (50.0%) |
| | 425 kb | 3073 | 1669 (54.3%) | 2231 (72.6%) | 1269 | 189 (14.9%) | 622 (49.0%) |
| platelet | single-SNP | 53 | 41 (77.4%) | 41 (77.4%) | 0 | NA | NA |
| | 3 kb | 246 | 168 (68.3%) | 230 (93.5%) | 1 | 0 (0.0%) | 0 (0.0%) |
| | 20 kb | 1002 | 374 (37.3%) | 778 (77.6%) | 102 | 7 (6.9%) | 49 (48.0%) |
| | 41 kb | 1261 | 409 (32.4%) | 934 (74.1%) | 220 | 8 (3.6%) | 127 (57.7%) |
| | 81 kb | 1570 | 436 (27.8%) | 1058 (67.4%) | 450 | 16 (3.6%) | 226 (50.2%) |
| | 208 kb | 1743 | 453 (26.0%) | 1017 (58.3%) | 686 | 29 (4.2%) | 256 (37.3%) |
| | 425 kb | 1653 | 429 (26.0%) | 922 (55.8%) | 742 | 34 (4.6%) | 297 (40.0%) |
| sbp | 3 kb | 83 | 69 (83.1%) | 69 (83.1%) | 0 | NA | NA |
| | 20 kb | 191 | 166 (86.9%) | 178 (93.2%) | 14 | 8 (57.1%) | 12 (85.7%) |
| | 41 kb | 511 | 359 (70.3%) | 441 (86.3%) | 145 | 43 (29.7%) | 97 (66.9%) |
| | 81 kb | 830 | 517 (62.3%) | 663 (79.9%) | 334 | 88 (26.3%) | 200 (59.9%) |
| | 208 kb | 1183 | 643 (54.4%) | 885 (74.8%) | 622 | 154 (24.8%) | 358 (57.6%) |
| | 425 kb | 1543 | 709 (45.9%) | 983 (63.7%) | 1005 | 225 (22.4%) | 474 (47.2%) |

Table S11: Numbers of KnockoffGWAS discoveries confirmed by other studies or enrichment analysis using independent GWAS summary statistics. Enrichment results are estimates. The results are stratified based on whether they are also detected by BOLT-LMM (as in Table S9).

| | | Total | | Not found by BOLT-LMM | |
|---|---|---|---|---|---|
| Phenotype | Resolution | Input | Confirmed | Input | Confirmed |
| cvd | 20 kb | 51 | 23–40 (45%–78%) | 31 | 15–27 (48%–87%) |
| | 41 kb | 73 | 33–53 (45%–73%) | 55 | 27–43 (49%–78%) |
| | 81 kb | 144 | 57–88 (40%–61%) | 125 | 45–74 (36%–59%) |
| | 208 kb | 202 | 8–47 (4%–23%) | 194 | 5–40 (3%–21%) |
| | 425 kb | 122 | 0–7 (0%–6%) | 120 | 0–5 (0%–4%) |
| diabetes | 3 kb | 1 | 1–1 (100%–100%) | 0 | NA |
| | 20 kb | 7 | 0–5 (0%–71%) | 5 | 0–5 (0%–100%) |
| | 41 kb | 21 | 3–14 (14%–67%) | 18 | 1–11 (6%–61%) |
| | 81 kb | 20 | 6–15 (30%–75%) | 19 | 5–14 (26%–74%) |
| | 208 kb | 16 | 4–14 (25%–88%) | 15 | 3–12 (20%–80%) |
| | 425 kb | 52 | 6–27 (12%–52%) | 51 | 4–25 (8%–49%) |
| hypothyroidism | single-SNP | 12 | 12–12 (100%–100%) | 0 | NA |
| | 3 kb | 16 | 11–16 (69%–100%) | 0 | NA |
| | 20 kb | 34 | 13–26 (38%–76%) | 13 | 2–9 (15%–69%) |
| | 41 kb | 117 | 53–81 (45%–69%) | 73 | 30–51 (41%–70%) |
| | 81 kb | 142 | 69–98 (49%–69%) | 114 | 49–76 (43%–67%) |
| | 208 kb | 139 | 55–86 (40%–62%) | 123 | 43–73 (35%–59%) |
| | 425 kb | 161 | 41–74 (25%–46%) | 153 | 40–73 (26%–48%) |
| bmi | 20 kb | 35 | 13–27 (37%–77%) | 18 | 8–18 (44%–100%) |
| | 41 kb | 262 | 146–184 (56%–70%) | 197 | 115–147 (58%–75%) |
| | 81 kb | 615 | 284–350 (46%–57%) | 498 | 231–289 (46%–58%) |
| | 208 kb | 1319 | 494–595 (37%–45%) | 1162 | 422–518 (36%–45%) |
| | 425 kb | 1369 | 424–529 (31%–39%) | 1257 | 361–461 (29%–37%) |
| height | single-SNP | 14 | 0–9 (0%–64%) | 0 | NA |
| | 3 kb | 153 | 90–123 (59%–80%) | 0 | NA |
| | 20 kb | 401 | 225–272 (56%–68%) | 27 | 7–19 (26%–70%) |
| | 41 kb | 777 | 353–426 (45%–55%) | 171 | 47–84 (27%–49%) |
| | 81 kb | 1127 | 460–552 (41%–49%) | 478 | 174–234 (36%–49%) |
| | 208 kb | 1453 | 555–660 (38%–45%) | 947 | 349–434 (37%–46%) |
| | 425 kb | 1404 | 509–615 (36%–44%) | 1080 | 387–478 (36%–44%) |
| platelet | single-SNP | 12 | 3–12 (25%–100%) | 0 | NA |
| | 3 kb | 78 | 53–70 (68%–90%) | 1 | 0–0 (0%–0%) |
| | 20 kb | 628 | 373–433 (59%–69%) | 95 | 29–55 (31%–58%) |
| | 41 kb | 852 | 488–561 (57%–66%) | 212 | 100–138 (47%–65%) |
| | 81 kb | 1134 | 578–665 (51%–59%) | 434 | 181–238 (42%–55%) |
| | 208 kb | 1290 | 514–614 (40%–48%) | 657 | 190–264 (29%–40%) |
| | 425 kb | 1224 | 442–542 (36%–44%) | 708 | 224–301 (32%–43%) |
| sbp | 3 kb | 14 | 3–12 (21%–86%) | 0 | NA |
| | 20 kb | 25 | 5–18 (20%–72%) | 6 | 2–6 (33%–100%) |
| | 41 kb | 152 | 67–97 (44%–64%) | 102 | 40–67 (39%–66%) |
| | 81 kb | 313 | 122–169 (39%–54%) | 246 | 90–133 (37%–54%) |
| | 208 kb | 540 | 209–273 (39%–51%) | 468 | 173–233 (37%–50%) |
| | 425 kb | 834 | 232–316 (28%–38%) | 780 | 209–289 (27%–37%) |

Table S12: Bootstrap confidence intervals (90%) for the proportion of novel discoveries confirmed by the enrichment analysis in Table S11.

| Phenotype | Discoveries | Contains gene | Known lead SNP consequence | Known lead SNP association |
|---|---|---|---|---|
| cvd | 31 | 26 (84%) | 28 (90%) | 21 (68%) |
| diabetes | 5 | 5 (100%) | 3 (60%) | 5 (100%) |
| hypothyroidism | 13 | 12 (92%) | 8 (62%) | 9 (69%) |
| respiratory | 6 | 5 (83%) | 4 (67%) | 3 (50%) |

Table S13: Numbers of novel discoveries (not found by BOLT-LMM and not confirmed by the other studies in Table S9) that either contain a gene or whose lead SNP has a known functional annotation or a known association with phenotypes closely related to that of interest.

| Phenotype | Associations |
|---|---|
| cvd | NA (10), blood pressure (9), BMI (8), obesity (3), cardiovascular disease (1), CCL2 (1), cholesterol (1), triglycerides (1), heart rate (1) |
| diabetes | diabetes (3), Factor VII (1), glyburide metabolism (1) |
| hypothyroidism | NA (4), autoimmune thyroid disease (2), psoriasis (2), diabetic nephropathy (1), Graves disease (1), hypothyroidism (1), rheumatoid arthritis (1), thyroid function (1) |
| respiratory | NA (3), hypersomnia (1), interaction with air pollution (1), serum IgE (1) |

Table S14: Associations of our novel discoveries (20 kb resolution) in Table S13 to related traits. The same discovery may have more than one relevant association in this table.

| Consequence | cvd | diabetes | hypothyroidism | respiratory |
|---|---|---|---|---|
| 2KB Upstream | | 1 | | |
| 3 Prime UTR | 2 | | | |
| 500B Downstream | | | | 1 |
| Intron | 19 | 2 | 6 | 3 |
| Missense | 3 | | 1 | |
| Non coding transcript exon | 1 | | | |
| Regulatory region | 2 | | | |
| Stop gained | | | 1 | |
| Tf binding site | 1 | | | |
| Unknown | 3 | 2 | 5 | 2 |
| Total | 31 | 5 | 13 | 6 |

Table S15: Numbers of lead variants with known consequences for our novel discoveries (20 kb resolution) in Table S13.

[14] B. Klaus, K. Strimmer, and M. K. Strimmer. “Package ‘fdrtool‘”. In: ().
